# EnCPdock: a web-interface for direct conjoint comparative analyses of complementarity and binding energetics in inter-protein associations

**DOI:** 10.1101/2023.02.26.530084

**Authors:** Gargi Biswas, Debasish Mukherjee, Nalok Dutta, Prithwi Ghosh, Sankar Basu

## Abstract

**Context:** Protein-protein interaction (PPI) is a key component linked to virtually all cellular processes. Be it an enzyme catalysis (‘*classic type functions*’ of proteins) or a signal transduction (‘*non-classic’*), proteins generally function involving stable or quasi-stable multi-protein associations. The physical basis for such associations is inherent in the combined effect of shape and electrostatic complementarities (Sc, EC) of the interacting protein partners at their interface, which provides indirect probabilistic estimates of the stability and affinity of the interaction. While Sc is a necessary criterion for inter-protein associations, EC can be favorable as well as disfavored (e.g., in transient interactions). Estimating equilibrium thermodynamic parameters (ΔG_binding_, K_d_) by experimental means is costly and time consuming, thereby opening windows for computational structural interventions. Attempts to empirically probe ΔG_binding_ from coarse-grain structural descriptors (primarily, surface area based terms) have lately been overtaken by physics-based, knowledge-based and their hybrid approaches (MM/PBSA, FoldX etc.) that directly compute ΔG_binding_ without involving intermediate structural descriptors.

**Methods:** Here we present EnCPdock (www.scinetmol.in/EnCPdock/), a user-friendly web-interface for the direct conjoint comparative analyses of complementarity and binding energetics in proteins. EnCPdock returns an AI-predicted ΔG_binding_ computed by combining complementarity (Sc, EC) and other high-level structural descriptors (input feature vectors), and, renders a prediction accuracy comparable to the state-of-the-art. EnCPdock further locates a PPI complex in terms of its {Sc, EC} values (taken as an ordered pair) in the two-dimensional Complementarity Plot (CP). In addition, it also generates mobile molecular graphics of the interfacial atomic contact network for further analyses. EnCPdock also furnishes individual feature trends along with the relative probability estimates (Pr_fmax_) of the obtained feature-scores with respect to the events of their highest observed frequencies. Together, these functionalities are of real practical use for structural tinkering and intervention as might be relevant in the design of targeted protein-interfaces. Combining all its features and applications, EnCPdock presents a unique online tool that should be beneficial to structural biologists and researchers across related fraternities.

## 1. Introduction

A host of biological processes thrives on the efficacy of protein-protein interactions (PPI) ranging from agglutination reactions to aggregation-based processes initiating the immune response cascade [1]. It would be of fundamental biophysical value to be able to find the determining premise between the dynamic interplay of different forces sustaining proteins in their bound forms and their differential affinities – recorded by skill-intensive, time-consuming and expensive experimental techniques. These experimental studies (so called, solution assays) are mostly *in vitro,* offering only a reduced representation of the system (dilute solutions) [2], unable to mimic the real biological context of the complex cellular milieu; whereas, *in vivo* crowding studies are few and far, and, even in them, cellular complexity can not be attained [2]. Some of these experimental techniques (e.g., fluorescence spectroscopy, surface plasmon resonance etc.) are again indirect [3–5], and, even direct calorimetric methods [6] are known to have their own limits of accuracy.

The wealth of high resolution experimental structural data harbored for these PPI complexes in the PDB [7] offers the basis of structure based thermodynamics – one unmet goal of which is to be able to predict affinities (in terms of either K_d_ ^1^ or ΔGG ^2^) of inter-protein associations with enough accuracy [8], directly from 3D atomic coordinates. In the course, the complex interplay of physicochemical principles (covering different local and non-local forces) underlying inter-protein association is also expected to be (at least, partially) unraveled and expressed in terms of bio-energetic estimates (T_m_, ΔGH, TΔGS, ΔGG etc.). Besides, one direct application of binding affinity determination is, of course, to serve the design and discovery of novel therapeutics and to aid mutagenesis studies. Combining kernels involving multiple layers with different regressor models such as Support Vector Machines (SVM), Ordinary Least-Squares Regressor (OLSR) and Random Forest Regressor (RFR) have been able to even extend the prediction of protein affinity directly from primary sequences of the binding partners [9] in absence of known 3D structures.

Computational prediction of protein binding affinities dates back to the days of accessible surface area measures (e.g., ΔGASA) [10] only attaining a modest correlation of r=–0.16 [11] with experimental binding free energy (ΔGG_exp_) even for the rigid complexes. Subsequent performances of different approaches, however have followed an ascending trend over time, till the current era of the sophisticated ‘affinity scores’ [11]. These mostly AI-trained scores are certainly preferred among affinity predictors due to their implicit ability to meticulously absorb any factor affecting PPIs (fed in as input feature vectors) and the flexibility of using empirical data instead of a fixed or predetermined function. The highest reported correlation (to the best of our knowledge) among these affinity scores (with ΔG_exp_) has been r=−0.73 [8], recorded on a standard protein affinity benchmark [12], by a linear regression model trained on ‘inter-residue contact (IC)’ descriptors of affinity, categorized based on their residue hydrophobicities [13].

Among non-AI approaches, there are semi-empirical, force-field based and hybrid energy functions which can be broadly classified into Physical (PEEF), Statistical (SEEF) [14] and Empirical (EEEF) [15] Effective Energy Functions. While PEEF and SEEF respectively are purely physics- (or, force field) and purely knowledge-based approaches (analogous to two independent orthogonal vectors in a vector-space), EEEF (much like their resultant) provides a mean to combine them with empirical energy terms scaled to better fit with experimental stability estimates [16]. For example, one of the most popular modern-day energy functions, FOLDEF (parameterized on a large 1000-mutation database) [17] is empirical in nature and was initially procured from a ‘folding pathway predictor’ in proteins (FOLD-X) [18] that implemented a combination of PEEF and SEEF terms; and, critically pointed out that the key to the improved prediction required elimination of side-chains with high B-factors and explicit consideration of water molecules in the protein cavities [15]. Among other approaches, are MM/PBSA^3^ and/or MM/GBSA^4^ methods (widely employed in protein-protein/drug interactions [19]) which are based on integrating energy contributions originated from different types of interactions and molecular groups, and, have found better feasibility than energy perturbation methods for their lowered costs [20], and, also perhaps for their specialized ability to identify near-native binding modes in PPIs [21].

Non-covalent interactions persisting within proteins (common in context to binding and folding [22]) can broadly be classified into short range Van der Waals forces (packing) and long range electrostatic forces (Hydrogen bonds, salt-bridges, charge-dipoles etc.), together, implicitly coupled with solvent effects (hydrophobic interactions and entropic costs). Complementarity is one concept that have been able to bridge the gap between binding and folding in proteins [23] taking account of all aforesaid features under one big umbrella. Categorically speaking, the very physical basis for macro-molecular (e.g., protein – protein) binding is inherently implicit in the dual nature of complementarity in shape and surface electrostatic potentials {Sc, EC} of the interacting macro-molecular (e.g., protein) surfaces [24]. Together, as part of the Complementarity Plot (CP) for docking (CPdock) [25], they provide indirect probabilistic estimates of the stability and affinity of the interaction [26]. Naturally, they are chosen as the first two input feature vectors in EnCPdock. Both, Sc and EC are surface-derived measures [27,28] and offer high-level of sophistication in the structural description of the (target) complexes. Moreover, the choice of both Sc, EC as input feature vectors accounts for both local (Sc) and non-local (EC) nature of complementarity of the interacting protein partners (at their interface). The shape of both the molecular partners should be such that the side-chain atoms be engaged in complementary interlocking at the interface for a feasible interaction [23,27]. Thus, an elevated Sc threshold (Sc>0.55, non-rigid [26]) serve as a necessary criteria for inter-protein [27,29–31] binding with extended surface overlaps [1]. This has merited the common use of shape complementarity measures as primary screening filters in many docking scoring algorithms/pipelines [6,21]. In parallel, favorable electrostatic interactions at the interfacial protein surfaces appreciably stabilizes them in their bound form [28]. Hence the desired anti-correlation of surface electrostatic potentials (hallmark of elevated EC values) between interacting (protein) surfaces should appreciably stabilize the complex [33,34]. Over the years, the use of EC [28] has thus also found its scope in several computational endeavors, e.g., in docking [24,25], homology modeling, structure validation etc [35]. However, sub-optimal, even negative EC values (i.e., unfavorable electrostatics) between the interacting protein surfaces have been recorded in ~20% of all binary PPI complexes [32], often compensated by a really elevated geometric fit between the interacting surfaces. These unfavorable electrostatic interactions often map to transient interactions [26,36], serving as molecular / conformational switches. Consequently, EC serves as a secondary or sufficient criteria for protein-protein interactions [26].

The current paper presents a unique user-friendly web-interface, namely, EnCPdock (www.scinetmol.in/EnCPdock/) that allows direct conjoint comparative analyses of complementarity as well as binding free energetics of PPI complexes. An AI-predicted ΔGG_binding_ is returned by EnCPdock, determined by combining complementarity (Sc, EC), surface area estimates and other high-level structural descriptors taken as input feature vectors. In addition, EnCPdock returns the usual mapping of the binary PPI complex in the Complementarity Plot (as in CPdock [25]). The detailed applications of the Complementarity Plot (with different built to meet different purposes: either {Sc, EC}: overall [25] or {S_m_, E_m_}: residue-wise [23,35]) have been demonstrated across several publications in the past [23,25,26,35–37] spanning from homology modeling [35], docking scoring and optimization [36,38] to protein, epitope and interface design [26,37,39]. In context to the current application, EnCPdock, indirect probabilistic stability and affinity estimates of an input (docked / bound) PPI complex are first procured by CPdock in terms of its {Sc, EC} values (taken as an ordered pair), mapped onto one of the three probabilistic regions (‘*probable*’, ‘*less probable*’, ‘*improbable*’ – analogous to ‘*allowed*’, ‘*partially allowed*’, ‘*disallowed*’ regions of the legendary Ramachandran Plot [40,41]). Over and above, the current web-interface (EnCPdock) goes deeper into protein binding energetics and predicts the actual free energy of interaction (ΔGG_binding_) using a non-linear support vector regression (SVM) machine trained on a collection of wisely chosen high-level structural descriptors. As a free energy predictor, EnCPdock performs comparable to the state-of-the-art on standard experimental benchmarks – as revealed by multiple independent validations.

In addition, EnCPdock also generates mobile molecular graphics of the interfacial atomic contact network and returns the contact map for further analyses. Together, this provides a direct visual and analytical platform to identify specific native interactions (contacts) contributing to the binding and their persistence or transience in a library of mutants. EnCPdock also furnishes individual feature trends along with the relative probability estimates (Pr_fmax_) of the obtained feature-scores with respect to the events of their highest observed frequencies – together, which presents a handy, first-hand tool for the targeted design of protein interfaces (or, dockable peptides), helping investigators to pinpoint structural defects, irregularity and sub-optimality – which would, in turn, aid in the subsequent re-design. Combining all its features and applications, EnCPdock presents a unique online tool that should be beneficial to structural biologists and researchers across related fraternities.

## 2. Materials and Methods

### 2.1. Datasets

The primary database was built from scratch for training and cross-validation of EnCPdock. To that end, first, protein binary complexes were collected from the Protein Data Bank (PDB) using its advanced search options with the following culling criteria:

1. Each target molecule must be a protein-protein dimer complex (homo- and hetero-dimers) and not a single protein.
2. All atomic models must be determined either by X-ray crystallography or by cryo Electron Microscopy.
3. All structures had to have a resolution of ≤ 2Å.
4. None of the structures had RNA/DNA chains.
5. For X-ray structures, they had to have a R-observed ≤ 0.2.
6. None of the structures contained large prosthetic groups or co-factors (i.e., a non-protein chemical component with MW > 200 Da), e.g., proto-porphyrin rings, NADP, acetyl CoA, etc.
7. Each complex should have had at least one residue coming from each partner at their interface buried upon association.

This led to 6658 structures (asymmetric units) which got further reduced to 3200 non-redundant binary PPI complexes upon implementation of a sequence identity cutoff (in NW-align: https://zhanggroup.org/NW-align/) of no more than 30% between any two receptor chains and separately for any two ligand chains. The PDB IDs along with the corresponding chain IDs of receptors and ligands of PPI complexes pertaining to the final training dataset are to be found enlisted in **Dataset S1, Supplementary Materials**.

For independent validation purposes, firstly, all binary PPI complexes (entirely non-overlapping to the training dataset) were culled from the updated version of the Affinity benchmark (v.2) [11]. All these structures had annotated experimental binding affinities and/or free energies (K_d_ or ΔG_binding_) associated with each entry. Structures with reported suboptimal accuracy in terms of binding energetics (https://github.com/haddocking/binding-affinity-benchmark) were further discarded. This led to a final size of 106 binary PPI complexes (targets) (**Dataset S2**, **Supplementary Materials**) for the 1^st^ dataset used in independent validation. A second experimental benchmark for independent validation was assembled by merging the available wild-type thermodynamic data retrieved from Proximate (https://www.iitm.ac.in/bioinfo/cgi-bin/PROXiMATE/wild.py) [42] and SKEMPI 2.0 (https://life.bsc.es/pid/skempi2/) [43] datasets, which together gave us (a second set of) 236 binary PPI complexes (targets) (**Dataset S3**, **Supplementary Materials**).

### 2.2. Built of EnCPdock

#### 2.2.1. The input feature vectors

Six input feature vectors that are effectively high-level (fine-grained) structural descriptors of the overall protein complex or the protein-protein interface have been utilized for training of EnCPdock. The thirteen structural features used to build EnCPdock can be broadly classified into four groups: (i) complementarity descriptors (Sc, EC) (ii) accessibility descriptors (nBSA, nBSA_p_, nBSA_np_, fracI) (iii) interfacial contact network descriptors (Ld, ACI, slope_dd_, Yinter_dd_, CCp_dd_) and (iv) size descriptors (logN, log_asp_); and, are defined as follows.

##### 2.2.1.1. Surface or shape complementarity (Sc)

Qualitatively, the shape complementarity (Sc) function measures the goodness of fit between two surfaces, and in case of PPI complexes this function serves as an essential condition for binding. For calculation of Sc, van der Waal’s surfaces of two interacting partners (e.g., protein-protein, protein-DNA or residue-residue) is generated, and for each pair of points from the respective surfaces, the distance between points and the dot product between the corresponding normals are taken into account [31]. The elaborate mathematical description of shape complementarity function can be obtained from the work of Lawrence and Colman [27] whose value ranges from −1 to +1 for an anti- and perfectly correlated surfaces respectively. The Sc value of a protein-protein interface was computed by the original program ‘sc’ (© Lawrence), distributed as a part of the CCP4 package [44].

##### 2.2.1.2. Electrostatic complementarity (EC)

The complementarity of charge distribution upon two interacting surfaces is measured by electrostatic complementarity (EC) function. Similar to Sc, EC is also a necessary determinant for binding. For the calculation of EC, the molecular surface is generated first using the software EDTSurf [45], and then using the DelPhi software [46] the electrostatic potential on each surface point at the protein-protein interface was determined. The atoms who have undergone a change (non-zero) in the solvent accessible surface area (ASA) upon binding, have been identified as the atoms at the protein-protein interface (the difference in ASA is calculated using NACCESS with a probe size (radius) trivially set to 1.4 Å, the hydrodynamic radius of water [47]. For the calculation of electrostatic potential, the partial charges and atomic radii were assigned from AMBER94 molecular mechanics force field [48], and the solution was obtained by iteratively solving a linearized version of Poission-Boltzman equation (employing DelPhi). Similar to Sc, EC value also ranges from −1 to +1, where approaching the value 1 signifies the anti-correlation of electrostatic potentials between two interacting surfaces, and negative EC implies unbalanced electric fields at the interface.

##### 2.2.1.3. Normalized buried surface area (nBSA)

Although Sc and EC are essential measures for binding, but they do not consider the area or size of the interface of PPI complexes. So, in order to incorporate that feature, two additional measures have been taken into account. For any atom (say, i^th^ atom) at the interface, for which there is a non-zero change in the ASA (as mentioned previously), the buried surface area has been calculated using the following formula,

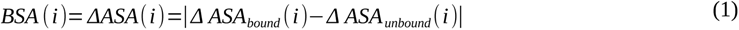

- where *ΔASA_bound_* stands for the accessible surface area (ASA) at the bound state and *ΔASA_unbound_* is the ASA at the unbound state. Now, for the calculation of nBSA, for a protein complex consisting of two proteins (say, protein A and protein B) the following formula have been used,

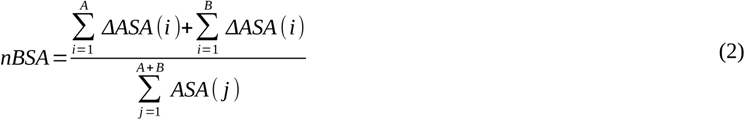

- where 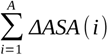 and 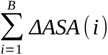 are sum over the *ΔASA* of all the interface atoms of protein A and protein B, and 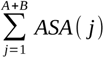 is the total ASA of all atoms of protein A and protein B. For calculation of nBSA, all the interface atoms, irrespective of their polarity have been taken into account.

Further, considering the polarity of the interface atoms, nBSA has been calculated separately for polar (N and O) and non-polar (C and H) atoms [10] and the corresponding nBSA have been called as nBSA_p_ and nBSA_np_ respectively.

##### 2.2.1.4. Fractional Interfacial content (fracI)

Similar to nBSA, the fraction of residues buried at the interface of a PPI complex (consisting of two proteins, say protein A and protein B) have been defined by the following formula,

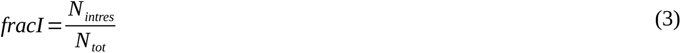

- where N_intres_ and N_tot_ are respectively the number of interfacial residues and total number of residues pertaining to the protein-protein complex.

In addition to the complementarity and accessibility based structural descriptors, several precise network parameters have also been included as input feature vectors. Given these are essentially binary (receptor – ligand) protein-protein interactions, the resultant PPI networks at the interconnected interface would necessarily map to bipartite graphs. Importantly, two residues at the interfaces from each of the partner molecules (receptor and ligand) are connected with a link if at least one non hydrogen atom from both the residues are within 4 Å from each other. Notably, the distance cutoff for C-C van der Waal’s contact is 3.8 Å, thus, threshold for formation of a link (i.e., 4 Å) applied here is reasonably stringent [49,50]. However, after defining the criteria for formation of an inter-residue inter-chain link, subsequent construction of bipartite network adjacency matrices was derived. In connection with these, a few network-based features were designed as described as below.

##### 2.2.1.5. Link density (Ld)

Let the receptor and the ligand have N1 and N2 interfacial residues in physical contact with one or more residues coming from the partner molecules for each. The link density for such a bipartite network could then be defined as follows,

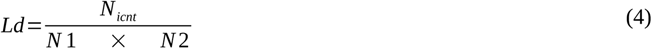

- where N_icnt_ is the number of inter-chain inter-residue contacts to have formed at the said receptor – ligand interface. In short, Ld can be defined as the ratio of the actual number of links to the theoretical maximum number of links between two chains and, it’s value will range from 0 to 1, where a complete bipartite graph (i.e., every node of the receptor is connected to every node of ligand and vice-versa) would hit a value of 1.

##### 2.2.1.6. Average contact intensity (ACI)

As defined in an earlier work from this laboratory in context to salt-bridges [50], ACI accounts for the individual intensities or atomic contact densities of each inter-residue interchain link to have formed in a PPI bipartite network. To that end, the actual number of inter-atomic contacts [atcon(i)] involved in each (i^th^) inter-residue interchain link was averaged over all links [36].

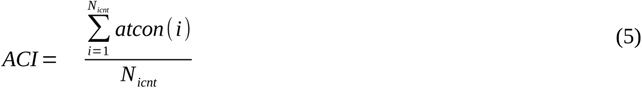

##### 2.2.1.7. Degree distribution profile-based features

Degrees of each node constituting these PPI bipartite networks were computed from their corresponding (unweighted) 1-0 adjacency matrices and their normalized frequency distributions were plotted in a log_10_-scale for both axes. Given their characteristic power law distributions, the obtained points in these log-log plots for each graph would follow linear relationships. To that end, a linear least-squares fitting was performed on the obtained points (for each graph) from which the slope (slope_dd_), the Y-intercept (Yinter_dd_) and the Pearson’s correlation coefficient (r) value between the observed (Y_obs_) and the expected abscissa (as in, Y_exp_ = slope_dd_.Xobs + Yinter_dd_) coming from the plotted points were analytically determined. The r value (let’s call it CCp_dd_) between Y_exp_ and Y_obs_ [r (Y_exp_,Y_obs_)] were determined using the well-known relation [51],

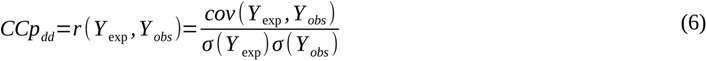

- where *cov* is the co-variance between the mentioned parameters within the bracket and σ is the variance of each parameter.

##### 2.2.1.8. Features based on comparative and absolute size of the interaction partners

The chain length of each interacting molecular partner (lenR: receptor, lenL: ligand) was taken as the estimate of their size. Two features were then defined based on these chain lengths as follows,

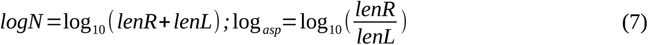

The second of these terms can be considered analogous to the aspect ratio (in log scale) of a crystal or a nano particle. For identical sizes of receptor and ligand (e.g., homo-dimeric biological assemblies) log_asp_ would hit a value of 0. The primary purpose of taking logarithm for these (comparative and absolute) chain length features was to render values in a desired (low numeric) range comparable to that of the other SVM input feature vectors.

#### 2.2.2. Training & cross-validations

A Support Vector Regression module (as in SVM^light^ [52]) have been used to train EnCPdock in a 10-fold cross validation scheme with a radial basis function (RBF) kernel – the details of which are as follows in the following sections.

##### 2.2.2.1. Cross validation test-set and parameter optimization

All the 3200 PPI complexes were divided into ten equal parts, nine of which were utilized for each round of training, while the remaining ones were used as the test set in that round. This was performed for each of the ten parts to get predictions for the whole set. Three parameters, namely, C, γ and ɛ have been optimized using a grid search method to obtain a maximum correlation (determined by Pearson’s correlation coefficient, r) between target function and the predicted output. Among the three parameters, C is the penalty value associated to the training error (for large and small value of C, a smaller and larger margin were accepted respectively if all the training points were correctly predicted); RBF γ-value defines how far the influence of a single training example reaches (low value indicates far and high value indicates close); and ɛ is the distance of the target function from the actual value within which no penalty is associated in the training loss function [53]. Optimization of C, γ and ɛ was performed within the ranges 1.0 to 5.0 (in steps of 0.5), 0.01 to 2.0 (in steps of 0.01) and 0.1 to 1.0 (in steps of 0.05) respectively.

##### 2.2.2.2. Target function and predicted output

During both the training and cross-validation procedure, ΔGG_binding_ was predicted by FoldX [54] which is a well-accomplished empirical force-field based method parameterized by empirical data from actual protein engineering experiments [55] and is known to return near-native binding/affinity estimates. The FoldX platform is based on a ‘fragment-based strategy’ which uses fragment libraries in an analogous way to the most fascinating ‘fragment assembly simulated annealing’ approach for protein structure prediction credited to David Baker and Rosetta [56]. The standalone (C++ with boost library) version (v.4) of FoldX (http://foldxsuite.crg.eu/) was used. The FoldX – derived ΔGG_binding_ (let’s call it ΔGG_FoldX_) was normalized by the number of interfacial residues, N_intres_ (let’s call it ΔGG_FoldX_norm_ denoting average interfacial contribution to binding free energy) for all the PPI complexes. This ΔGG_FoldX_norm_ was suitable numerically to be used as the target function in the training of the free-energy predictor (in the EnCPdock web-interface) which naturally returned the predicted output interpretable in terms of the average interfacial contribution (i.e., normalized) to ΔGG_binding_ (let’s call it, ΔGG_EnCPdock_norm_). The normalization (of the target function) also ensured similar numerical ranges for both the target function (TF) and the predicted output (PO). In the final output of the EnCPdock web-interface, both average interfacial (i.e., normalized) and total binding free energies were returned.

##### 2.2.2.3. Support Vector Regression (SVM) training & parametrization

To train EnCPdock, a Support Vector (regression) Machine (as in SVM^light^ [52]) have been implemented using a 10-fold cross validation scheme with a Radial Basis Function (RBF) kernel. The popularity and efficiency of SVM methods in pattern recognition has been evident ever since its introduction [57]. Among different SVM models, again, RBF kernels serve as optimal predictors when implemented with appropriate regularization, minimizing estimation and approximation errors [58]. SVM is built with the rare abilities (for example) to (i) produce an unique solution, (ii) remain robust to outliers, (iii) easily update the decision model, and, (iv) allow combination of classifiers trained on different data types by applying probability rules [57]. SVM is easy to implement (by virtue of standalone packages like SVM^light^ [52]) and less computationally demanding [57,59]. Motivated by these factors as well as the successful combined use of the SVM-RBF combo across many problems in computational biology [9,32,60–62] inclusive of closely related ones like protein docking scoring and sequence based affinity prediction [9,32], the ‘SVM-RBF’ combo offered an automatic first choice in EnCPdock. The RBF kernel ensured a non-linear relationship between the input feature vectors and the target function (ΔGG_EnCPdock_norm_). In a SVM classification, for a given set of n observations given by (x1, y1), …, (xn, yn), predictions are made from the following formula [63].

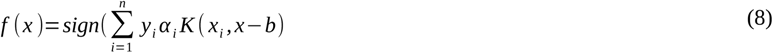

- where b is a numeric offset threshold and α_i_ denote the weight of each observation, known as Kühn-Tucker coefficient. The Kernel function K (x_i_, x) defines a dot product to transform the observations from input space to the feature space. Here, we have used a Gaussian Kernel function, popularly known as an RBF Kernel, given by,

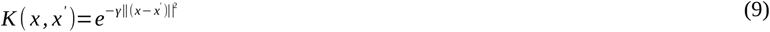

- where γ is a parameter (>0) controlling the width of the Gaussian Kernel. γ is a free parameter whose value can be optimized. Apart from γ, two other free parameters have been optimized during the SVM training, namely, C and ε. C is a cost parameter which controls the trade-off between minimizing the number of misclassified samples in the training set and maximizing the margin width; while ε can be defined as the sum of classification error and the complexity penalty given by,

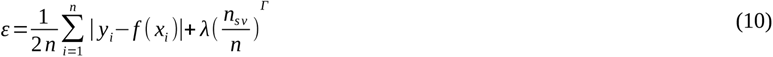

- where the first term is the classification error and, n_sv_, λ and Γ are number of input feature vectors, regularization parameter and the penalty respectively. During the SVM training and cross validation, these three parameters c, γ and ε have been optimized using a grid search method to obtain a maximum correlation (determined by the Pearson’s correlation coefficient, r, eqn. 11) between target function and the predicted output. As detailed earlier (see section 2.2.2.2, **Materials and Methods**), for each individual training cycle, the target function was taken to be ΔG_FoldX_norm_ while the predicted output was the ΔG_EnCPdock_norm_ for the test set predicted by the rest of the nine subsets. Thus, for a well-trained predictor, a high correlation should be expected in the performed cross-validation between the predicted values of ΔG_binding_norm_ by EnCPdock with reference to FoldX predictions in the same test set. However, a more authentic indicator of the same would be the recovery of performance of the trained predictor in an independent validation performed on a separate dataset of PPI complexes (completely non-overlapping to that of the training set) with their annotated experimental binding free energies taken as references.

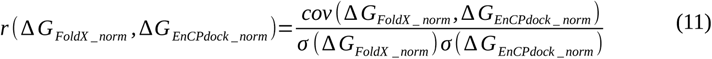

To obtain a maximum r value during the training the c, γ and ε were optimized within the ranges 1.0 to 5.0 (in steps of 0.5), 0.01 to 2.0 (in steps of 0.01) and 0.1 to 1.0 (in steps of 0.05) respectively.

#### 2.2.3. Evaluation metrics

To evaluate the performance of an SVM model on test predictions, firstly, the number of true positives (TP), true negatives (TN), false positives (FP) and false negatives (FN) were quantified. Based on these measures, the performance of the predictor (EnCPdock) was estimated by the following evaluation metrices.

First, the actual positives and actual negatives that were correctly identified out of all predicted positives and predicted negatives respectively were determined by the true positive rate (TPR) and false positive rate (FPR), as defined below.

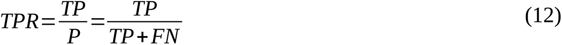

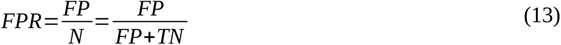

- where P is the total number of positives, consisting of true positives and false negatives while N is the total number of negatives consisting of false positives and true negatives. The positive and negative cases were identified based on various cutoffs spanning the entire range of the target function (i.e., ΔGG_FoldX_norm_). Further, the receiver operating characteristics curves (ROC) were plotted (TPR vs. FPR) and the Area Under the Curve (ROC-AUC) were computed. For a random predictor, the AUC will exhibit a value of 0.5 and a perfect prediction model would hit a value of 1.0 [64]. In parallel, the balanced accuracy (BACC) score was computed, defined as follows.

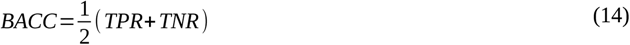

- where TPR and TNR are the true positive rate and true negative rates respectively. TNR can be defined similar to TPR by the following formula,

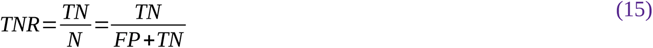

To note, since BACC score is the arithmetic mean of TPR and TNR, swapping the definition of TPR and TNR does not alter the BACC score.

## 3. Results and Discussion

The objective of the current study was to develop an integrated web-interface (EnCPdock) to analyze complementarity and energetics of protein – protein interactions, serving as a common conceptual platform, that could be of practical use to researchers across the modern biology fraternity. The potential beneficiaries include experimental as well as computational structural biologists, and, also other members of the broad biological community working with specific protein – protein interactions involved in enzyme catalysis, signal transductions and/or other protein functionalities. The energetics part of EnCPdock has been based on supervised learning, hitting comparable (if not better) accuracy to predict binding free-energies (ΔG_binding_) of PPI complexes from the knowledge of fine-grained high-level structural descriptors. PPI complexes are probed by EnCPdock in terms of its {Sc, EC} values (taken as an ordered pair) in the two-dimensional Complementarity Plot (as in CPdock [25]) evaluating indirect probabilistic estimates of stability. The uniqueness of our approach lies in the operational credibility of the EnCPdock system, which is a comprehensible web-interface (www.scinetmol.in/EnCPdock/) that permits direct, synchronized comparisons of complementarity and binding energies of PPI complexes. The performance of EnCPdock has been evaluated and authenticated in a two-step validation pipeline: (i) firstly, by a 10-fold cross validation performed on a large non-redundant training set consisting of binary PPI complexes with structural coordinates alone (irrespective of having experimental references to their ΔG_binding_ values) and (ii) backed up by an independent validation performed on one of the latest non-redundant datasets of binary PPI complexes [11] with recorded experimental ΔG_binding_ values. The independent validation clearly shows recovery of performance with respect to the cross-validated results – which is comparable (if not better) to the state-of-the-art. In addition, EnCPdock not only returns the mapping of the binary PPI complex in the Complementarity Plot but also further returns mobile molecular graphics of the atomic contact network probed at the protein-protein interface, along with the detailed lists of weighted inter-residue contacts for further analyses for interested parties.

### 3.1. SVM Training and Cross-validation

As described in the **Materials and Methods** (section 2.2.1), thirteen high-level structural descriptors were taken as input feature vectors to build and train the binding free-energy predictor in EnCPdock. These features together may be considered to be harboring necessary and sufficient fine-grained structural details of the PPI complexes. Due to the scarcity of sufficient high quality experimental binding or affinity data, the training of EnCPdock had to be performed on FoldX – derived binding free energies (ΔG_binding_) [65], appropriately normalized to ΔG_binding_norm_ (by the number of interfacial residues [26]) to be taken as target functions for the training set consisting of 3200 non-redundant high resolution PPI complexes (targets) (see section 2.1, **Materials and Methods**). Consequently, the normalized binding free-energy values could be interpreted as average interfacial contribution to the overall binding free energy of the interactions which also matched the numerical ranges to be effectively used as a target function in SVM regression. Given the near-native element in FoldX [55], its utilization ensured a sufficiently large dataset for training, scaling to those of the modern-day supervised learning methods (rather than having to rely on a few hundreds of experimental data-points which all we could procure for independent validations).

In order to implement a rigorous training procedure, a large dataset was needed and chosen (as described above) and a ten-fold cross validation was performed during the EnCPdock training – which is a standard technique for training SVM regressors [66–68]. To that end, the whole dataset was first divided into ten equal parts (subsets) yielding a statistically significant number of datapoints (320 PPI complexes) in each subset. The details of the training and cross-validation is to be found in **Supplementary Note S2**. Finally, a maximum correlation (r) value of 0.745 (**Fig. 1**) was obtained between the target function (ΔG_FoldX_norm_) and the predicted output (ΔG_EnCPdock_norm_) with a corresponding maximum BACC score of 0.833. The values of training parameters (C, γ and ε, see section 2.2.2, **Materials and Methods**) exhibiting the highest BACC score and maximum r values (**Fig. S1, Table S1** in **Supplementary Materials**) were then accumulated (closely ranged between 0.744 to 0.745) and considered for independent validations. The corresponding ROC-AUC of TPR vs. FPR plots (see section 2.2.3.1, **Materials and Methods**) were found to be ~0.75 (**Fig. S2** in **Supplementary Materials**). For each of the combinations of C, γ and ɛ in **Table S1**, ten models were obtained (pertaining to each cross-validation subset) resulting in a total of 90 models that demonstrated excellent correlations between the target function and the predicted outputs. Thus, a target test PPI complex undergoing EnCPdock predictions will have 90 predicted outputs. The central tendencies (mean, median, mode) along with maximum and minimum values of these predictions were studied in great detail. Most distributions were not really symmetric, rather than varying from left-skewed, bi-modal, multi-modal to right-skewed (**Fig. S3** in **Supplementary Materials**) which led us to opt for the ‘median’ as a better measure of central tendency than mean or mode – which was set as the final measure for the predicted ΔG_binding_norm_ in EnCPdock. The finally selected 90 models were then taken forward for subsequent independent validations to eliminate the effect of over-fitting that might have occurred during the course of cross-validation.

**Figure 1.**
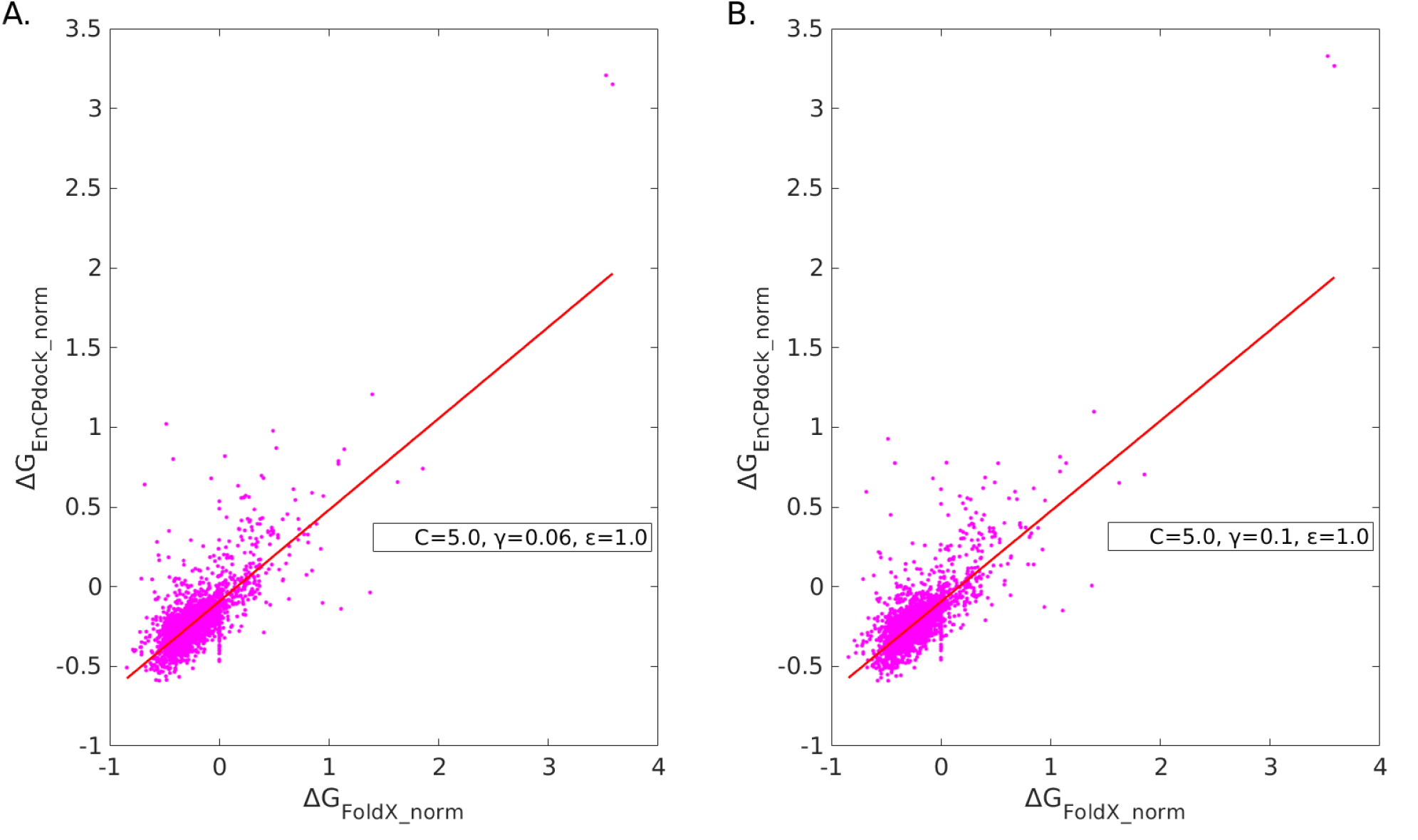
Cross-validated correlation (scatter) plots. The predicted output of ΔG_binding_norm_ from EnCPdock (ΔG_EnCPdock_norm_) for 3200 PPI complexes (during cross-validation) plotted against their corresponding average interfacial binding free energies predicted by FoldX (or ΔG_FoldX_norm_) for two top-performing training sets trained at particular sets of values for of C, γ and ɛ (values provided in insets of panels A, B) both of which exhibited BACC scores of 0.833 and r values of 0.745.

### 3.2. Feature Trends and Contributions

Numeric trends and contributions of individual features in the overall cross-validated performance of EnCPdock were investigated in the training dataset by (i) estimating their descriptive statistics (mean, standard deviations, relative frequency distributions) followed by (ii) enumerating their (Pearson’s) correlations (r) with the predicted ΔG_binding_norm_. Shape Complementarity (Sc), being a necessary criteria (acting like a threshold filter) for inter-protein associations, gave rise to a sharp, narrowly ranged (mildly left-skewed) distribution with a high mean value (0.73; SD^5^: 0.07, **Table 1**) – which, as could be anticipated, is consistent with the short range nature of (van der Waals’) force responsible for it. In distinct contrast, electrostatic complementarity (EC), being resulted from a long range (Coulomb’s) force has a much broader width with only a modest average (0.18; SD: 0.22, **Table 1**) in the training dataset. This again is consistent with the earlier findings that EC can be favorable as well as disfavored in bound protein complexes [25,32], wherein, the later, less frequent event (in ~20% cases) often signals for transient interactions [26]. Other features gave characteristic distributions, mostly tight and uni-modal, sometimes, with a few low-frequency (outlier-like) disjoint borderline modes (**Fig. 2**). Among all features, ACI has arguably the thinnest width (or, the sharpest peak) while on the other hand, the chain-length features, logN and log_asp_ account for large variability in the PPI complexes.

**Figure 2.**
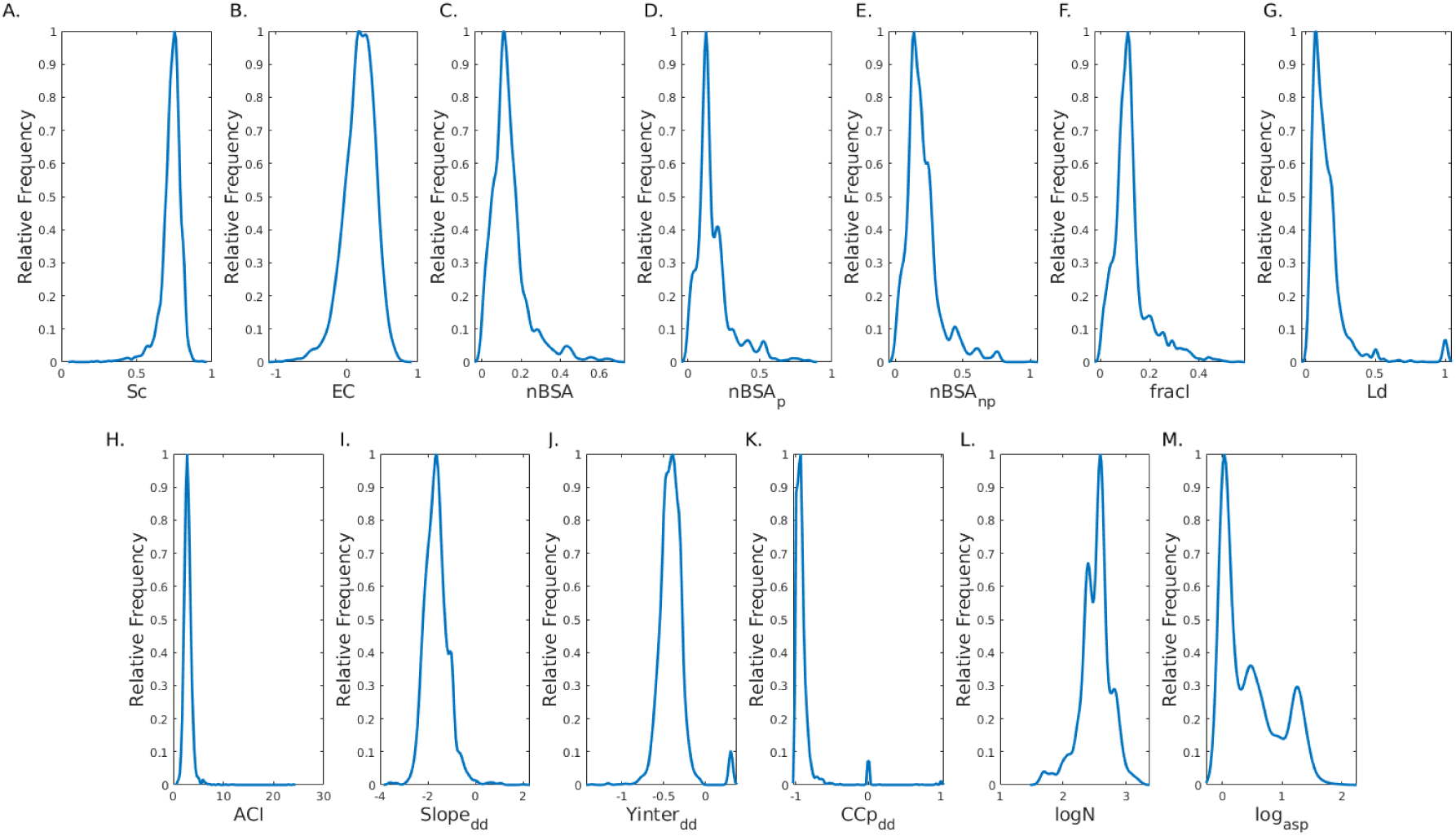
Trends of Features in terms of their relative frequency distribution patterns. Relative frequencies (with respect to the highest observed frequencies for each feature) were computed in the observed ranges (X-axes) individually for all features (Panels. A – M) using kernel density distribution functions (MATLAB, v. R2018b) and plotted against the corresponding feature scores.

**Table 1.**
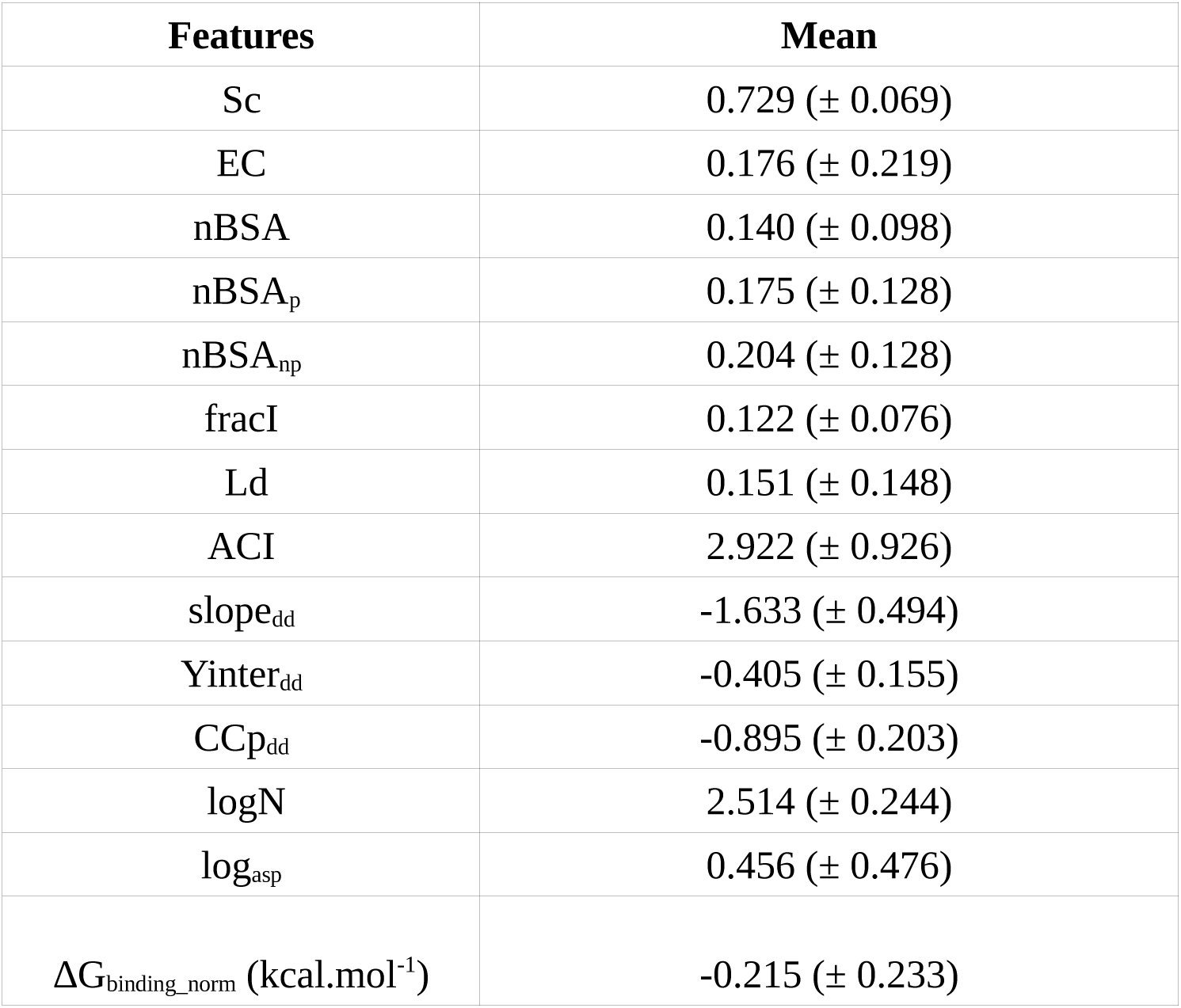
Feature trends in terms of descriptive statistics. Descriptive statistics are furnished in terms of mean and standard deviations (within parentheses) for all features. Alongside, the statistics for the predicted output (ΔG_binding_norm_) is also tabulated.

| Features | Mean |
| --- | --- |
| Sc | 0.729 (± 0.069) |
| EC | 0.176 (± 0.219) |
| nBSA | 0.140 (± 0.098) |
| nBSA<sub>p</sub> | 0.175 (± 0.128) |
| nBSA<sub>np</sub> | 0.204 (± 0.128) |
| fracI | 0.122 (± 0.076) |
| Ld | 0.151 (± 0.148) |
| ACI | 2.922 (± 0.926) |
| slope<sub>dd</sub> | -1.633 (± 0.494) |
| Yinter<sub>dd</sub> | -0.405 (± 0.155) |
| CCp<sub>dd</sub> | -0.895 (± 0.203) |
| logN | 2.514 (± 0.244) |
| log<sub>asp</sub> | 0.456 (± 0.476) |
| $\Delta G_{\text{binding\_norm}}$ (kcal.mol<sup>-1</sup>) | -0.215 (± 0.233) |

To analyze the relative contributions of the different features to the overall (cross-validated) performance of EnCPdock, they were first individually surveyed by correlation (scatter) plots plotted against the predicted output (ΔG_binding_norm_), followed by recording their corresponding Pearson’s correlations (r). It is perhaps good to note that these correlations are not additive and the associated signs (+/-) in the correlations simply refer to the directionality of the feature with respect to the predicted output, wherein, the canceling effect of terms naturally gets nullified during their training (via the non-linear combinatorial optimization in the Support Vector Regression machine). Thus, higher the magnitude of the correlation (irrespective of the sign), higher is the feature contribution. Let’s recall that the overall cross-validated performance of EnCPdock has an r-value of ~0.75. Interestingly, buried surface area based terms as well as most network parameters performed particularly well wherein average contact intensity (ACI) and Link density (Ld) had the best individual correlations (0.51, 0.43 respectively) (**Fig. 3**). Complementarity based terms showed an opposite directionality to the predicted output. Sc, being the threshold filter attained a fairly high correlation (−0.37), while, the low correlation in EC (−0.16) accounted for the variability in terms of electrostatic match or miss-match at the interface of the complexes.

**Figure 3.**
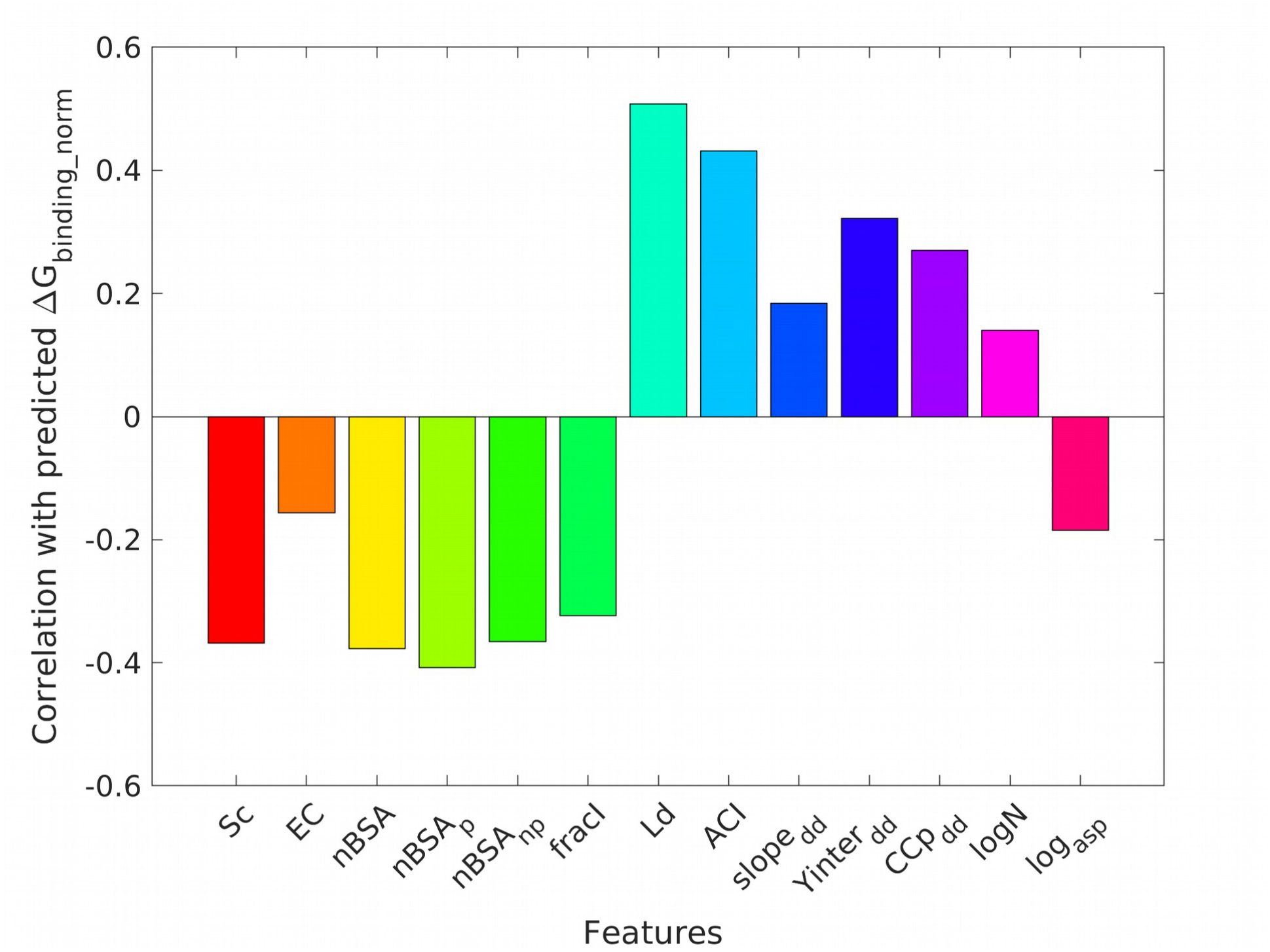
Feature Contributions in terms of correlation with the predicted output. The Y-axis represents the Pearson’s correlation (r) value for each feature against the predicted output (ΔG_binding_norm_), displayed as colored bars. Similar features have been put with nearer colors in the color bars.

### 3.3. Independent Validations on available Experimental Benchmarks

To check for any bias (and eliminate) that might have occurred from the training dataset during cross validation, or, in other words, to take care of over-fitting issues in training, it is customary to perform independent validations on datasets, strictly non-overlapping with the training set. To that end, we attempted to compile databases consisting of native PPI complexes with annotated experimental affinity (K_d_ and/or ΔG_binding_) values. We faced numerous difficulties browsing through different published ‘readily available annotated’ datasets (see **Supplementary Note S1**.) to accumulate such complexes with enough number and with the desired accuracy of their experimental affinities in our repeated web-search. Most databases lacked enough structural (PDB) information for the interactions as well as wild type affinity data (K_d_ and/or ΔG_binding_) and was more populated with mutant thermodynamic data (ΔΔG_binding_).

One of the prime sources, an Affinity benchmark dataset (v.1), once made available was recently removed from the public domain (https://bmm.crick.ac.uk/~bmmadmin/Affinity/). One of the significant findings in our study was the discovery of 3 new targets as mentioned in the updated benchmark, v.2.114 of them were binary and were already taken into account. Subsequent declarations were made with regard to detected inaccuracies in the dataset (https://github.com/haddocking/binding-affinity-benchmark) in terms of the reported experimental thermodynamic parameters (ΔG_binding_ and/or K_d_) and an upgraded version of the dataset was made available by incorporating fresh new entries (though accounting for comparatively a lower number of targets) having the desired accuracy. Together, from earlier [12] and updated versions [69] of the Affinity benchmark (v.2), 225 PPI complexes were first assembled out of which 46 complexes were new entries that could be considered fully accurate (https://github.com/haddocking/binding-affinity-benchmark).

Barring these experimental inaccuracies and updates, these affinity benchmark datasets (both versions) however ensured sampling of a diverse spectrum of PPI complexes in terms of biological functions (covering G-proteins, receptor extracellular domains, antigen/antibody, enzyme/inhibitor, and enzyme/substrate complexes), as well as in terms of affinities (from low to high) of the binding partners with K_d_ ranging from 10^−5^ to 10^−14^ M [12,69] and was well categorized into different classes of comparative difficulty of affinity prediction (rigid body, medium, difficult) etc.

As a methodology, we had chosen to restrict our calculations strictly to binary complexes^6^, also compelled by the fact that EnCPdock was trained on binary PPI complexes (see section 2.1, **Materials and Methods**). This reduced the number of complexes to 106 (see **Dataset S2** in **Supplementary Materials**) which have crystallographic resolutions in the range of 1.1-3.5 Å, and R-factors (working set) within the range of 0.125 to 0.252. This accumulated set contained 31 binary complexes out of the new 46 entries.

During the revision of the manuscript, we looked into the literature and doubled our efforts to retrieve more available experimental affinity data (with proclaimed accuracy) for protein-protein binding in an extensive second round of search. A second dataset was thus assembled by merging the available wild-type thermodynamic data retrieved from Proximate (https://www.iitm.ac.in/bioinfo/cgi-bin/PROXiMATE/wild.py) [42] and SKEMPI 2.0 (https://life.bsc.es/pid/skempi2/) [43] datasets, which together gave us a second experimental benchmark for independent validation consisting of 236 PPI targets with crystallographic resolutions in the range of 1.1-4.4 Å, and R-factors (working set) within the range of 0.125 to 0.321.

To start with the post-prediction (performance evaluation) analyses, ΔG_binding_norm_ values predicted by EnCPdock (let’s call them ΔG_EnCPdock_norm_) for each of the 106 and 236 targets compiling the two filtered independent datasets (I. Affinity benchmark II. SKEMPI+Proximate – merged) were plotted separately against their corresponding experimental values (ΔG_exp_norm_) and FoldX – derived values (ΔG_FoldX_norm_) [16,17] into two distinct sets of pairwise 2D scatter plots (as a mean to compare) (**Fig. 4, Fig. 5**).

**Figure 4.**
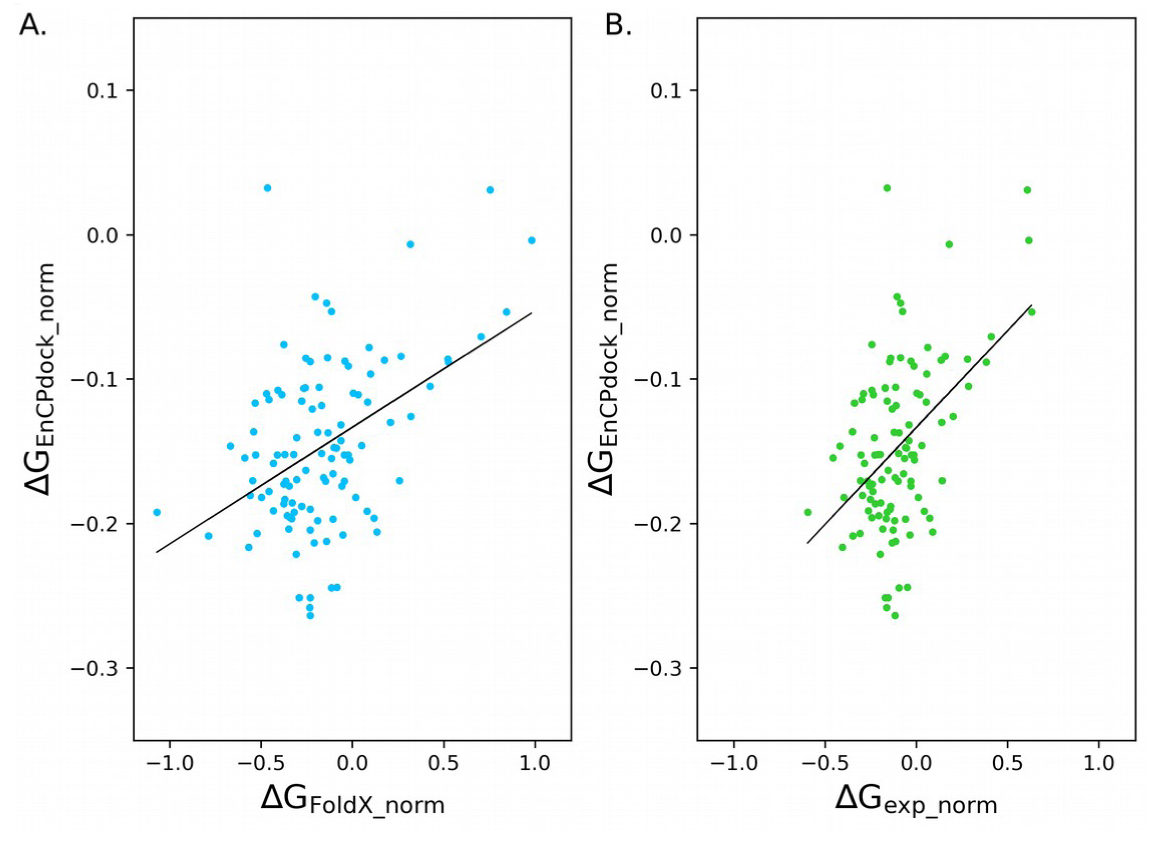
Correlation (scatter) plots in independent validation - 1. The predicted output of ΔG_binding_norm_ from EnCPdock (ΔG_EnCPdock_norm_) for 106 PPI complexes (during independent validation – 1: on the Affinity Benchmark (v.2)) plotted against their corresponding average interfacial binding free energy predicted by FoldX (ΔG_FoldX_norm_) and average interfacial binding free energy estimated by experimental (calorimetric, spectroscopic etc.) means (ΔG_exp_norm_), plotted in panels A and B respectively (with their least-squares fitted lines).

**Figure 5.**
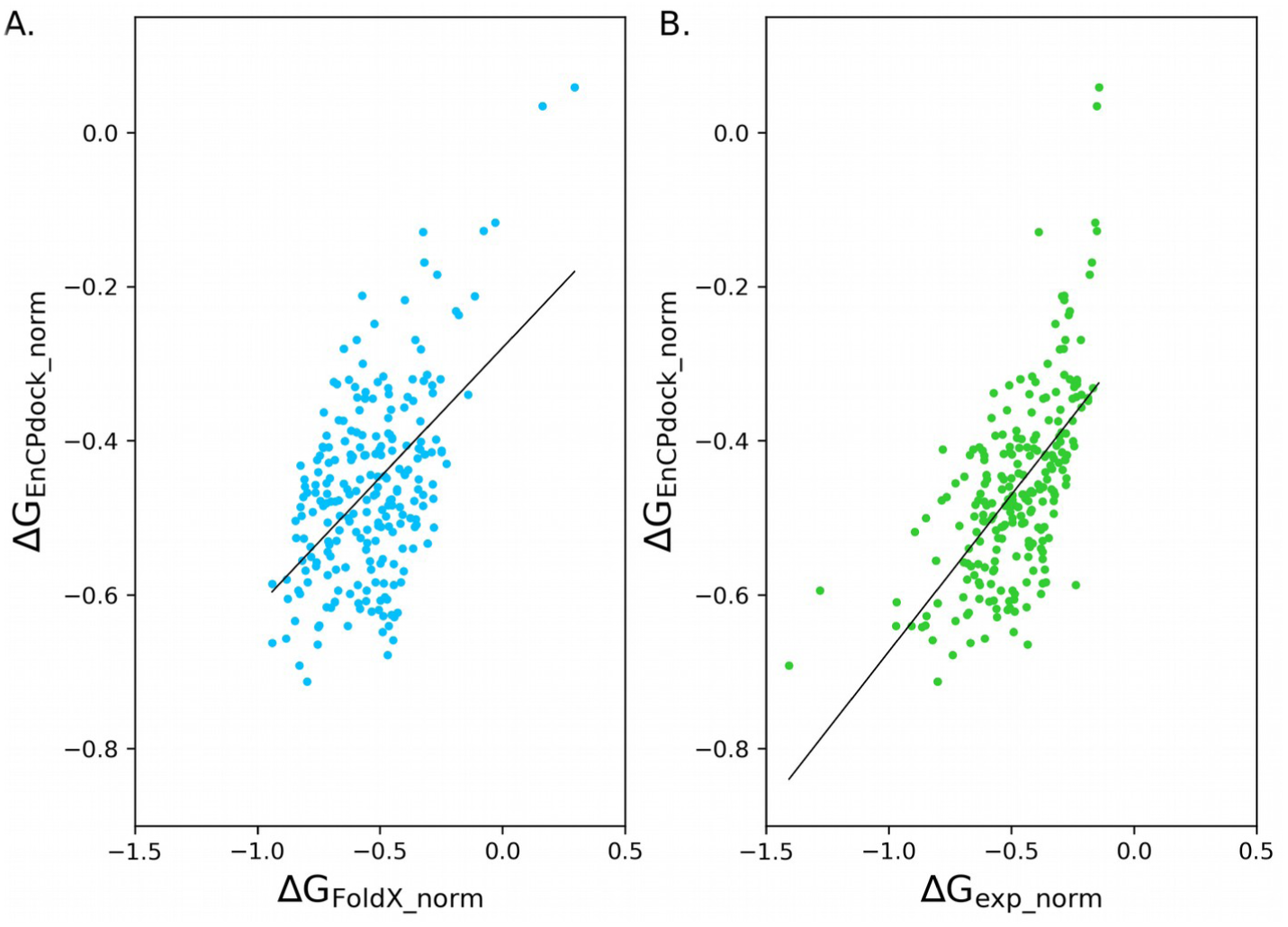
Correlation (scatter) plots in independent validation - 2. The predicted output of ΔG_binding_norm_ from EnCPdock (ΔG_EnCPdock_norm_) for 236 PPI complexes (during independent validation – 2 : on the SKEMPI+Proximate – merged dataset) plotted against their corresponding average interfacial binding free energy predicted by FoldX (ΔG_FoldX_norm_) and average interfacial binding free energy estimated by experimental (calorimetric, spectroscopic etc.) means (ΔG_exp_norm_), plotted in panels A and B respectively (with their least-squares fitted lines).

Note, each of the 90 models predicts ΔG_EnCPdock_norm_ for each target, leading to an array of that many ΔG_EnCPdock_norm_ scores for each target. To check the comparative trends among statistical measures computed from and representative of these 90 values, first, in the first of the two datasets (Affinity Benchmark: 106 targets), the central tendencies (mean, median and mode: ΔG_EnCPdock_norm-mean_, ΔG_EnCPdock_norm-median_, ΔG_EnCPdock_norm-mode_) and the extreme values (minimum and maximum: ΔG_EnCPdock_norm-minimum_, ΔG_EnCPdock_norm-maximum_) were plotted individually, twice for each measure, against (i) taking ΔG_FoldX_norm_ and (ii) ΔG_exp_norm_ as benchmarks. All statistical measures led to similar scatter plots (**Fig. S4** in **Supplementary Materials**) and similar range of correlation values (**Table S2** in **Supplementary Materials**) which ensured similar predictive abilities of all 90 models. As reasoned earlier, ‘median’ was chosen as the preferred statistic over the other measures (see section 3.1). In line with that, the primary comparison was carried out between the corresponding ‘median’ values in both datasets for independent validations (**Fig. 4, Fig. 5**) while the other statistical measures were also tested side-by-side in the first (Affinity benchmark) of the two datasets. All these other statistical measures (ΔG_EnCPdock_norm-mean_, ΔG_EnCPdock_norm-mode_, ΔG_EnCPdock_norm-minimum_, ΔG_EnCPdock_norm-maximum_) exhibited correlations slightly lesser than that of ΔG_EnCPdock_norm-median_ (**Table S2**), consistently in both the cases (against ΔG_exp_norm_ and ΔG_foldX_norm_). This also validates the choice of ‘median’ as the correct statistical measure (among the tested alternatives) which automatically leads to a mapping of ΔG_EnCPdock_norm_ → ΔG_enCPdock_norm-median_ in all subsequent use of the term ΔG_EnCPdock_norm_.

The main purpose of carrying out these plots (side-by-side) was to decipher if the performance of EnCPdock (in terms of the predicted ΔG_EnCPdock_norm_ values) obtained against the target function taken in training (i.e., ΔG_FoldX_norm_) is retained (or, if there’s any improvement) when compared to the corresponding experimental (ΔG_exp_norm_) values. To that end, in the left, right panels of these plots (**Fig. 4**, **Fig. 5**), ΔG_FoldX_norm_, ΔG_exp_norm_ values (both serving as the benchmarks) were taken along the X-axis (independent variables) respectively while the corresponding predicted ΔG_EnCPdock_norm_ were taken along the Y-axis (dependent variable).

Linear least squares fitting was performed (in MATLAB, version R2018b) to each of these plots using perpendicular offsets. Any empirical correlations between these methods were first surveyed by visually analyzing these 2D scatter plots (and by the linear least squares fitting performed on them) followed by quantifying the obtained correlations by computing the Pearson’s correlation coefficients (r) between them. EnCPdock performed reasonably well on the Affinity benchmark consisting of 106 binary complexes giving rise to a correlation of r=0.45 between the predicted ΔG_EnCPdock_norm_ and the corresponding FoldX – derived values taken as the benchmark. When compared to the actual experimental values (taken as the benchmark), the performance was not only retained but in fact was slightly improved leading to a correlation of r=0.48 with ΔG_EnCPdock_norm_.

In the second dataset (SKEMPI+Proximate – merged) consisting of 236 binary complexes, EnCPdock performed even better with a correlation of r=0.52 between the predicted ΔG_EnCPdock_norm_ and the corresponding FoldX – derived values (ΔG_FoldX_norm_) taken as the benchmark. In consistency with the earlier finding (that in the Affinity benchmark), the performance was further improved to r=0.63 when compared to the actual experimental values (ΔG_exp_norm_) taken as the benchmark. All correlations were significant at 99.9% level (as revealed from their corresponding p-values).

It is indeed interesting to note that ΔG_EnCPdock_norm_ (in spite of being trained on ‘near-native’ FoldX estimates) correlates actually better than ΔG_FoldX_norm_ values taken as benchmark (r=0.45, 0.52 for Affinity benchmark (v.2), SKEMPI+Proximate – merged datasets respectively) as compared to the corresponding ΔG_exp_norm_ values (r=0.48, 0.63 – in the same order). One plausible rationale behind this seemingly non-trivial finding is most probably the use of fine-grained structural descriptors like shape and electrostatic complementarities (Sc, EC [23,25]) in EnCPdock which are surface-based features that have been benchmarked over decades of careful parameterization [25,27,28,32,37,49] on upgraded lists of available native high-resolution crystal/cryo-EM structures. Together, as an ordered pair, {Sc, EC} takes into account the entire spectrum of long-range (electrostatic) as well as local (shape) effects in finest detail. In effect, most physical (short and long range) forces acting on the complex is inherently taken care of in EnCPdock, thereby making the predictions perhaps even more native-like. Furthermore, the results may also be envisaged as a performance booster of FoldX (or similar semi-empirical force-field based methods) when used as a target function in an advanced AI-predictor compared to when used as the exclusive predictor itself. This leaves the scope to come up with a plausible hybrid method combining FoldX and EnCPdock to raise the performance even more in the future.

As a cross-check, we also estimated the performance of FoldX (ΔGG_FoldX_norm_) directly against experimental affinity data (ΔGG_exp_norm_) and obtained correlations of r=0.67, 0.70 for the two independent validation benchmarks respectively. These correlations (r) and their corresponding significance (p-value <0.00001 for both at 99.9% level) clearly reflect the near-native element in FoldX, consistent across both independent benchmarks.

Further, to have an idea of the Receiver Operating Characteristics of the free-energy predictor (ΔG_binding_norm_) in EnCPdock, the final ΔG_EnCPdock___norm_ scores for each target in both independent validation benchmarks were converted to 1-0 binary variables (0: stable, 1: unstable), independently, in tandem with ΔG_exp_norm_ and ΔG_FoldX_norm_ taken as the point of reference. In the two independent calculations, the conversion from floating to binary variables were based on a set of cutoffs (threshold values) spanning the ranges of ΔG_exp_norm_ and ΔG_foldX_norm_ respectively. Subsequent to this conversion, a set of TPR, TNR and BACC scores (see section 2.2.3, **Materials and Methods**) were computed for each sampled cutoff on ΔG_exp_norm_ and ΔG_foldX_norm_ respectively (**Table S3, Table S4** in **Supplementary Materials**). Among different sampled cutoffs, the free-energy predictor (ΔG_binding_norm_) in EnCPdock attained to a maximum BACC score (BACC_max_) of 0.892, 0.922 and 0.884, 0.777 with respect to FoldX and experimental values taken as benchmarks respectively (**Table S3, Table S4**) in I. Affinity benchmark, II. SKEMPI+Proximate – merged datasets.

### 3.4. Comparison with the state-of-the-art in affinity predictions

Decades long effort have been devoted in structure-based predictions of equilibrium thermodynamic parameters of protein-protein binding with gradual (linear) amplification of available experimental data of the desired quality. The history goes back to the early days of the trivial use of accessible surface area measures (ΔGASA) which never (systematically) correlated well (even for the rigid complexes) with binding free energy (r=−0.16) barring a few case studies with much of empirical (*ad-hoc*) formalism [10]. It was found that both large and not-so-large ΔGASA-complexes could give rise to high affinities [69]. To that end, it was clear that change in solvent accessible surface area upon binding was definitely not the sole determinant of binding affinity in proteins. Consequently, root-mean-square deviation measures defined on the interface residues alone (I-RMSD) were attempted as an affinity-predictor which also gave only a modest correlation with ΔG_binding_ (r=−0.24), and only a nominal improvement was found using a minimal linear model combining ΔGASA and I-RMSD (r=0.31) [70]. In the latter years, a coulomb based model (defined over interfacial distances) was used to determine the effect of electrostatic force on PPI complexes which also could not lead to much higher a correlation [71]. Also, since, protein structure and function affect the interaction and binding affinity of proteins, sequence-based prediction is challenging.

We are now in the era of the sophisticated ‘affinity scores’ - many of which (like that of our method) are AI-trained. Machine learning is certainly preferred among computational methods for predicting the binding affinity of proteins due to its implicit consideration of any factors affecting PPIs and the flexibility of using empirical data instead of a fixed or predetermined function. The correlation (with ΔG_binding_) was further improved by prediction methods (affinity scores) including the specific geometry and composition of the interaction, raising the overall correlations of up to r=0.53 over a collection of flexible and rigid targets [69]. The best of these methods were trained either using an earlier version of the affinity benchmark [12] or using changes in affinity upon mutation(s) [72]. Interestingly, in spite of this elevated performance in the earlier benchmark (reported best correlation of r=0.63), these methods [73–75] could not better their performance in the updated benchmark [69]. Consequently, performance of 12 commonly used algorithms compiling affinity scores as well as free energy predictors were found to give poor correlations on the earlier version of the benchmark consisting of 81 PPI complexes (https://github.com/haddocking/binding-affinity-benchmark) which slightly improved (up to r=0.53) on the subset of 46 fully accurate complexes [69]. Among MM(P/G)BSA approaches [19], the strongest correlation observed was r=−0.64; by invoking MM-GBSA without considering conformational entropy effects in a low interior dielectric constant of 1 [76] – which outperformed the results obtained by MM-PBSA (r=−0.52). The highest reported correlation [8] we could procure from the literature was r=–0.73, by one of the affinity scores, using a linear regression model trained on ‘inter-residue contact (IC)’ descriptors of affinity, categorized based on their residue hydrophobicities (e.g., polar–polar, polar–nonpolar, etc.)’ [13].

In this background, here, in this paper, we tested the operational credibility of the EnCPdock system (www.scinetmol.in/EnCPdock/) to probe binding energetics in protein interactions. Let’s recall that unlike other predictors, EnCPdock is a comprehensive web-interface for direct conjoint comparative analyses of complementarity and binding energetics in inter-protein associations. Trained by non-linear support vector regression machines on thirteen judiciously chosen high-level structural descriptors (and on FoldX – derived free energies as its target function), EnCPdock returns a performance of r=0.745 (with associated BACC score of 0.833, **Table S1**) in cross-validation over 3200 PPI complexes and r=0.48 against actual experimental data (ΔG_binding_norm_) in an independent validation performed on a filtered set of 106 binary complexes, culled from the updated benchmark. Thus, considerable retention of performance is reflected in the independent validation on the updated affinity benchmark, comparable to the state-of-the-art. In addition, EnCPdock returns the mapping of the binary PPI complex in the Complementarity Plot and further returns mobile molecular graphics of the atomic contact network probed at the protein-protein interface for further analyses.

### 3.5. Case studies demonstrating the utilities of EnCPdock in probing inter-protein associations of various types and biological origins

EnCPdock has been built to exercise complementarity and energetics from a common interactive platform to probe protein – protein interactions. Currently the web-server runs only for binary protein (receptor – ligand) complexes, with a future plan to cover even more complex oligomeric protein associations. To demonstrate its utility in protein energetics, individual cases were studied in finer details, sampling a diverse plethora of biological origin and function of protein binary complexes. These included antigen – antibody interactions, protein – inhibitor complexes, interactions involved in signal transduction pathways, enzyme – substrate complexes and others.

One such example was that of a RAB GTPase bound to Rabenosyn-5, its cognate ligand (an effector molecule) (PDB ID: 1Z0K, [77]) involved in different stages of membrane trafficking, serving as a transport protein. The complex was found to raise an Sc value of 0.7 with an EC value of 0.12 which together made it fall into the ‘probable’ region of the Complementarity plot (CP_dock_) (**Fig. 6.A, Table 2**). The inter-chain contact network spanned across a densely connected extended interface involving two α-helices of the ligand and primarily β-sheets coming from the receptor (**Fig. 6.B**). The Link density (Ld) of the interfacial network was found to be 0.11 with an average contact intensity (ACI) of 2.89 (see section 2.2.1.6, **Materials and Methods**). Scale-freeness was evident from the degree distribution profile based features (combined) with a Pearsons correlation (r) of −0.987 (see section 2.2.2, **Materials and Methods,**). The interface involved 22 residues combining both partners which eventually gave rise to a predicted normalized (average interfacial) binding free energy (ΔG_binding_norm_) of −0.183 kcal.mol^−1^ – which was much closer to the experimental (Calorimetric [78]) value for the same (−0.196 kcal.mol^−1^) as compared to FoldX predictions (−0.330 kcal.mol^−1^).

**Figure 6.**
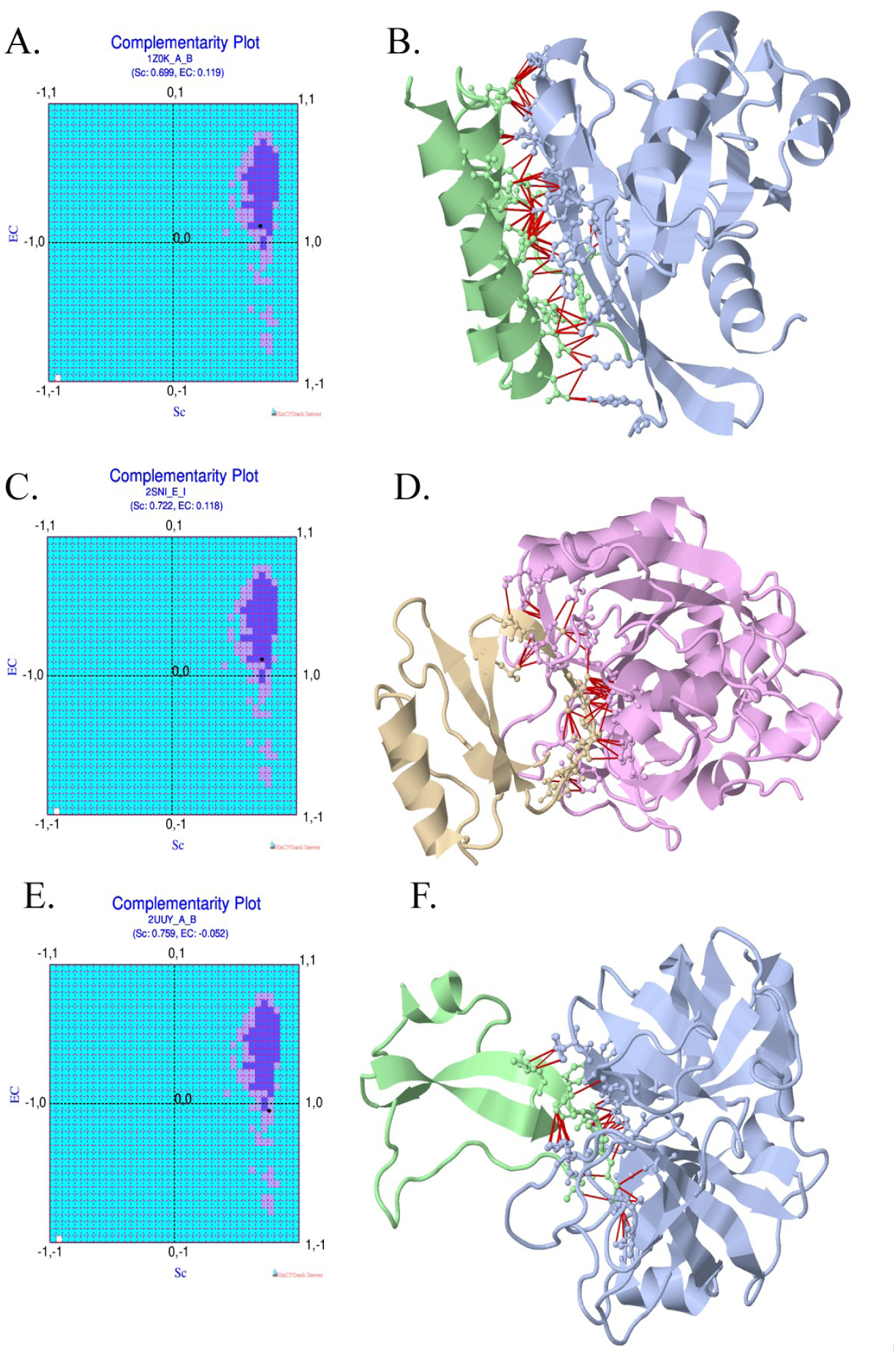
EnCPdock outputs for three different PPI complexes, taken as demonstrative case studies. The Complementarity Plots (CP_dock_) (panels A, C, E) and the interfacial interaction networks (panels B, D, F) returned by the EnCPdock web-server for the three chosen complexes with PDB IDs: 1Z0K (A and B respectively), 2SNI (C and D respectively) and 2UUY (E and F respectively) are portrayed.

**Table 2.** Feature scores and predicted free energies returned by EnCPdock for the demonstrative examples chosen as case studies.

| Features | PDB ID |  |  |
| --- | --- | --- | --- |
|  | 1Z0K | 2SNI | 2UUY |
| $N_{\text{intres}}$ | 32 | 22 | 19 |
| Sc | 0.699 | 0.722 | 0.759 |
| EC | 0.119 | 0.118 | -0.052 |
| nBSA | 0.166 | 0.128 | 0.109 |
| nBSA<sub>p</sub> | 0.155 | 0.101 | 0.095 |
| nBSA<sub>np</sub> | 0.175 | 0.149 | 0.120 |
| fracI | 0.217 | 0.142 | 0.156 |
| Ld | 0.105 | 0.134 | 0.155 |
| ACI | 2.889 | 3.200 | 2.692 |
| Slope<sub>dd</sub> | -1.545 | -2.179 | -1.657 |
| Y<sub>inter</sub><sub>dd</sub> | -0.434 | -0.299 | -0.312 |
| CC<sub>pdd</sub> | -0.987 | -0.979 | -0.982 |
| logN | 2.361 | 2.530 | 2.439 |
| log<sub>asp</sub> | 0.443 | 0.633 | 0.632 |
| $\Delta G_{\text{EnCPdock\_norm}}$<br>(kcal.mol<sup>-1</sup>) | -0.183 | -0.183 | -0.170 |
| $\Delta G_{\text{FoldX\_norm}}$<br>(kcal.mol<sup>-1</sup>) | -0.330 | -0.365 | -0.158 |
| $\Delta G_{\text{exp\_norm}}$<br>(kcal.mol<sup>-1</sup>) | -0.196 | -0.268 | -0.115 |

One other example was that of a protein – inhibitor complex (PDB ID: 2SNI [79]) involving Subtilisin Novo, a Serine-Protease and its inhibitor. The complementarity values were much similar to the earlier complex (Sc: 0.72, EC: 0.12) again mapping to the ‘probable’ region of CP_dock_ (**Fig. 6.C, Table 2**). The inter-chain contact network was more extended (involving 32 residues) also mapped to a similar (link) densely (Ld: 0.13) however mostly involving β-sheets and loops coming from both binding partners (**Fig. 6.D**) with an even higher ACI value of 3.2 (see **Materials and Methods**). Scale-freeness was also evident here (r: −0.979) as with most bipartite PPI networks (see **Materials and Methods**). The complex lead to a similar predicted ΔG_binding_norm_ value of −0.182 kcal.mol^−1^ – which again was nearer to the experimental (Calorimetric [80]) reference (−0.268 kcal.mol^−1^) as compared to FoldX predictions (−0.365 kcal.mol^−1^).

Yet another example was also that of a protein – inhibitor complex from the Hydrolase family involving bovine trypsin and a tick tryptase inhibitor (PDB ID: 2UUY [81]). This complex presented a repulsive electrostatic interaction (EC: −0.052) compensated by stronger shape constraints (0.76) in terms of complementarity. Such compensation has been found in native PPI complexes at a statistically significant extent (~20% of all native inter-protein interactions), often representing transient interactions [25,26,32]. Consistent with these (statistical) estimates, here, the {Sc, EC} point fell into the ‘less probable’ region of CP_dock_ (**Fig. 6.E, Table 2**). Here the interface was less-extended (involving 19 residues), albeit, more dense (Ld: 0.16, ACI: 2.69), again involving primarily β-sheets and loops coming from both binding partners (**Fig. 6.F**). As with the other examples, scale-freeness was also evident here (r: −0.98). The complex here led to a similar predicted ΔG_binding_norm_ value of −0.17 kcal.mol^−1^ – which was comparable to FoldX predictions (−0.158 kcal.mol^−1^) with respect to the experimental [82] benchmark (−0.115 kcal.mol^−1^). Together these examples adequately demonstrate the apt usage of EnCPdock as a combined graphical and analytical platform to survey complementarity and energetics attributed to protein – protein interactions.

### 3.6. Application of EnCPdock in probing peptide binding specificity and mutational effects

In principle, EnCPdock can also be used effectively to probe protein – peptide interactions and mutational effects. In order to test this, the human protein T-cell Lymphoma Invasion and Metastasis-1 (PDZ domain) was chosen in its native (PDB ID: 4GVD) and available Quadruple Mutant (QM) forms, in complex with similar classes of (native: Syndecan1; QM: caspr4) peptides (PDB ID: 4NXQ) [83]. The QM is defined to be the simultaneous four mutations: L911M, K912E, L915F and L920V in the native receptor chain. The availability of crystal structures and related experimental affinity data (although, indirect: spectroscopic) for analogous complexes [84] allowed us to carry out their direct comparison in EnCPdock. The native protein (4GVD) has two symmetry-related homologous bio-assemblies (chains: A-D; B-C), while the mutant form has three (chains: A-D, B-E, C-F). In effect, the asymmetric units of the native and the mutant forms contain (homo-) dimeric and trimeric complexes. All bio-assemblies of the native and the mutant were individually surveyed in EnCPdock, giving similar estimates in terms of most high-level structural descriptors (**Table 3**), yet, their cumulative difference adding to effective measures of (binding) affinity and stability (mutant vs. native) could be probed unambiguously in terms of the predicted energetics (**Table 3**). The interaction of these classes of proteins (taken as receptors) and peptides (ligands) seem to have a characteristic suboptimal electrostatic complementarity (EC), often associated with transient (or, quasi-stable) complexes [26] – wherein the repulsive electrostatic effect are usually compensated by stronger shape constraints [25,32]. Indeed, all bio-assemblies pertaining to both the native and the mutant, unambiguously gave rise to negative EC values (EC_native_4GVD_: −0.515 (A-D), −0.277 (B-C); EC_mutant_4NXQ_: −0.354 (A-D), −0.608 (B-E), −0.282 (C-F)) indicative of their quasi-stability. As is characteristic with such metastable interactions, the corresponding shape complementarities (Sc) were found to be appreciably high, closely ranging between 0.69 to 0.73 among all complexes. Consequently, the {Sc, EC} ordered pair points (corresponding to all complexes) had fallen to the 4^th^ quadrant of the Complementarity Plot (CP_dock_) only managing to hit ‘less probable’ or ‘improbable’ regions of the plot (**Fig. 7**). The combined drop in stability and affinity (in terms of binding energetics) in the mutant compared to the native complexes, however, was unambiguous across the different bio-assemblies for both (**Table 3**).

**Figure 7.**
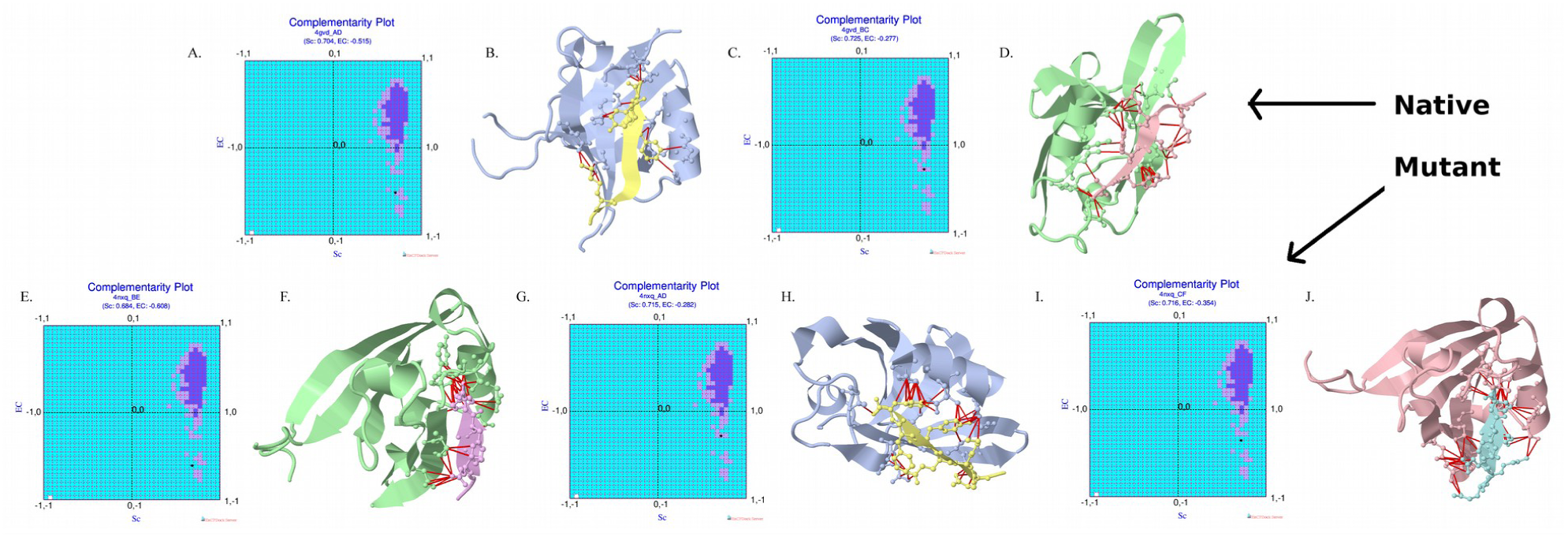
EnCPdock outputs to probe peptide binding specificity and mutational effects. EnCPdock outputs for the native human protein T-cell Lymphoma Invasion and Metastasis-1 protein (PDZ) domain (PDB ID: 4GVD) and its quadruple (QM) mutant (PDB ID: 4NXQ) bound to peptides of similar classes. The Complementarity Plots (CP_dock_) and the interfacial contact networks of the native PPI complex have been analyzed and displayed for its two symmetry-related bio-assemblies (4GVD, chains: A-D, and B-C respectively), and for three symmetry-related bio-assemblies pertaining to the mutant (4NXQ, chains: A-D, B-E and C-F respectively).

**Table 3.**
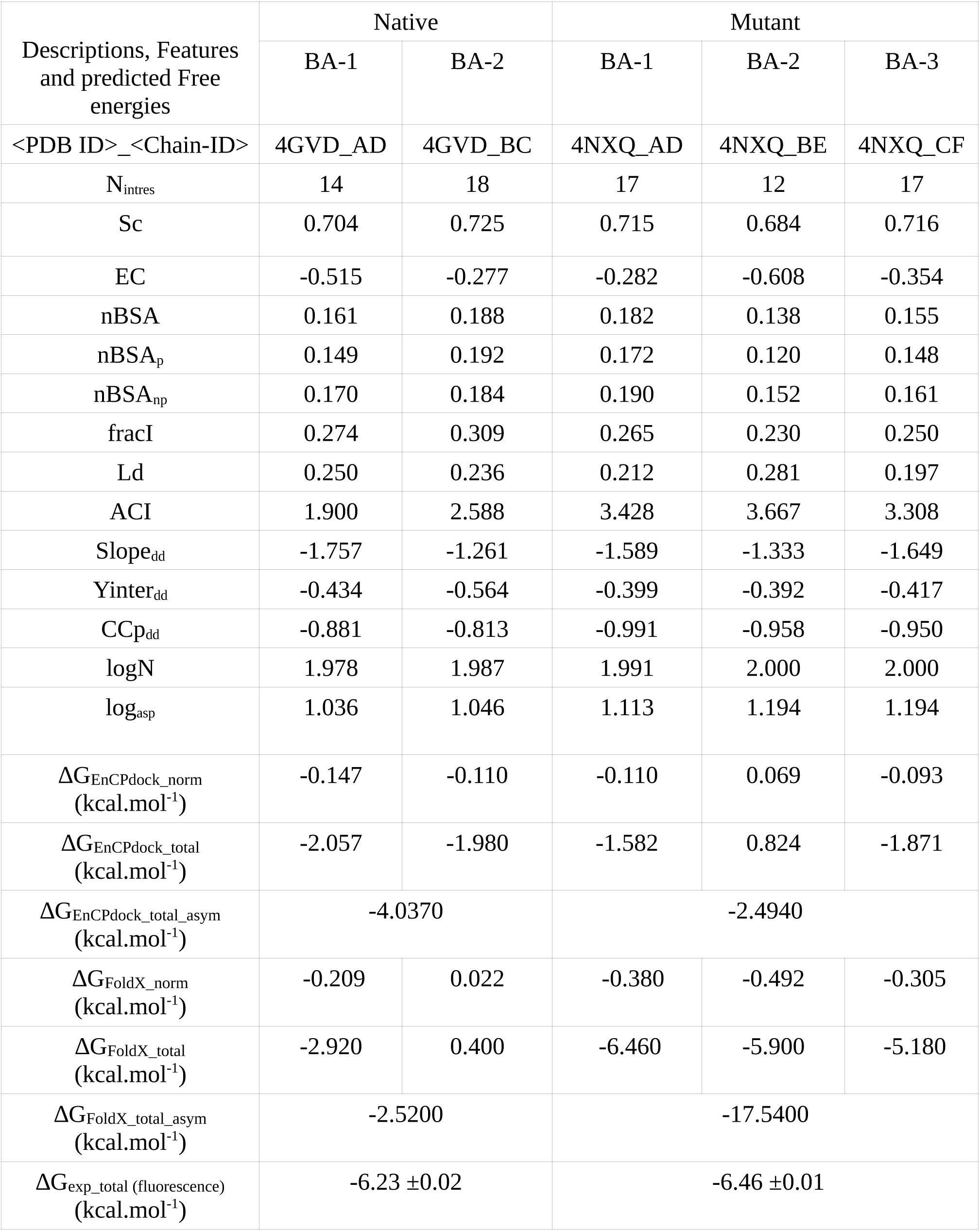
Feature scores and predicted free energies returned by EnCPdock for the peptide-protein complexes used in probing peptide binding specificity and mutational effects. BA stands for Bio-assemblies present in a PDB structure. The native and mutant protein consists of two and three bio-assemblies respectively.

| Descriptions, Features and predicted Free energies | Native |  | Mutant |  |  |
| --- | --- | --- | --- | --- | --- |
|  | BA-1 | BA-2 | BA-1 | BA-2 | BA-3 |
| <PDB ID>_<Chain-ID> | 4GVD_AD | 4GVD_BC | 4NXQ_AD | 4NXQ_BE | 4NXQ_CF |
| $N_{\text{intres}}$ | 14 | 18 | 17 | 12 | 17 |
| Sc | 0.704 | 0.725 | 0.715 | 0.684 | 0.716 |
| EC | -0.515 | -0.277 | -0.282 | -0.608 | -0.354 |
| nBSA | 0.161 | 0.188 | 0.182 | 0.138 | 0.155 |
| nBSA<sub>p</sub> | 0.149 | 0.192 | 0.172 | 0.120 | 0.148 |
| nBSA<sub>np</sub> | 0.170 | 0.184 | 0.190 | 0.152 | 0.161 |
| fracI | 0.274 | 0.309 | 0.265 | 0.230 | 0.250 |
| Ld | 0.250 | 0.236 | 0.212 | 0.281 | 0.197 |
| ACI | 1.900 | 2.588 | 3.428 | 3.667 | 3.308 |
| Slope<sub>dd</sub> | -1.757 | -1.261 | -1.589 | -1.333 | -1.649 |
| Y<sub>inter</sub><sub>dd</sub> | -0.434 | -0.564 | -0.399 | -0.392 | -0.417 |
| CC<sub>p</sub><sub>dd</sub> | -0.881 | -0.813 | -0.991 | -0.958 | -0.950 |
| logN | 1.978 | 1.987 | 1.991 | 2.000 | 2.000 |
| log<sub>asp</sub> | 1.036 | 1.046 | 1.113 | 1.194 | 1.194 |
| $\Delta G_{\text{EnCPdock\_norm}}$ (kcal.mol<sup>-1</sup>) | -0.147 | -0.110 | -0.110 | 0.069 | -0.093 |
| $\Delta G_{\text{EnCPdock\_total}}$ (kcal.mol<sup>-1</sup>) | -2.057 | -1.980 | -1.582 | 0.824 | -1.871 |
| $\Delta G_{\text{EnCPdock\_total\_asym}}$ (kcal.mol<sup>-1</sup>) | -4.0370 | | -2.4940 | | |
| $\Delta G_{\text{FoldX\_norm}}$ (kcal.mol<sup>-1</sup>) | -0.209 | 0.022 | -0.380 | -0.492 | -0.305 |
| $\Delta G_{\text{FoldX\_total}}$ (kcal.mol<sup>-1</sup>) | -2.920 | 0.400 | -6.460 | -5.900 | -5.180 |
| $\Delta G_{\text{FoldX\_total\_asym}}$ (kcal.mol<sup>-1</sup>) | -2.5200 | | -17.5400 | | |
| $\Delta G_{\text{exp\_total}}$ (fluorescence) (kcal.mol<sup>-1</sup>) | -6.23 ±0.02 | | -6.46 ±0.01 | | |

Moving ahead, the ΔG_binding_ for the asymmetric units (ΔG_binding_asym_) were computed by adding the individual ΔG_binding_ for all bio-assemblies (energy being an explicit thermodynamic function, the terms add up) for both native and mutant. These final ΔG_binding_asym_ values were then compared across EnCPdock, FoldX and indirect experimental affinity values available for analogous complexes, probed by fluorescence binding assays [84]. As the numbers got revealed (**Table 3**), FoldX under-predicts the native stability by quite some amount (−2.527 compared to −6.23 ±0.02 kcal.mol^−1^) and over-predicts the mutant stability to a very large extent (−17.54 compared to −6.46 ±0.01 kcal.mol^−1^) with respect to the available experimental data. On the other hand, EnCPdock is much closer in terms of native stability (−4.037 compared to −6.23 ±0.02 kcal.mol^−1^). In terms of comparative stability of the mutant (with respect to the native), although EnCPdock predicts a reverse trend (soft destabilization), there is far lesser error in its predicted ΔG_binding_ (−2.494 in reference to −6.46 ±0.01 kcal.mol^−1^) compared to that of FoldX (−17.54 kcal.mol^−1^). As a result, the absolute value of ΔΔG for EnCPdock (1.553 kcal.mol^−1^) is ~10 times smaller than that of FoldX (−15.02 kcal.mol^−1^) while ΔΔG is just −0.23 kcal.mol^−1^ in terms of the available experimental mean values (i.e., only mild stabilization upon the QM mutation). Overall, in this case study, EnCPdock appears to have better near-native estimates than FoldX – which can plausibly again be rationalized by the ‘sophisticated surface based treatments of the complementarity measures’ as discussed in independent validations.

### 3.7. Feature Trends and Relative Probabilities of Feature-scores – a tool for targeted design of protein interfaces

One of the most effective user-interfaces of EnCPdock is perhaps the ‘Feature Trends’ tab (see help-page: https://scinetmol.in/EnCPdock/help.php) which should be of real practical use in the design of targeted protein/peptide interfaces and/or mutational surveys. This functionality portrays the relative frequency distribution profiles (as in **Fig. 2**) for each feature, computed based on their native kernel densities (see section 3.2). It then plots the obtained feature scores (for an input PPI complex) as vertical red dashed lines and numerically determines the ‘points of intersection’ (highlighted as red dots, **Fig. 8**) between these vertical lines and the corresponding relative frequency distribution profiles. The abscissa of this ‘point of intersection’ (for each feature) can be interpreted as the relative probability (Pr_fmax_) of the obtained feature-score with respect to the event of the highest observed frequency for that feature. These relative probability estimates reflect whether an input PPI complex is regular or terminal with respect to the obtained feature-scores for each feature. This way, the built functionality could be really helpful to pinpoint the structural defect in a targeted inter-protein association to the level of individual structural descriptors. This makes the ‘Feature Trends’ tab (EnCPdock) effective in the structural tinkering and intervention as might be relevant to targeted design of protein-interfaces, mutational studies and peptide design.

**Figure 8.**
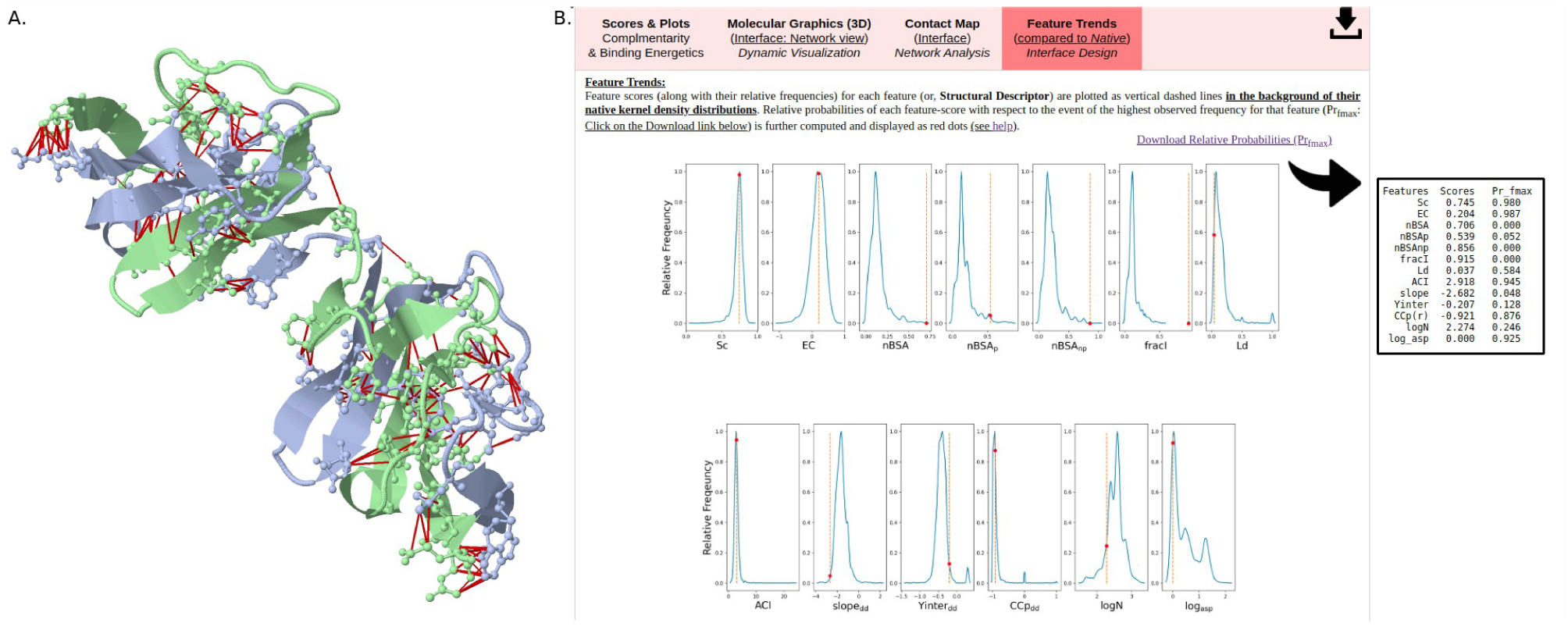
A demonstrative case study of rare lateral inter-twining in homo-dimeric assembly in an engineered Rat CD2 protein that exemplifies the apt and utility of the ‘Feature Trends’ tab in EnCPdock. Panel (A) shows the molecular graphics generated by EnCPdock highlighting the interfacial contact network (links displayed as red lines) which clearly reveal the lateral intertwining of the two chains of 1A64. Panel (B) shows the Feature trends for 1A64 (as returned in EnCPdock) with the relative probability estimates (Pr_fmax_) tabulated in the inset.

One demonstrative example (**Fig. 8**) of the apt and utility of this functionality was found in the case of the engineered form of the otherwise misfolded protein of Rat CD2 (PDB ID: 1A64). While surveying through random control trials, to our great surprise, the fractional interfacial (residue) content (fracI) for 1A64 was found to be ~92% of the whole protein surface (fracI=0.915). In accordance, normalized buried surface area was also high (nBSA=0.706). Naturally, both events being extremely rare in native protein-protein interactions (**Fig. 2**), left their signatures in the corresponding relative probability estimates (Pr_fmax_(fracI)=0.000; Pr_fmax_(nBSA)=0.000). A follow-up visual structural inspection revealed that the two (identical) chains were laterally inter-twined to form a metastable homo-dimer during their assembly – which is said to be characteristic and idiosyncratic of the said protein [85]. Being a homo-dimer, log_asp_ was found to be zero (by definition) which appeared to be a frequent event (Pr_fmax_(log_asp_)=0.925) in native PPI complexes.

## 4. Conclusion

EnCPdock was envisaged and built with the objective of developing a comprehensive web-interface for the conjoint comparative analyses of physico-chemically relevant high-level structural descriptors (complementarity in particular) and binding energetics of interacting protein partners. With that broad objective in mind, the current version of the web-sever provides fine-grained interface properties of binary PPI complexes inclusive of complementarity and other high-level structural features, and, concomitantly predicts their free energies of binding (average interfacial contribution as well as total) from atomic coordinates. In addition, the web-server also generates mobile molecular graphics (in JSmol) of the interfacial atomic contact network and returns the contact map of the interface. Furthermore, trends of individual features (Sc, Ld etc.) can be analyzed against their native (kernel density) distributions. This would be benifitial for structural tinkering and intervention, both in case of comparing among docked poses as well as in the interface design of targeted complexes. A non-linear support vector regression machine has been implemented to train and build EnCPdock, elegantly combining fine-grained structural descriptors of size, shape, electrostatics, accessible and buried surface areas, network parameters of the interfacial contact network etc. from an input protein complex. Trained on this knowledge-base, EnCPdock predicts the corresponding binding free energy with the desired accuracy and reasonable speed (~2 mins for each average-sized input). A detailed documentation of the web-interface can be found in the associated help page (https://scinetmol.in/EnCPdock/help.php) with exhaustive description of its Input-Output (I/O) interface. A flowchart (**Fig. 9**) compiling all the different steps and functionalities (i.e., the entire I/O-interface) involved in EnCPdock has also been appended for easy understanding and navigation of the user. In spite of being trained on near-native FoldX – derived estimates (due to lack of enough experimental data for training), EnCPdock performs comparable to (if not better than) the state-of-the-art in prediction accuracy of binding energetics – *as* revealed by cross-validations on a large dataset. The performance is well retained in the independent validations carried out on multiple experimental benchmarks. Demonstrative examples presented as case studies further show the robustness of EnCPdock across protein complexes coming from a variety of biological origin and types. The utility of EnCPdock in probing peptide binding specificity has also been exemplified, coupled with its use in mutational analyses. The applications can naturally be extended to rank and re-rank docked poses of protein – protein / peptides. Lastly, the ‘Feature Trends’ tab of the web-interface (along with the Pr_fmax_ estimates) present a handy, first-hand tool for the targeted design of protein interfaces (or, dockable peptides), helping investigators to pinpoint structural defects, irregularity and sub-optimality – which would, in turn, aid in the subsequent re-design. While the current version of EnCPdock is built to handle binary complexes alone, we plan to extend the web-server in the future enabled with the functionality to deal with higher order protein assemblies.

**Figure 9.**
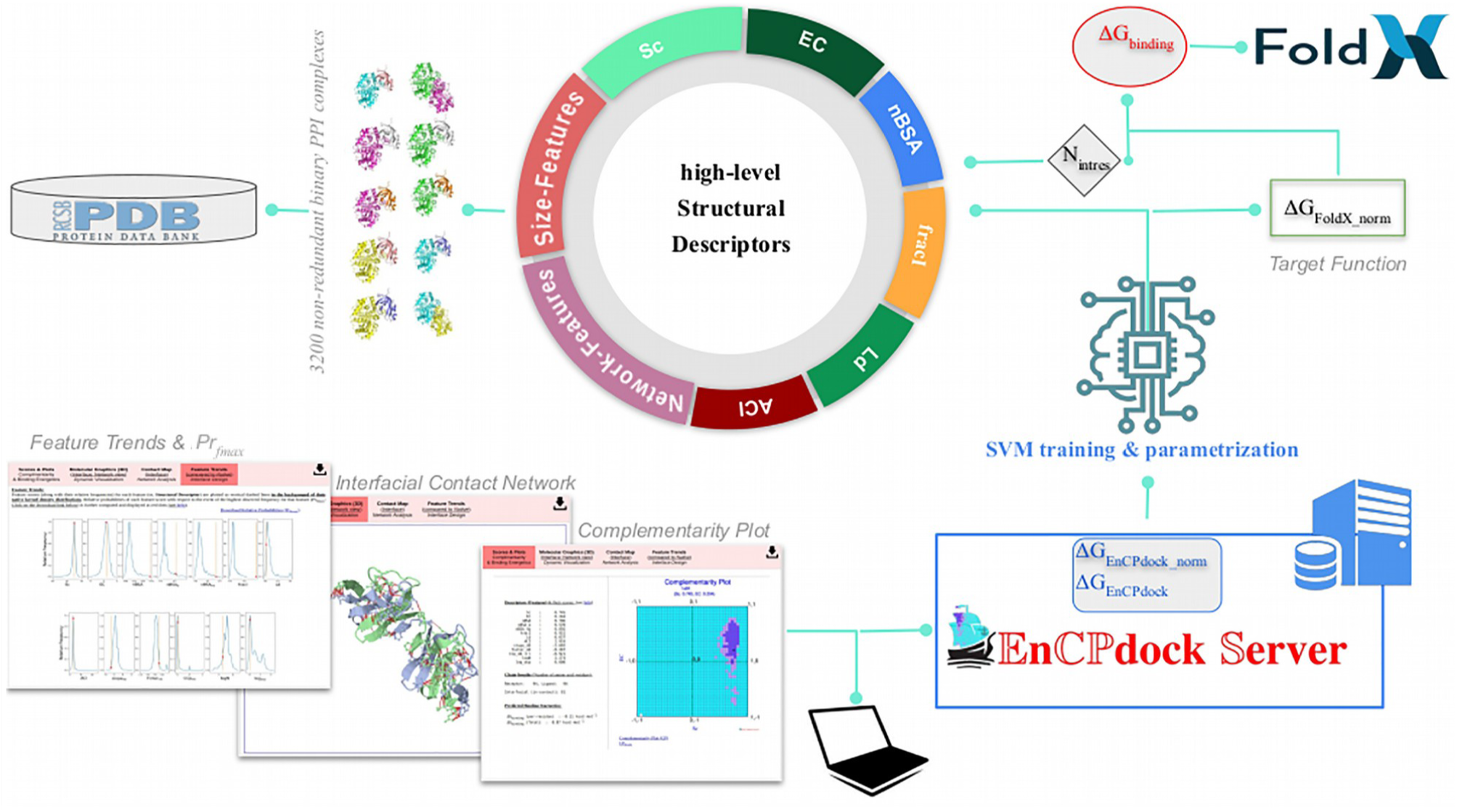
Schematic diagram of EnCPdock web server workflow. The high level structural descriptors used as input parameters are as follows: shape and electrostatic complementarities (Sc, EC); normalized buried surface areas (nBSA, nBSA_P_ & nBSA_np_) and fractional interfacial content (fracI); link density (Ld); average contact intensity (ACI); Network (degree distribution) based Features (slope_dd_, Yinter_dd_, CCp_dd_); Size-Features (logN, log_asp_). While the normalized FoldX – derived ΔGG_binding_ was used as target function. The output showcase ‘Complementarity plot along with Feature information’, ‘Interfacial contact network’ and ‘Feature trends combining relative probability (Pr_fmax_) of the obtained feature-score’ in separate tabular format.

## Declarations

### Ethical Approval

Not Applicable

### Author’s Contributions

SB conceptualized the idea, designed the calculations and wrote the main manuscript with help from GB, ND. DM developed the web-server with assistance from SB. ND and PG did the required literature survey. ND wrote the Introduction. GB performed all training, validations and actively participated in drafting the results and discussion. All the authors participated during the revisions, read and approved the final manuscript.

## Supporting information

The revised Supplementary Materials File (PDF)

## Acknowledgment

We convey our sincerest gratitude to Prof. Dhananjay Bhattacharyya (Saha Institute of Nuclear Physics, Kolkata, India, retired) for his time and thoughts on the mater in the course of one extremely helpful discussion during the revision.

## Availability of data and Materials

Relevant tracking information for all entries in all datasets used, can be found in the online Supplementary Material. Over and above this, any specific data that might be required can be made accessible on request.

## Preprint

The manuscript is also available as a pre-print in BiorXiv.

*EnCPdock: a web-interface for direct conjoint comparative analyses of complementarity and binding energetics in inter-protein associations*

Gargi Biswas, Debasish Mukherjee, Nalok Dutta, Prithwi Ghosh, Sankar Basu*

pre-print to be found at BioRXiv (2023), https://doi.org/10.1101/2023.02.26.530084

Publishers: Cold Spring Harbor Laboratory

## Footnotes

1 Dissociation constant

2 (Gibbs) Free energy of binding

3 Molecular Mechanics combined with Poisson–Boltzmann electrostatics and accessible surface area estimates

4 Molecular Mechanics combined with Generalized Born electrostatics and accessible surface area estimates

5 SD: Standard Deviations

6 receptor and ligand each consisting of a single polypeptide chain

