## Supplementary material for "EnCPdock: a web-interface for direct conjoint comparative analyses of complementarity and binding energetics in inter-protein associations": The revised Supplementary Materials File (PDF)

**Dataset S1. Dataset culled and used in training and cross-validation, consisting of 3200 binary PPI complexes.** The comma-separated names consists of the PDB ID and the chain IDs for both receptor and ligand in the format: <PDB ID>_<chain ID of receptor>_<chain ID of ligand>.

1A09_A_B, 1A1A_A_B, 1A30_A_B, 1A3L_L_H, 1A4Y_A_D, 1A5M_B_C, 1AB9_B_C, 1ABO_A_B, 1AFQ_B_C, 1AHT_L_H,
1AJS_A_B, 1AKS_A_B, 1APY_A_C, 1AVW_A_B, 1B19_A_B, 1B2D_A_B, 1BBB_B_D, 1BCU_L_H, 1BPH_A_B, 1C5C_L_H,
1C5O_L_H, 1C7C_A_D, 1C7D_A_D, 1CB7_B_D, 1CDK_A_B, 1CHO_F_G, 1CM4_A_B, 1CPH_A_B, 1CXP_C_D, 1D0D_A_B,
1D4T_A_B, 1D5L_C_D, 1D8D_A_B, 1DAN_L_H, 1DEI_B_D, 1DGW_A_Y, 1DHK_A_B, 1DLF_L_H, 1DOJ_A_B, 1DOW_A_B,
1DPH_A_B, 1DS2_E_I, 1DSF_L_H, 1DTD_A_B, 1DVF_B_D, 1E3A_A_B, 1E9O_A_B, 1EB1_H_L, 1EG9_A_B, 1EGP_A_B,
1ELW_A_B, 1EO9_A_B, 1EPT_B_C, 1EUV_A_B, 1F2S_E_I, 1F2T_A_B, 1F58_L_H, 1F60_A_B, 1F7Z_A_I, 1FGL_A_B,
1FGV_L_H, 1FIP_A_B, 1FKN_A_B, 1FLT_V_W, 1FM0_D_E, 1FMA_D_E, 1FNS_L_H, 1FVU_A_C, 1FWA_B_C, 1FWB_B_C,
1FWC_B_C, 1FWD_B_C, 1FWF_B_C, 1FWG_B_C, 1FWH_B_C, 1FWI_B_C, 1FXH_A_B, 1FY8_E_I, 1FZ1_A_B, 1FZO_A_B,
1G08_B_D, 1G0B_A_B, 1G0V_A_B, 1G72_A_C, 1GCW_A_C, 1GHB_F_G, 1GHV_L_H, 1GHW_L_H, 1GHY_L_H, 1GJ5_L_H,
1GJ8_A_B, 1GJ9_A_B, 1GJA_A_B, 1GJD_A_B, 1GYB_A_B, 1H4I_A_C, 1H9I_E_I, 1HE1_C_D, 1HLE_A_B, 1HTR_P_B,
1HTV_J_L, 1I7Q_A_C, 1I8K_A_B, 1IE7_B_C, 1IRE_A_B, 1IWH_A_B, 1IZB_B_D, 1J34_A_B, 1JCR_A_B, 1JF1_A_B,
1JFQ_L_H, 1JGD_A_B, 1JQ8_A_B, 1JW9_B_D, 1K22_L_H, 1K3U_A_B, 1K7F_A_B, 1K7X_A_B, 1K8Y_A_B, 1KEL_L_H,
1KFB_A_B, 1KFC_A_B, 1KFE_A_B, 1KFJ_A_B, 1KIQ_B_C, 1KIR_B_C, 1KJV_A_B, 1KL3_A_C, 1KL5_A_C, 1KPU_A_B,
1KUJ_E_G, 1LA6_A_B, 1LES_A_C, 1LI1_A_D, 1LK2_A_B, 1LK3_H_I, 1LOB_E_G, 1LOE_A_C, 1LSH_A_B, 1LT3_H_A,
1LT4_H_A, 1M45_A_B, 1M5A_B_D, 1MCT_A_I, 1MCV_A_I, 1MEE_A_I, 1MHY_B_D, 1MJU_L_H, 1MTY_D_E, 1MVU_A_B,
1MZ8_B_D, 1MZC_A_B, 1MZW_A_B, 1N7F_A_B, 1N9G_B_E, 1NF5_B_D, 1NKH_B_D, 1NKZ_C_E, 1NLB_L_H, 1NME_A_B,
1NMM_B_D, 1NQP_B_D, 1NS9_A_B, 1NT1_A_H, 1NVI_D_E, 1O1J_A_D, 1O1N_A_D, 1O1P_A_D, 1O6L_A_C, 1O94_A_B,
1OAQ_H_L, 1OEB_A_B, 1OGT_A_B, 1OLS_A_B, 1OLU_A_B, 1OP9_A_B, 1OPG_L_H, 1OR0_B_D, 1OW3_A_B, 1OX1_A_B,
1P2J_A_I, 1P2K_A_I, 1P5U_A_B, 1P7T_A_B, 1PNK_A_B, 1PPB_L_H, 1Q0Y_L_H, 1Q40_B_D, 1Q8T_A_B, 1Q8U_A_B,
1QBV_L_H, 1QJ1_A_B, 1QQP_2_3, 1QTN_A_B, 1QTX_A_B, 1QXE_B_D, 1QYG_L_H, 1R1P_B_C, 1R1Y_B_D, 1R4P_A_C,
1R8O_A_B, 1R8S_A_E, 1RBC_S_A, 1RBD_S_A, 1RBE_S_A, 1RBF_S_A, 1RBG_S_A, 1RBH_S_A, 1RBI_S_A, 1RI8_A_B,
1RIU_L_H, 1RJK_A_C, 1RKG_A_C, 1RXZ_A_B, 1RZF_L_H, 1S4V_A_B, 1S63_A_B, 1S6V_A_C, 1S7Q_A_B, 1S7S_A_B,
1S9D_A_E, 1SB2_A_B, 1SBS_H_L, 1SCJ_A_B, 1SEM_A_B, 1SEQ_L_H, 1SFQ_B_E, 1SGP_E_I, 1SGQ_E_I, 1SGR_E_I,
1SHH_B_E, 1SHR_B_D, 1SLE_B_D, 1SLU_A_B, 1SLW_A_B, 1SQ2_L_N, 1SR4_A_B, 1SSA_A_B, 1SSB_A_B, 1SSC_A_B,
1SVD_A_M, 1T0H_A_B, 1T1Y_A_B, 1T1Z_A_B, 1T3F_A_B, 1T3Q_B_E, 1T44_G_A, 1T61_D_E, 1T6V_L_M, 1TA3_A_B,
1TA6_A_B, 1TAW_A_B, 1TJG_L_H, 1TN6_A_B, 1TOM_L_H, 1TSQ_A_B, 1TW6_A_B, 1TZY_A_G, 1U5B_A_B, 1UAC_H_Y,
1UBH_S_L, 1UBJ_S_L, 1UBK_S_L, 1UBL_S_L, 1UBM_S_L, 1UBO_S_L, 1UBP_B_C, 1UBR_S_L, 1UBS_A_B, 1UBT_S_L,
1UBU_S_L, 1UC4_A_L, 1UGP_A_B, 1UGQ_A_B, 1UGR_A_B, 1UGS_A_B, 1UGX_A_B, 1UKM_A_B, 1UKV_G_Y, 1UMD_A_C,
1USP_A_B, 1UVQ_A_B, 1UXW_A_B, 1UZ9_A_B, 1V02_E_F, 1V11_A_B, 1V16_A_B, 1V1M_A_B, 1V1R_A_B, 1V1T_A_B,
1V5I_A_B, 1VFA_A_B, 1VGC_B_C, 1VGE_L_H, 1VPP_V_W, 1VRA_A_B, 1VRK_A_B, 1W7J_A_B, 1W7X_H_L, 1W9Q_A_B,
1WA5_B_C, 1WBJ_A_B, 1WCI_A_B, 1WDC_B_C, 1WDD_A_E, 1WHS_A_B, 1WHT_A_B, 1WQJ_B_I, 1WUH_S_L, 1WUI_S_L,
1WUJ_S_L, 1WUK_S_L, 1WUL_S_L, 1WVM_A_B, 1WX4_A_B, 1WXC_A_B, 1X7W_A_B, 1X7Y_A_B, 1X7Z_A_B, 1X80_A_B,
1XG2_A_B, 1XH3_A_B, 1XH6_A_B, 1XH9_A_B, 1XR9_A_B, 1XU5_A_B, 1XVG_A_B, 1Y43_A_B, 1Y4H_A_B, 1Y4Z_A_B,
1Y5I_A_B, 1YAG_A_G, 1YC5_A_B, 1YDI_A_B, 1YEF_L_H, 1YEG_L_H, 1YGC_H_L, 1YKT_A_B, 1YLC_A_B, 1YLD_A_B,
1YMT_A_B, 1YPE_L_H, 1YPG_L_H, 1YPJ_L_H, 1YPL_L_H, 1YPM_L_H, 1YRK_A_B, 1YRO_B_D, 1YU6_A_B, 1YVQ_B_D,
1YVT_A_B, 1YWO_A_P, 1Z3E_A_B, 1Z3L_S_E, 1Z3M_S_E, 1Z3P_S_E, 1Z7K_A_B, 1Z7X_W_Y, 1Z8I_A_B, 1ZAN_L_H,
1ZBA_2_3, 1ZGX_A_B, 1ZLH_A_B, 1ZNV_B_D, 1ZUZ_A_B, 1ZXI_B_E, 2A0Q_B_D, 2A2Q_H_T, 2A56_B_D, 2A5D_A_B,
2A78_A_B, 2AD6_A_C, 2AD7_A_C, 2AD8_A_C, 2ADF_H_L, 2AER_H_T, 2AHJ_B_D, 2AHK_A_B, 2AJU_L_H, 2AKA_A_B,
2AKR_A_C, 2AL2_A_B, 2AOI_A_B, 2AOJ_A_B, 2AQ2_A_B, 2ARQ_A_P, 2ARR_A_P, 2B1X_C_E, 2BBA_A_P, 2BBK_H_J,
2BEU_A_B, 2BEV_A_B, 2BEW_A_B, 2BFB_A_B, 2BFC_A_B, 2BFD_A_B, 2BFE_A_B, 2BFF_A_B, 2BGR_A_B, 2BLF_A_B,
2BN1_A_B, 2BN3_A_B, 2BPB_A_B, 2BQZ_A_E, 2BRR_H_Y, 2BUM_A_B, 2BUQ_A_B, 2BUR_A_B, 2BUT_A_B, 2BUU_A_B,
2BUV_A_B, 2BUW_A_B, 2BUX_A_B, 2BUY_A_B, 2BUZ_A_B, 2BV0_A_B, 2BVP_A_B, 2BZ6_H_L, 2C8W_A_B, 2C9W_A_B,
2C9X_A_B, 2CA3_A_B, 2CDR_A_B, 2CF8_H_L, 2CF9_H_L, 2CIK_A_B, 2CKL_A_B, 2CLK_A_B, 2CN0_H_L, 2CNK_A_B,
2CNL_A_B, 2CO6_A_B, 2CWG_A_B, 2CYZ_A_B, 2CZ0_A_B, 2CZ1_A_B, 2CZ6_A_B, 2CZ7_A_B, 2D0O_A_C, 2D0Q_A_B,
2D5X_A_B, 2D60_B_D, 2D7T_H_L, 2DBW_A_C, 2DBX_A_C, 2DE5_B_C, 2DE6_B_C, 2DFX_E_I, 2DG5_A_C, 2DJF_A_B,
2DKO_A_B, 2DLF_L_H, 2DN1_A_B, 2DN3_A_B, 2DQC_H_Y, 2DQD_H_Y, 2DQE_H_Y, 2DQJ_H_Y, 2DQT_L_H, 2DRK_A_B,
2E0X_A_C, 2E27_L_H, 2E4M_A_B, 2EEO_A_B, 2ESL_E_F, 2F3Y_A_B, 2F4M_A_B, 2F9I_A_C, 2FBJ_L_H, 2FCW_A_B,
2FHZ_A_B, 2FIV_A_B, 2FLU_X_P, 2FMK_A_B, 2FP7_A_B, 2FVJ_A_B, 2FYC_B_D, 2G2U_A_B, 2G2W_A_B, 2G4M_A_B,
2G81_E_I, 2GA4_A_F, 2GFC_A_I, 2GGV_A_B, 2GL9_C_D, 2GMT_B_C, 2GNG_A_I, 2GPO_A_C, 2GZE_A_B, 2GZF_A_B,
2GZG_A_B, 2H5I_A_B, 2H6F_A_B, 2H6G_A_B, 2H6H_A_B, 2H6K_A_B, 2H6N_A_B, 2H6Q_A_B, 2HBE_A_B, 2HMJ_A_B,
2HMK_A_B, 2HML_A_B, 2HMM_A_B, 2HMN_A_B, 2HMO_A_B, 2HO2_A_B, 2HPE_A_B, 2HPZ_A_B, 2HY5_A_B, 2I3H_A_B,
2IEJ_A_B, 2INC_A_B, 2IPT_L_H, 2IUC_A_B, 2IVF_A_B, 2IZX_A_B, 2J12_A_B, 2J2U_A_B, 2J4I_A_B, 2JDO_A_C,
2JH0_C_D, 2KAU_B_C, 2LAL_A_C, 2LTN_A_C, 2NL9_A_B, 2NS1_A_B, 2NU0_E_I, 2NU1_E_I, 2NU2_E_I, 2NU3_E_I,
2NU4_E_I, 2NXY_A_D, 2NY2_A_D, 2O4R_A_C, 2O5G_A_B, 2O88_A_B, 2O9Q_A_C, 2O9V_A_B, 2OD3_A_B, 2OM1_R_T,
2OMU_A_B, 2OMX_A_B, 2OVH_A_B, 2OZL_A_C, 2P1M_A_B, 2P1O_A_B, 2P1Q_A_B, 2P1T_A_B, 2P3T_A_B, 2P3U_A_B,

2P42_A_C, 2P43_A_B, 2P44_A_B, 2P45_A_B, 2P49_A_B, 2P58_A_C, 2P5E_A_E, 2P8O_B_C, 2PAV_A_P, 2PBD_A_P, 2PGB_A_B, 2PGQ_A_B, 2PLX_A_B, 2POY_A_B, 2PQK_A_B, 2PTT_A_B, 2PUX_A_B, 2Q0N_A_B, 2Q5W_E_D, 2Q76_A_C, 2Q7I_A_B, 2Q7J_A_B, 2Q7K_A_B, 2Q7L_A_B, 2Q7Y_A_C, 2QAC_A_T, 2QDY_A_B, 2QFA_A_B, 2QKH_B_A, 2QQC_J_L, 2QTW_A_B, 2QWO_A_B, 2R7G_A_C, 2RAO_B_D, 2RH9_A_B, 2RIV_A_B, 2RLN_S_E, 2RMC_E_G, 2SEC_E_I, 2TCI_B_D, 2UBP_B_C, 2UUF_A_B, 2UUJ_A_B, 2UUK_A_B, 2UVX_A_I, 2UW8_A_I, 2UWL_A_B, 2UYN_A_B, 2UYZ_A_B, 2UZI_H_R, 2V17_H_L, 2V1T_A_B, 2V3H_H_L, 2V3O_H_L, 2V3S_A_B, 2V52_B_M, 2V6X_A_B, 2V8C_A_C, 2V9T_A_B, 2VGX_A_B, 2VH0_A_B, 2VH6_A_B, 2VLN_A_B, 2VLO_A_B, 2VLP_A_B, 2VN6_A_B, 2VNF_A_C, 2VO0_A_I, 2VO3_A_I, 2VO6_A_I, 2VO7_A_I, 2VOG_A_B, 2VOH_A_B, 2VPB_A_B, 2VSM_A_B, 2VT1_A_B, 2VU8_E_I, 2VWF_A_B, 2VXQ_H_L, 2VXT_H_L, 2VXV_H_L, 2VZG_A_B, 2W60_A_B, 2WBW_A_B, 2WEL_A_D, 2WJM_C_M, 2WJN_C_M, 2WKO_A_F, 2WP3_O_T, 2WPV_A_G, 2WRU_A_B, 2WS1_A_B, 2WY7_A_Q, 2WY8_A_Q, 2WYC_A_B, 2WYD_A_B, 2WYE_A_B, 2WYG_A_B, 2X39_A_C, 2X4W_A_B, 2X70_A_D, 2X7R_C_E, 2XFG_A_B, 2XFX_A_B, 2XGY_A_B, 2XPP_A_B, 2XRW_A_B, 2XT1_A_B, 2XU7_A_B, 2XWT_A_C, 2XXM_A_B, 2XXN_A_B, 2XYH_A_B, 2XYI_A_B, 2XYP_A_B, 2Y1L_A_F, 2Y1N_A_C, 2Y3N_A_C, 2Y8N_A_C, 2Y9Q_A_B, 2YLE_A_B, 2YPK_A_B, 2YPV_A_L, 2YQ6_A_B, 2YQ7_A_B, 2YVJ_A_P, 2Z2Y_A_C, 2Z30_A_B, 2Z56_A_B, 2Z57_A_B, 2Z58_A_B, 2Z8K_A_C, 2ZC9_L_H, 2ZCF_A_B, 2ZDA_L_H, 2ZDV_L_H, 2ZFD_A_B, 2ZFF_L_H, 2ZFQ_L_H, 2ZFR_L_H, 2ZG0_L_H, 2ZGB_L_H, 2ZHF_L_H, 2ZHQ_L_H, 2ZI2_L_H, 2ZIQ_L_H, 2ZLT_A_B, 2ZLU_A_B, 2ZMY_A_B, 2ZNK_L_H, 2ZO3_L_H, 2ZON_B_C, 2ZPB_A_B, 2ZPE_A_B, 2ZPF_A_B, 2ZPG_A_B, 2ZPH_A_B, 2ZPI_A_B, 2ZPK_L_H, 2ZS0_B_C, 2ZS1_B_C, 2ZWD_A_B, 3A3O_A_B, 3A3P_A_B, 3A4U_A_B, 3A67_H_Y, 3A6B_H_Y, 3A6C_H_Y, 3A8G_A_B, 3A8H_A_B, 3A8L_A_B, 3A8M_A_B, 3A8O_A_B, 3AA6_A_B, 3ABD_A_B, 3AMA_A_B, 3AON_A_B, 3AVR_A_B, 3AWV_A_B, 3AWX_A_B, 3AWY_A_B, 3AWZ_A_B, 3AXK_A_B, 3AXM_F_H, 3AYU_A_B, 3B3I_A_B, 3B6S_A_B, 3B7V_A_B, 3B80_A_B, 3BDG_A_B, 3BEI_A_B, 3BFQ_G_F, 3BH9_A_B, 3BKJ_L_H, 3BP4_A_B, 3BP5_A_B, 3BP6_A_B, 3BP7_A_B, 3BRL_A_C, 3BRP_A_B, 3BV9_A_B, 3BW9_A_B, 3BWA_A_B, 3BXN_A_B, 3BY4_A_B, 3BYA_A_B, 3BZX_A_B, 3BZY_A_B, 3BZZ_A_B, 3C4M_A_B, 3C4O_A_B, 3C4P_A_B, 3C9A_A_B, 3C9N_A_B, 3CBJ_A_B, 3CFC_H_L, 3CI5_A_G, 3CIP_A_G, 3CLR_C_D, 3CLS_C_D, 3CLU_C_D, 3CNQ_P_S, 3D18_A_B, 3D1K_A_B, 3D25_A_B, 3D4U_A_B, 3D6M_A_B, 3D9A_L_H, 3DA9_A_B, 3DAB_A_G, 3DDC_A_B, 3DDQ_A_D, 3DGP_A_B, 3DHI_A_B, 3DHK_L_H, 3DLQ_I_R, 3DNE_A_I, 3DRA_A_B, 3DX7_A_B, 3DZ2_B_A, 3DZ7_B_A, 3E1Z_A_B, 3E37_A_B, 3E94_A_B, 3ECB_A_B, 3EG1_A_B, 3EGV_A_B, 3EQ0_L_H, 3EQS_A_B, 3ET3_A_P, 3F1N_A_B, 3F1O_A_B, 3F62_A_B, 3F6Q_A_B, 3F75_A_P, 3F9X_C_D, 3FDL_A_B, 3FDO_A_B, 3FGR_A_B, 3FJQ_E_I, 3FJU_A_B, 3FT3_A_B, 3FT4_A_B, 3FU7_A_B, 3G08_A_B, 3G2T_A_B, 3G5Y_A_B, 3G9K_L_D, 3GE3_A_B, 3GIC_A_B, 3GIV_A_D, 3GJ7_A_C, 3GJ8_A_C, 3GKV_A_B, 3GML_A_B, 3GMM_A_B, 3GMN_A_B, 3GMO_A_B, 3GMP_A_B, 3GMQ_A_B, 3GMR_A_B, 3GO1_L_H, 3GP2_A_B, 3GSO_A_B, 3GSR_A_B, 3H0W_B_A, 3H1Z_A_P, 3H3G_A_B, 3H6P_B_C, 3H6R_A_B, 3H7P_A_B, 3H7W_A_B, 3H82_B_A, 3H8K_A_B, 3HC3_H_L, 3HC4_H_L, 3HE5_B_F, 3HHS_A_B, 3HK3_A_B, 3HNS_H_L, 3HNT_H_L, 3HNV_H_L, 3HS8_A_P, 3HYU_A_B, 3I3Z_A_B, 3I5J_A_B, 3I75_A_B, 3I9G_H_L, 3IG6_B_D, 3ILG_B_D, 3IU4_H_L, 3IXE_A_B, 3JXT_A_B, 3JZO_A_P, 3JZP_A_P, 3K74_A_B, 3K9O_A_B, 3KDF_D_C, 3KJ0_A_B, 3KJ1_A_B, 3KJF_A_B, 3KJN_A_B, 3KJQ_A_B, 3KL6_A_B, 3KLA_A_D, 3KLD_A_B, 3KMR_A_C, 3KMW_A_B, 3KNB_A_B, 3KPE_A_B, 3KPN_A_B, 3KPP_A_B, 3KRA_A_D, 3KTA_A_B, 3KUC_A_B, 3KUJ_A_B, 3KVS_A_B, 3KXC_A_C, 3KYJ_A_B, 3L0F_A_B, 3L1O_H_L, 3L3D_A_B, 3L3I_A_B, 3L3X_A_B, 3L3Z_A_B, 3L51_A_B, 3L91_A_B, 3L94_A_B, 3LCN_A_B, 3LEX_L_B, 3LEY_H_L, 3LIZ_A_L, 3LKO_A_B, 3LKQ_A_B, 3LL8_A_C, 3LN4_A_B, 3LN5_A_B, 3LU9_B_E, 3LXR_A_F, 3M18_A_B, 3M1I_A_C, 3M61_U_P, 3M7F_A_B, 3M7Q_A_B, 3MC0_B_D, 3MCB_A_B, 3MEZ_B_D, 3MHP_A_B, 3MHS_A_C, 3ML1_A_B, 3MLY_H_I, 3MM6_A_D, 3MN5_A_S, 3MNE_A_B, 3MNP_A_B, 3MNZ_A_B, 3MO8_A_B, 3MRC_A_B, 3MRE_A_B, 3MRG_A_B, 3MRJ_A_B, 3MRK_A_B, 3MRM_A_B, 3MRR_A_B, 3MSX_A_B, 3MTH_B_D, 3MXN_A_B, 3MXV_L_H, 3N06_A_B, 3N1M_B_C, 3N4I_A_B, 3N9G_H_L, 3NCE_A_B, 3NDD_A_B, 3NHE_A_B, 3NIK_B_D, 3NIM_A_B, 3NL7_A_B, 3NPS_A_B, 3NRZ_C_L, 3NV0_A_B, 3NVV_C_L, 3NVW_C_L, 3NVY_C_L, 3NY7_A_B, 3NZH_L_H, 3O5A_A_B, 3O8Q_A_B, 3OD5_A_B, 3OKL_A_B, 3OR1_A_D, 3OTS_A_B, 3OVV_A_B, 3OW3_A_B, 3OWP_A_B, 3OXR_A_B, 3OXS_A_B, 3P0Y_H_L, 3P17_L_H, 3P73_A_B, 3P77_A_B, 3P8F_A_I, 3P92_A_E, 3P95_A_E, 3PDH_A_D, 3PEL_A_B, 3PHX_A_B, 3PLF_B_D, 3PLU_A_B, 3PLX_A_B, 3POO_A_B, 3PR2_A_B, 3PTH_A_B, 3PWU_A_B, 3Q3K_A_B, 3Q3N_A_B, 3Q3O_A_B, 3Q6G_H_I, 3Q7A_A_B, 3QHF_H_L, 3QJB_A_B, 3QKL_A_C, 3QN1_A_B, 3QN7_A_B, 3QQ8_A_B, 3QSK_A_B, 3QTO_L_H, 3QTV_L_H, 3QWC_L_H, 3QWO_H_A, 3QX5_L_H, 3R24_A_B, 3R3G_A_B, 3RGW_L_S, 3RL1_A_B, 3RLW_L_H, 3RLY_L_H, 3RM0_L_H, 3RM2_L_H, 3RML_L_H, 3RMM_L_H, 3RMN_L_H, 3RMO_L_H, 3RNK_A_B, 3RNQ_B_A, 3RUB_L_S, 3RUI_A_B, 3RVV_A_D, 3S43_A_B, 3S7H_A_B, 3S7K_B_D, 3S8F_A_B, 3S8G_A_B, 3S90_A_B, 3S96_A_C, 3SBT_A_B, 3SDE_A_B, 3SE8_G_H, 3SE9_G_H, 3SFV_A_B, 3SHA_L_H, 3SHC_L_H, 3SHG_A_B, 3SI3_L_H, 3SI4_L_H, 3SIC_E_I, 3SJH_A_B, 3SKM_A_B, 3SO6_A_Q, 3SOB_B_H, 3SPV_A_B, 3SRB_A_B, 3SRC_A_B, 3SRI_A_B, 3SRN_A_B, 3SUI_A_B, 3SV2_L_H, 3SXU_A_B, 3SY0_A_B, 3T1F_A_B, 3T4Y_A_B, 3T5F_L_H, 3T5G_A_B, 3T65_B_A, 3T6D_C_M, 3TGI_E_I, 3TH2_H_T, 3TID_A_B, 3TJ5_A_B, 3TLG_A_B, 3TMP_C_E, 3TU3_A_B, 3TV3_L_H, 3TVJ_A_B, 3TWC_L_H, 3TWU_A_B, 3U0W_H_L, 3U1J_B_E, 3U23_A_B, 3U43_A_B, 3U4N_A_B, 3U52_A_B, 3U69_L_H, 3U8O_L_H, 3U8R_L_H, 3U8T_L_H, 3U8X_A_C, 3U98_L_H, 3U9A_L_H, 3UA7_A_B, 3UBP_B_C, 3UBU_A_B, 3UJI_L_H, 3UJJ_L_H, 3UL0_B_C, 3UL1_B_A, 3UL4_A_B, 3ULR_A_B, 3UO1_H_L, 3UOA_B_C, 3UOU_A_B, 3UP0_A_B, 3UP3_A_P, 3UTQ_A_B, 3UVK_A_B, 3UVM_A_B, 3UVW_A_B, 3UWJ_L_H, 3UXG_A_B, 3UYO_A_D, 3UYP_A_B, 3UYR_H_L, 3UZQ_A_B, 3V0W_L_H, 3V2X_A_B, 3V30_A_B, 3V31_A_B, 3V3B_A_B, 3V49_A_B, 3V4A_A_B, 3V52_H_L, 3V6C_A_B, 3VA4_A_B, 3VB7_A_B, 3VCL_A_B, 3VFG_L_H, 3VFM_A_B, 3VFR_A_B, 3VFS_A_B, 3VFT_A_B, 3VFU_A_B, 3VFV_A_B, 3VGX_C_D, 3VMG_B_C, 3VMH_B_C, 3VQH_A_B, 3VRD_A_B, 3VRF_A_B, 3VRG_A_B, 3VRI_A_B, 3VRJ_A_B, 3VT7_A_C, 3VTC_A_B, 3VTP_C_D, 3VXE_L_H, 3VXN_A_B, 3VYH_A_B, 3VZ9_B_D, 3W0H_A_C, 3W19_C_D, 3W7Z_B_D, 3WA5_A_B, 3WHQ_A_B, 3WHR_A_B, 3WHS_A_B, 3WHT_A_B, 3WIF_A_B, 3WL9_A_B, 3WLB_A_B, 3WMI_A_B, 3WMJ_A_B, 3WOO_A_B, 3WOP_A_B, 3WOQ_A_B, 3WQB_A_B, 3WT5_A_C, 3WVD_A_B, 3WVE_A_B, 3WWN_A_B, 3X12_A_B, 3X13_A_B, 3X14_A_B, 3X24_A_B, 3X25_A_B, 3X26_A_B, 3X28_A_B, 3X2V_A_S, 3X2W_A_S, 3ZCB_A_B, 3ZE6_A_B, 3ZG9_A_B, 3ZI3_A_B, 3ZIN_A_B, 3ZKW_A_B, 3ZN6_A_B, 3ZNZ_A_B, 3ZO0_A_B, 3ZO1_A_I, 3ZO2_A_I, 3ZO4_A_I, 3ZOQ_A_C, 3ZQH_A_C, 3ZRZ_A_B, 4A1U_A_B, 4A5U_A_B, 4A7E_A_B, 4A8X_A_C, 4ABI_A_B, 4AC7_B_C, 4AE4_A_B, 4AFS_A_C, 4AFU_A_B, 4AJY_B_V, 4AL8_H_L, 4ALA_H_L, 4AOM_A_T, 4AQA_A_B, 4AT7_A_B, 4B0M_A_M, 4B1U_B_M, 4B1W_B_M, 4B1X_B_M, 4B1Y_B_M, 4B4N_A_B, 4B4S_A_B, 4BA1_A_B, 4BH4_A_B, 4BI8_A_B, 4BL7_A_B, 4BQK_A_B, 4BTH_A_B, 4BU0_A_C, 4BVY_A_B, 4BWC_A_B, 4BWL_A_D, 4C33_A_I, 4C34_A_I, 4C37_A_I, 4C4K_O_T, 4C4P_A_B, 4CBU_A_G, 4CEU_B_C, 4CEX_B_C, 4CFT_A_B, 4CJ0_A_B, 4CJ1_A_B, 4CJ2_A_B, 4CMH_A_C, 4CRU_A_B, 4CRW_A_B, 4CRY_B_G, 4CRZ_A_B, 4CSR_A_B, 4CT7_A_B, 4CU1_A_B, 4CXF_A_B, 4CXN_A_B, 4CZX_A_B, 4D07_A_B, 4D0K_A_C, 4D7Z_A_B, 4DEX_A_B, 4DFX_E_I, 4DG0_E_I, 4DG2_E_I, 4DG3_E_A, 4DGV_H_L, 4DGY_H_L, 4DH1_A_I, 4DH7_A_I, 4DHI_B_D, 4DJC_A_B, 4DKA_A_B, 4DLQ_A_B, 4DOR_A_B, 4DRI_A_B, 4DRJ_A_B, 4DS1_A_C, 4DT7_B_D, 4DXA_A_B, 4DZB_A_B, 4E3B_A_B, 4EB2_A_B, 4EGC_A_B, 4EIK_A_B, 4EIS_A_B, 4EP8_C_B, 4EPB_C_B, 4EPD_C_B, 4ERY_A_D, 4ESG_A_B, 4EUK_A_B, 4EWR_A_C, 4F0H_A_B, 4F0Z_A_B, 4F14_A_B, 4F27_A_Q, 4F57_L_H, 4F6T_B_A, 4F7P_A_B, 4FBJ_A_B, 4FQ2_H_L, 4FQL_H_L, 4FZE_L_H, 4G27_B_R, 4G28_B_R, 4G35_A_B, 4G43_A_D, 4G5Z_H_L, 4G6K_H_L, 4G7S_A_B, 4G91_B_C, 4G9D_A_B, 4GAG_H_L, 4GBC_B_D, 4GBN_B_D, 4GDX_A_B, 4GED_A_B, 4GF3_A_B, 4GFT_A_B, 4GHI_A_B, 4GI3_A_C, 4GLY_A_B, 4GN4_B_A, 4GQ6_A_B, 4GS9_A_B, 4GUT_A_B, 4GVD_A_B, 4GXB_A_B, 4H20_L_H,

4H2L_A_B, 4H3K_A_D, 4H4F_A_B, 4H5S_A_B, 4H8W_G_L, 4H9N_A_C, 4H9Q_A_C, 4HAT_A_C, 4HAU_A_C, 4HAV_A_C, 4HAW_A_C, 4HAZ_A_C, 4HB2_A_C, 4HDK_A_B, 4HDM_A_B, 4HEP_A_G, 4HGC_A_I, 4HGW_A_B, 4HI8_A_B, 4HI9_A_B, 4HJL_A_B, 4HKV_A_B, 4HM1_A_B, 4HM2_A_B, 4HM3_A_B, 4HM4_A_B, 4HM5_A_B, 4HM6_A_B, 4HM7_A_B, 4HM8_A_B, 4HN4_A_B, 4HOM_A_B, 4HPO_H_L, 4HPU_E_I, 4HPY_H_L, 4HRT_A_B, 4HST_A_B, 4HT3_A_B, 4HTF_A_B, 4HVU_A_B, 4HVV_A_B, 4HVW_A_B, 4HX1_A_B, 4I13_A_B, 4I1N_A_B, 4I4W_A_B, 4I5Y_A_B, 4I5Z_A_B, 4I77_H_L, 4IAD_A_S, 4IAI_A_S, 4IAK_A_S, 4IAY_A_S, 4IB1_A_S, 4IBL_H_T, 4IE9_A_I, 4IFL_X_P, 4IG7_A_B, 4ILH_A_B, 4ISH_H_L, 4ISV_A_B, 4IU3_A_B, 4IUB_L_S, 4IUC_L_S, 4IUD_L_S, 4IUM_A_B, 4J2Y_A_B, 4J32_A_B, 4J46_A_C, 4J6R_G_H, 4J82_A_B, 4J84_A_B, 4J8G_A_B, 4J8S_A_B, 4J9C_A_B, 4J9H_A_B, 4J9Y_B_R, 4J9Z_B_R, 4JB8_A_P, 4JBN_A_B, 4JIF_A_B, 4JK5_A_B, 4JML_A_E, 4JN1_H_L, 4JN6_A_C, 4JQU_A_B, 4JQV_A_C, 4JQX_A_C, 4JS0_A_B, 4JW2_A_B, 4JYU_H_L, 4JZE_H_L, 4JZF_H_L, 4JZZ_A_R, 4K12_A_B, 4K1E_A_B, 4K1R_A_C, 4K2F_A_B, 4K5B_A_C, 4K6Y_A_B, 4K72_A_B, 4K78_A_B, 4K7L_A_B, 4K8Y_A_B, 4K90_A_B, 4KA2_A_R, 4KDL_A_B, 4KEL_A_B, 4KFB_A_B, 4KML_A_B, 4KPU_A_B, 4KQ3_L_H, 4KT3_A_B, 4KT6_A_C, 4KTE_H_L, 4KTS_A_B, 4KTU_A_B, 4KV1_A_B, 4KXQ_A_B, 4KZL_A_B, 4L2I_A_B, 4L32_A_B, 4L4V_A_C, 4L6T_A_C, 4LCI_L_H, 4LCY_A_F, 4LDT_A_C, 4LGR_A_B, 4LHJ_A_B, 4LJO_A_B, 4LK9_A_B, 4LKA_A_B, 4LKF_A_B, 4LKX_A_B, 4LLQ_A_B, 4LMS_B_D, 4LN2_A_B, 4LNB_A_B, 4LNP_A_B, 4LNR_A_B, 4LOY_H_L, 4LRI_P_C, 4LRS_A_B, 4LV5_A_B, 4LV8_A_B, 4LX2_A_B, 4LX3_A_B, 4LXB_H_L, 4LZF_A_B, 4LZX_A_B, 4M0W_A_B, 4M1D_H_I, 4M3K_A_B, 4M43_L_H, 4M5S_A_B, 4M6M_L_H, 4M6N_L_H, 4MD4_A_B, 4MNV_A_B, 4MNW_A_B, 4MNX_A_B, 4MQJ_B_H, 4MRT_C_A, 4MS4_A_B, 4MS8_C_D, 4MVI_A_B, 4MZ6_C_E, 4N0H_A_B, 4N0Y_H_L, 4N1C_A_C, 4N39_A_B, 4N9O_A_B, 4NBC_B_C, 4NBF_A_B, 4NBG_B_C, 4NBX_A_B, 4NDR_B_A, 4NI2_A_B, 4NIB_A_B, 4NMR_A_B, 4NMS_A_B, 4NMX_A_B, 4NR3_A_B, 4NRG_A_B, 4NT6_A_B, 4NUG_L_H, 4NUJ_A_B, 4NUT_A_B, 4NZR_H_M, 4O21_A_S, 4O22_A_S, 4O2C_A_B, 4O2E_A_D, 4O2F_A_D, 4O36_A_B, 4O37_A_B, 4O4B_A_B, 4O4Y_L_H, 4O5L_L_H, 4O8X_A_B, 4O8Y_A_B, 4OB0_A_B, 4OB1_A_B, 4OB2_A_B, 4OB3_A_B, 4OBD_B_C, 4OCR_L_H, 4OD8_B_A, 4ODC_A_B, 4ODN_A_B, 4ODT_H_L, 4OEL_A_B, 4OEM_A_B, 4OIC_A_B, 4OJF_H_L, 4OKF_A_B, 4OMF_A_B, 4ONF_H_L, 4ONL_A_B, 4ONM_A_B, 4ONN_A_B, 4OS4_A_B, 4OS6_A_B, 4OS7_A_B, 4OWT_A_B, 4P3C_H_L, 4PAS_A_B, 4PID_A_B, 4PJ2_B_C, 4PKG_A_G, 4PL8_A_B, 4PLD_A_B, 4POU_A_B, 4POZ_C_D, 4PRE_A_B, 4PRN_A_B, 4PTT_A_B, 4PTU_A_B, 4PZ3_A_B, 4PZ5_A_B, 4Q4Y_1_2, 4Q57_A_B, 4Q5E_A_C, 4Q5H_A_C, 4Q6F_A_B, 4QE6_A_B, 4QH7_B_F, 4QH8_A_B, 4QLP_A_B, 4QO1_A_B, 4QRQ_A_B, 4QRS_A_B, 4QRU_A_B, 4QVF_A_B, 4QXT_A_B, 4QY8_A_B, 4QYO_A_B, 4R1D_A_B, 4R1E_A_B, 4R3S_A_B, 4R6P_E_G, 4R6R_E_G, 4R90_L_H, 4REY_A_B, 4RGO_S_L, 4RKJ_A_B, 4RKO_B_A, 4RLJ_A_B, 4RTV_A_B, 4RTY_A_B, 4RWF_A_B, 4RWV_A_B, 4RXZ_A_B, 4S1I_A_B, 4SRN_A_B, 4TOY_H_L, 4TPR_L_H, 4TQ1_A_B, 4TQE_L_H, 4TR1_A_B, 4TSB_H_L, 4TSH_A_B, 4TTT_L_S, 4TUL_H_L, 4TXR_A_C, 4U1E_I_G, 4U1H_A_B, 4U1I_A_B, 4U1J_A_B, 4U1M_A_B, 4U1N_A_B, 4U1P_A_B, 4U1S_A_B, 4U2W_B_A, 4U32_X_A, 4U3S_A_B, 4U4P_A_B, 4U5Y_A_D, 4U68_A_B, 4U6X_A_B, 4U6Y_A_B, 4U7E_B_A, 4U7I_A_B, 4U91_A_E, 4U9H_S_L, 4U9I_S_L, 4UAF_B_E, 4UBP_B_C, 4UDW_H_L, 4UE7_H_L, 4UE8_A_B, 4UFD_H_L, 4UFE_H_L, 4UIK_H_L, 4UJ1_A_B, 4UJ9_A_B, 4UJA_A_B, 4UJB_A_B, 4UN2_A_B, 4UQY_A_B, 4UQZ_A_B, 4UT7_H_L, 4UU5_A_B, 4UWX_A_B, 4V0X_A_B, 4V3L_A_B, 4W4O_A_C, 4W6X_A_B, 4W6Y_A_B, 4WB5_A_I, 4WB8_A_I, 4WCY_H_L, 4WEM_A_B, 4WEN_A_B, 4WHO_D_F, 4WI0_A_B, 4WIH_A_B, 4WJK_A_B, 4WJN_A_B, 4WJO_A_B, 4WJQ_A_C, 4WK0_A_B, 4WKS_C_A, 4WKT_C_A, 4WKZ_A_B, 4WLR_A_B, 4WMI_A_D, 4WN2_A_D, 4WND_A_B, 4WNH_A_D, 4WSF_A_B, 4WUK_H_L, 4WVF_A_C, 4WW5_A_B, 4WW7_A_B, 4WW9_A_B, 4WX2_A_B, 4WX4_A_C, 4WY4_C_D, 4WZX_A_E, 4X1C_G_H, 4X28_A_B, 4X2H_A_B, 4X2O_A_B, 4X33_A_B, 4X34_A_B, 4X5W_A_B, 4X6F_A_B, 4XAW_L_H, 4XBE_L_H, 4XC1_L_H, 4XC3_L_H, 4XCF_L_H, 4XHV_A_B, 4XLG_B_A, 4XMP_G_H, 4XO9_A_B, 4XOD_B_A, 4XOJ_A_B, 4XSG_A_B, 4XVS_H_G, 4XW4_A_B, 4XW5_A_B, 4XW6_A_B, 4XYM_A_C, 4Y0Y_E_I, 4Y0Z_E_I, 4Y10_E_I, 4Y11_E_I, 4Y1D_A_B, 4Y29_A_B, 4Y6G_A_B, 4Y71_A_B, 4Y76_A_B, 4Y7A_A_B, 4Y7B_A_B, 4Y7M_C_D, 4YB8_A_C, 4YDL_G_A, 4YEC_A_B, 4YGM_B_A, 4YH8_A_B, 4YHO_H_L, 4YI0_C_A, 4YII_U_A, 4YJE_A_B, 4YK6_B_A, 4YL8_A_B, 4YN3_A_B, 4YNY_C_D, 4YO0_A_B, 4YON_A_B, 4YZ6_A_B, 4YZU_A_B, 4Z09_A_C, 4Z0E_A_C, 4Z0K_A_B, 4Z0X_A_B, 4Z2O_A_P, 4Z68_A_E, 4Z8A_A_B, 4Z9K_A_B, 4Z9W_A_B, 4ZA3_A_B, 4ZAE_A_B, 4ZEQ_A_B, 4ZGM_A_B, 4ZGR_A_B, 4ZHA_A_B, 4ZHY_A_B, 4ZIE_A_C, 4ZKS_U_P, 4ZOX_A_B, 4ZOZ_A_B, 4ZQC_A_B, 4ZTP_L_H, 4ZV0_A_B, 4ZW2_A_B, 4ZYK_L_B, 5A29_A_B, 5A6T_B_C, 5A6W_B_C, 5ABX_A_B, 5ABY_A_C, 5ADO_H_L, 5ADS_A_B, 5AFG_A_B, 5AJJ_A_B, 5ALC_H_L, 5AQB_A_B, 5AQH_A_B, 5AQI_A_C, 5AQT_A_B, 5AQU_A_B, 5AQV_A_B, 5AUO_A_B, 5AWN_H_L, 5AYU_L_H, 5AZ8_A_B, 5B3G_A_B, 5B41_A_C, 5B5B_A_D, 5B5T_A_C, 5B6C_A_B, 5B75_A_B, 5B76_A_B, 5B77_A_B, 5B78_A_B, 5BN3_A_B, 5BN6_G_H, 5BOQ_B_F, 5BTS_A_B, 5BW6_A_B, 5BWD_A_C, 5BX6_A_B, 5BX7_A_B, 5BY8_A_B, 5C0C_A_F, 5C0D_A_B, 5C0E_A_B, 5C0F_A_B, 5C0G_A_B, 5C0I_A_B, 5C0J_A_B, 5C2U_B_A, 5C3F_A_B, 5CEC_A_B, 5CGQ_A_B, 5CRW_A_B, 5CTD_B_C, 5CTT_A_B, 5CYZ_A_C, 5D1K_A_B, 5D1L_A_B, 5D1M_A_B, 5D51_L_S, 5D5K_C_B, 5D94_A_B, 5D9S_A_B, 5DC4_A_B, 5DC9_A_B, 5DCM_A_B, 5DD1_H_L, 5DDH_A_B, 5DF6_A_C, 5DHM_C_D, 5DJT_A_B, 5DQD_L_H, 5DR5_H_L, 5DRN_A_B, 5DT1_H_L, 5E00_A_B, 5E0L_A_C, 5E0Q_A_B, 5E0W_A_B, 5E2V_L_H, 5E5U_A_C, 5E6O_A_B, 5E8M_A_B, 5E95_B_A, 5E97_A_B, 5E9C_A_B, 5EFQ_A_C, 5EGM_A_B, 5EI3_A_B, 5EKF_C_A, 5ELJ_A_B, 5ELQ_A_B, 5EN2_A_B, 5EN9_A_B, 5ENA_A_B, 5EOQ_H_L, 5EPP_A_B, 5EU0_A_B, 5EU3_A_B, 5EU5_A_B, 5EUI_A_B, 5EWI_H_L, 5EY5_B_D, 5EZI_H_L, 5F0E_A_B, 5F1N_A_D, 5F21_A_B, 5F2U_A_B, 5F67_A_B, 5F6L_B_A, 5F7E_H_L, 5F86_A_B, 5F8Z_A_B, 5FB6_A_B, 5FBY_A_B, 5FCF_A_B, 5FCU_H_L, 5FG8_A_B, 5FGB_A_C, 5FHA_H_L, 5FHB_H_L, 5FIW_C_D, 5FMK_A_B, 5FOS_A_C, 5FSD_B_C, 5FVB_W_X, 5FVD_A_C, 5G44_A_C, 5G46_A_C, 5G4H_B_C, 5G4K_A_B, 5G5G_B_C, 5GGQ_L_H, 5GGS_C_D, 5GGV_L_H, 5GIS_H_L, 5GJY_A_B, 5GP7_A_B, 5GPG_A_B, 5GRG_A_B, 5GRQ_A_B, 5GRU_A_H, 5GSB_A_B, 5GTU_A_B, 5GVI_A_B, 5H19_A_B, 5H3J_A_B, 5H5Q_A_B, 5H5R_A_B, 5H5S_A_B, 5H5Z_A_B, 5H8F_A_B, 5H8Q_A_B, 5H94_A_D, 5HBR_A_C, 5HE9_A_E, 5HHP_A_B, 5HKQ_A_I, 5HKY_A_B, 5HKZ_A_B, 5HU3_A_B, 5HUW_C_A, 5HVP_A_B, 5HXG_A_C, 5I18_L_H, 5I1K_L_H, 5I1L_L_H, 5I66_A_B, 5I8C_A_B, 5I8K_L_H, 5I8O_H_L, 5IAA_A_B, 5IB1_A_B, 5IB2_A_B, 5IB3_A_B, 5IB4_A_B, 5IED_A_B, 5IEE_A_B, 5IEH_A_B, 5IEK_A_B, 5IG7_D_J, 5IIB_A_B, 5IL0_A_B, 5IL1_A_B, 5IL2_A_B, 5IM0_A_B, 5IMK_B_A, 5IML_A_B, 5IMM_B_A, 5INB_A_B, 5IP4_A_B, 5IT2_H_L, 5ITB_H_L, 5IU0_A_B, 5IWB_A_B, 5J03_A_B, 5J1T_A_B, 5J3T_A_B, 5J56_A_B, 5J57_A_B, 5JB8_E_S, 5JB9_E_S, 5JBA_E_S, 5JBB_E_S, 5JBC_E_S, 5JBT_A_Y, 5JCA_L_S, 5JDS_B_A, 5JEJ_A_B, 5JEL_A_B, 5JFC_L_S, 5JFD_L_H, 5JGE_D_E, 5JI0_A_D, 5JJE_A_B, 5JKL_E_F, 5JKM_E_F, 5JLB_A_B, 5JM1_A_G, 5JQA_A_B, 5JQY_A_B, 5JR1_H_L, 5JRP_L_H, 5JSY_A_B, 5JTH_A_B, 5JUE_L_H, 5JXA_H_L, 5JZY_L_H, 5K39_A_B, 5K7M_A_B, 5K7U_A_B, 5K7W_A_B, 5K8A_H_V, 5K9J_H_L, 5KDO_A_B, 5KNH_D_I, 5KP7_A_B, 5KP8_A_B, 5KSA_B_D, 5KVE_E_L, 5KVG_L_H, 5KVL_H_L, 5KY0_A_B, 5KY5_A_B, 5KY7_A_B, 5KY8_A_B, 5KY9_A_B, 5L0U_A_B, 5L20_A_B, 5L21_A_B, 5L23_A_B, 5L2Y_H_L, 5L2Z_H_L, 5L30_H_L, 5L6D_A_B, 5L6E_A_B, 5L6M_J_K, 5L6N_L_H, 5L6Y_H_L, 5L7E_A_B, 5L7H_A_B, 5L7X_H_L, 5L88_H_L, 5L8H_A_B, 5L8L_A_B, 5L8X_A_B, 5L9B_A_B, 5L9D_H_L, 5L9Y_A_B, 5L9Z_A_B, 5LB7_A_B, 5LCE_L_H, 5LCP_A_B, 5LCQ_A_B, 5LCR_A_B, 5LCT_A_B, 5LCU_A_B, 5LDA_A_B, 5LGH_H_L, 5LI1_A_B, 5LLB_C_D, 5LNP_B_C, 5LPD_L_H, 5LQB_H_L, 5LY3_A_B, 5LYN_A_B, 5M01_A_H, 5M0B_A_B, 5M0C_A_B, 5M0L_A_B, 5M0U_A_B, 5M0Y_B_A, 5M13_A_B, 5M2J_A_D, 5M2O_A_B, 5M45_G_J, 5M6V_A_B, 5M6Y_A_B, 5M71_A_B, 5M72_A_B, 5M75_A_B, 5MAW_D_E, 5MBL_A_B, 5MCQ_A_D, 5MDJ_L_S, 5MDK_L_S, 5MDL_L_S, 5ME5_A_B, 5MEO_A_B, 5MHC_A_P, 5MHI_A_B, 5MJ5_A_B, 5MJT_L_H, 5MKU_A_B, 5MLS_L_H, 5MM6_L_H, 5MO3_H_L, 5MOC_A_P, 5MP6_H_L, 5MRV_A_B, 5MTJ_A_B, 5MV8_A_B, 5MVV_A_G, 5MYC_A_P,

5MYK_A_B, 5MYO_C_D, 5N1D_A_B, 5N1E_A_B, 5N1K_A_B, 5N1L_A_B, 5N1M_A_B, 5N1N_A_B, 5N1O_A_B, 5N1Y_A_B, 5N32_A_B, 5N33_A_B, 5N36_A_B, 5N37_A_B, 5N39_A_B, 5N3A_A_B, 5N3B_A_B, 5N3C_A_B, 5N3D_A_B, 5N3E_A_B, 5N3F_A_B, 5N3U_A_B, 5N48_A_C, 5N4J_L_H, 5N5P_A_C, 5N7E_A_B, 5N7P_A_B, 5N7U_A_B, 5N85_A_B, 5NCT_A_C, 5NCU_A_B, 5NCW_A_B, 5NES_C_D, 5NEY_B_D, 5NF0_B_C, 5NGQ_B_C, 5NGV_H_L, 5NHN_B_A, 5NHW_H_L, 5NI9_A_B, 5NIB_A_C, 5NIG_A_B, 5NIV_A_B, 5NMK_A_B, 5NMV_L_H, 5NPH_H_L, 5NPI_A_B, 5NPJ_A_B, 5NPZ_A_B, 5NQ0_A_B, 5NQ2_A_B, 5NQF_A_B, 5NRM_A_B, 5NTJ_A_B, 5NX1_A_C, 5O02_A_C, 5O0E_A_C, 5O5M_A_C, 5O8W_A_B, 5O9E_A_B, 5OB0_A_B, 5OB1_A_B, 5OB2_A_C, 5OB5_H_L, 5OBF_H_L, 5OCK_L_H, 5ODB_A_B, 5OHG_B_I, 5OJR_A_B, 5OJW_A_B, 5OK3_A_D, 5OL0_A_B, 5OL3_A_D, 5OL4_B_C, 5OM2_A_B, 5OM3_A_B, 5OM5_A_B, 5OM7_A_B, 5OOV_A_B, 5OTG_A_D, 5OTT_A_B, 5OTU_A_C, 5OTX_A_C, 5OUA_A_B, 5OUC_A_E, 5OVO_A_B, 5OW0_B_A, 5OWU_A_B, 5OXZ_A_B, 5PA8_A_C, 5PA9_A_C, 5PAA_A_C, 5PAB_H_L, 5PAE_A_B, 5PAF_A_B, 5PAG_A_B, 5PAI_A_B, 5PAK_A_C, 5PAM_A_B, 5PAN_A_B, 5PAO_A_C, 5PAQ_A_B, 5PAT_A_B, 5PAU_A_C, 5PAV_A_C, 5PAX_A_C, 5PAY_A_C, 5PB0_A_B, 5PB1_A_D, 5PB2_A_C, 5PB3_A_C, 5PB5_A_B, 5PB6_A_C, 5Q0I_A_B, 5Q12_A_B, 5Q1E_A_B, 5Q1I_A_B, 5QU9_A_B, 5R0D_A_B, 5R0L_A_B, 5R0O_A_B, 5R0T_A_B, 5SVX_A_B, 5SVY_A_B, 5SW7_A_B, 5SWQ_A_B, 5SY8_H_L, 5SZB_A_H, 5SZC_A_H, 5T7G_A_C, 5T86_A_I, 5TDE_A_B, 5TDF_A_B, 5TDR_A_B, 5TDW_A_B, 5TDZ_A_B, 5TED_B_A, 5TEG_A_B, 5TEZ_A_J, 5TGI_A_B, 5TL5_L_H, 5TNO_A_B, 5TNT_A_B, 5TPP_L_H, 5TQE_H_L, 5TQF_H_L, 5TQG_H_L, 5TVO_A_B, 5TX4_A_B, 5TXK_A_B, 5TXS_A_B, 5TY6_H_L, 5TZP_A_B, 5U3D_A_B, 5U3N_H_L, 5U3O_H_L, 5U3P_H_L, 5U52_A_B, 5U5F_A_B, 5U5M_A_B, 5U66_B_A, 5U6A_A_B, 5U8C_A_B, 5UCB_H_L, 5UD9_H_L, 5UEK_H_L, 5UEL_H_L, 5UKN_H_L, 5UKP_H_L, 5UL6_A_M, 5ULH_A_B, 5UMM_A_C, 5URC_B_D, 5UUK_A_B, 5UUL_A_B, 5UUP_A_B, 5UW3_B_C, 5V02_B_R, 5V03_B_R, 5V2P_A_B, 5V3R_A_B, 5V62_A_I, 5V89_A_C, 5VAB_A_F, 5VAC_A_C, 5VAG_B_C, 5VAK_A_B, 5VGB_A_B, 5VHB_A_B, 5VKD_H_L, 5VKO_A_B, 5VLI_A_B, 5VMO_A_B, 5VPG_A_D, 5VPL_A_C, 5VT9_A_B, 5VUD_A_B, 5VUE_A_B, 5VUF_A_B, 5VVP_A_B, 5VWD_A_B, 5VWF_A_B, 5VWH_A_B, 5VWI_A_B, 5VWJ_A_B, 5VWV_A_B, 5VWY_A_B, 5W05_L_H, 5W0D_A_B, 5W3P_L_H, 5W5C_C_F, 5W6A_A_C, 5W6C_H_L, 5W89_A_B, 5WA1_A_B, 5WBX_B_R, 5WCD_H_L, 5WEQ_A_B, 5WK2_L_H, 5WKE_A_B, 5WMO_A_B, 5WMP_A_B, 5WMQ_A_B, 5WMR_A_B, 5WN9_H_A, 5WRD_A_B, 5WRV_A_B, 5WSH_A_B, 5WUV_L_H, 5WVO_B_C, 5WXH_A_C, 5WXK_A_B, 5WXL_A_C, 5WXO_U_P, 5WXQ_U_P, 5WXR_U_P, 5WY2_A_C, 5X03_A_B, 5X5X_H_L, 5XA5_A_B, 5XBF_A_B, 5XCO_A_B, 5XCQ_A_B, 5XCR_A_B, 5XCT_A_B, 5XEC_C_A, 5XHG_C_D, 5XIU_A_B, 5XLE_S_L, 5XLF_S_L, 5XLG_S_L, 5XLH_S_L, 5XLN_A_B, 5XLU_A_B, 5XMY_A_C, 5XOQ_A_B, 5XOS_A_B, 5XU8_A_B, 5XVE_A_B, 5XW1_A_B, 5XW5_A_B, 5XW9_A_B, 5XWA_A_B, 5XXK_A_B, 5XZH_A_C, 5Y1J_A_U, 5Y21_A_B, 5Y27_A_B, 5Y4N_S_L, 5Y5S_C_M, 5Y8X_A_B, 5Y91_A_B, 5Y96_A_B, 5Y9K_L_H, 5YAY_A_B, 5YCA_A_C, 5YE3_B_A, 5YIP_A_B, 5YMW_G_J, 5YN5_A_B, 5YN6_A_B, 5YN8_A_B, 5YNB_A_B, 5YNF_A_B, 5YNM_A_B, 5YNN_A_B, 5YNO_A_B, 5YNQ_A_B, 5YOF_A_B, 5YR0_A_B, 5YT0_A_B, 5YWR_A_B, 5Z0E_A_B, 5Z0G_A_B, 5Z0H_A_B, 5Z0I_A_B, 5Z0J_A_B, 5Z0K_A_B, 5Z0L_A_B, 5Z6S_A_C, 5ZJY_A_B, 5ZJZ_A_B, 5ZK9_A_B, 5ZMJ_H_L, 5ZML_A_B, 5ZOO_G_A, 5ZQG_A_B, 5ZQP_A_B, 5ZQR_A_B, 5ZRZ_A_B, 5ZWX_A_B, 5ZYS_A_B, 6A3W_C_I, 6A76_L_H, 6A7T_A_B, 6A7V_C_G, 6A84_A_B, 6A9K_L_A, 6AAW_A_B, 6ACI_A_H, 6AE8_A_B, 6AOD_B_C, 6AOR_A_B, 6AOT_A_B, 6AOU_A_B, 6AOV_A_B, 6APC_H_L, 6APP_A_B, 6AQ7_H_L, 6AR2_A_B, 6AT6_B_A, 6ATH_A_B, 6ATI_B_E, 6ATV_A_M, 6AU8_A_C, 6AZM_A_C, 6B05_A_B, 6B0G_C_D, 6B0S_H_L, 6B0W_L_H, 6B12_B_C, 6B4E_A_B, 6B5N_H_L, 6B5R_H_L, 6B5S_H_L, 6B7M_A_B, 6B7O_A_C, 6B9H_A_B, 6B9Z_A_B, 6BBL_B_D, 6BC8_A_B, 6BCB_F_A, 6BCD_A_B, 6BDV_A_B, 6BE2_H_L, 6BE3_H_L, 6BFJ_A_B, 6BFL_A_B, 6BFO_A_B, 6BFS_L_H, 6BG1_A_C, 6BGS_A_C, 6BHA_A_B, 6BHD_A_B, 6BI2_H_L, 6BJ8_A_H, 6BKR_A_B, 6BLA_H_L, 6BLH_L_H, 6BLQ_A_B, 6BLR_A_B, 6BQB_L_H, 6BSB_A_B, 6BSC_A_B, 6BTJ_H_L, 6BVH_A_I, 6BVI_A_B, 6BVJ_A_B, 6BVK_B_C, 6BVL_A_B, 6BVM_B_C, 6BW9_A_B, 6BXP_A_B, 6BXQ_B_C, 6BZ9_A_B, 6BZY_H_L, 6C5H_H_L, 6C5I_H_L, 6C6J_A_B, 6C73_A_B, 6CA7_L_H, 6CBV_H_L, 6CHA_B_F, 6CKZ_A_B, 6CL0_A_B, 6CO6_A_B, 6CO9_A_B, 6COJ_A_B, 6CR1_L_H, 6CT7_H_B, 6CUO_A_B, 6CUP_A_B, 6CUR_A_B, 6CW9_D_A, 6CXT_A_B, 6D0X_A_B, 6D29_A_B, 6D2R_A_B, 6D2T_A_B, 6D3Y_A_C, 6D3Z_A_C, 6D40_A_C, 6D55_B_C, 6D56_A_B, 6D59_A_B, 6D5E_B_C, 6D5G_B_C, 6D5H_B_C, 6D5J_A_B, 6D5L_A_B, 6D64_A_B, 6D7Y_A_B, 6DAQ_C_D, 6DB6_H_L, 6DBF_A_B, 6DC4_H_L, 6DC7_L_M, 6DC8_L_H, 6DCW_L_H, 6DDM_A_B, 6DEY_A_D, 6DGN_A_B, 6DI4_B_D, 6DN7_A_C, 6DNO_A_B, 6DRE_A_B, 6DUC_A_B, 6DZ4_A_B, 6DZO_A_B, 6E3I_A_B, 6E3J_A_B, 6E48_A_B, 6E5X_A_B, 6E65_L_H, 6EA3_A_B, 6EGW_A_B, 6EH2_A_B, 6EH3_A_C, 6EH4_D_E, 6EH5_A_B, 6EH6_A_B, 6EH7_A_B, 6EI1_A_B, 6EI2_A_B, 6EM6_A_C, 6EM7_A_D, 6EMA_A_C, 6EMB_A_D, 6EMD_A_D, 6ER6_B_A, 6ERJ_B_A, 6ERS_A_B, 6ERT_A_D, 6ERW_A_D, 6ES1_A_B, 6ESA_A_E, 6F0F_A_B, 6F0W_A_S, 6F14_A_B, 6F45_D_B, 6F4U_A_D, 6F4V_A_G, 6F6D_A_B, 6F6R_A_B, 6FBA_A_C, 6FBX_A_B, 6FBZ_A_B, 6FC0_A_B, 6FC3_A_B, 6FDK_A_B, 6FFA_A_B, 6FG8_A_B, 6FGE_A_C, 6FJT_L_H, 6FLC_A_L, 6FP7_A_B, 6FP8_A_B, 6FR4_A_B, 6FR5_A_B, 6FR9_A_B, 6FRC_A_B, 6FRX_A_B, 6FTO_B_C, 6FTP_A_B, 6FUB_B_A, 6FUD_A_B, 6FUM_A_B, 6FUN_A_B, 6FUP_A_B, 6G10_B_C, 6G48_B_C, 6G4J_A_B, 6G5G_A_B, 6G9Q_A_H, 6GBW_L_H, 6GCL_A_D, 6GEV_A_B, 6GFX_A_C, 6GGG_A_B, 6GJS_A_B, 6GKU_H_L, 6GP7_B_A, 6GPZ_B_A, 6GQN_A_B, 6GR8_A_B, 6GSC_A_B, 6GU2_A_B, 6GUC_A_C, 6GUE_A_C, 6GUM_A_B, 6GWB_B_A, 6GWJ_B_K, 6GY5_A_U, 6GZJ_A_B, 6GZS_A_B, 6H1D_A_B, 6H1E_A_B, 6H2U_A_B, 6H47_A_B, 6H6W_B_A, 6H74_B_A, 6H8J_B_C, 6H9J_A_D, 6H9U_A_B, 6HA6_A_D, 6HAR_A_E, 6HC8_A_E, 6HEI_A_B, 6HER_A_B, 6HFA_A_B, 6HGD_A_B, 6HGF_A_B, 6HGG_A_B, 6HGI_A_B, 6HGJ_A_B, 6HGK_A_B, 6HGL_A_B, 6HGM_A_B, 6HGN_A_B, 6HKG_A_B, 6HKP_A_S, 6HL1_A_B, 6HL2_B_D, 6HL5_A_S, 6HL6_A_S, 6HM3_A_B, 6HPR_B_C, 6HRN_A_B, 6HSO_A_I, 6HSX_L_H, 6HW2_A_B, 6HY2_X_A, 6I1S_A_B, 6I2A_A_D, 6I2C_A_B, 6I2D_A_D, 6I2G_A_B, 6I2H_A_D, 6I42_A_B, 6I4M_A_G, 6I51_L_H, 6I68_A_C, 6I7Q_B_V, 6I7R_B_V, 6IDX_A_C, 6IEU_A_C, 6IF3_A_B, 6IFC_E_G, 6IFG_A_B, 6IGK_A_B, 6IN7_A_B, 6IQJ_B_A, 6IR1_A_B, 6IR2_A_B, 6IU7_A_B, 6IUA_A_B, 6IWD_A_B, 6IXX_A_I, 6IYH_A_B, 6J14_A_B, 6J19_A_B, 6J1W_A_B, 6J29_A_B, 6J2A_A_B, 6J4P_A_B, 6J4U_A_B, 6J56_A_B, 6J5D_H_L, 6J5F_H_L, 6J7B_A_B, 6J8O_A_B, 6J9J_A_B, 6J9O_H_L, 6JAU_A_B, 6JB2_A_B, 6JB5_A_B, 6JB8_A_B, 6JH9_A_B, 6JHZ_A_B, 6JLE_A_E, 6JMU_A_B, 6JP7_H_L, 6JTN_A_B, 6JTO_A_B, 6JWJ_A_C, 6JWN_A_C, 6JX3_B_A, 6JY3_A_C, 6JY4_A_C, 6K0Y_A_B, 6K3B_A_B, 6K3M_H_A, 6K64_A_B, 6K65_L_H, 6K6A_B_A, 6KAO_A_B, 6KAP_A_B, 6KAQ_A_B, 6KAR_A_B, 6KAS_B_D, 6KAT_B_D, 6KAU_B_D, 6KAV_B_D, 6KBR_A_C, 6KGC_A_B, 6KGD_A_B, 6KHS_A_B, 6KIP_A_B, 6KKB_X_D, 6KM7_A_B, 6KMJ_A_C, 6KMR_A_B, 6KN1_A_C, 6KRO_A_B, 6KWL_A_B, 6KX1_A_B, 6KXD_A_B, 6KXE_A_B, 6KXF_A_B, 6KXX_A_B, 6KYI_A_B, 6KZ1_A_B, 6KZJ_A_C, 6L4P_A_B, 6L5V_A_B, 6L5W_A_B, 6L5X_B_D, 6L5Y_B_D, 6L7R_A_B, 6LB4_A_B, 6LDV_H_L, 6LIT_A_B, 6LKT_A_B, 6LPH_A_B, 6LRA_H_L, 6LSB_A_B, 6LTG_B_D, 6LUN_B_D, 6M01_A_B, 6M0Q_I_K, 6M3I_A_B, 6MA3_A_B, 6MA4_A_B, 6MA5_A_B, 6MBB_A_B, 6MBC_A_B, 6MEE_C_D, 6MEG_H_L, 6MEH_H_L, 6MFG_E_F, 6MGN_A_B, 6MIB_A_B, 6MJ4_D_A, 6MJJ_D_A, 6MM5_E_C, 6MM8_C_D, 6MNQ_H_L, 6MRQ_A_I, 6MS7_A_B, 6MT3_A_C, 6MT6_A_C, 6MTL_A_B, 6MV4_H_L, 6MVL_H_L, 6MYD_A_C, 6MYE_A_B, 6MYU_A_B, 6N2N_A_C, 6N87_A_C, 6NB8_H_L, 6NBE_A_N, 6NE2_B_A, 6NE4_B_A, 6NMT_A_B, 6NRQ_A_C, 6NUC_A_B, 6NUY_L_H, 6NUZ_L_H, 6NWK_A_B, 6NWL_A_B, 6NYO_A_E, 6NYQ_L_C, 6NYX_D_F, 6O17_A_B, 6O21_A_B, 6O24_A_B, 6O26_A_B, 6O3K_L_H, 6O3X_A_C, 6O40_A_B, 6O4Y_A_B, 6O4Z_A_B, 6O51_A_B, 6O53_A_B, 6O7O_B_D, 6O7Q_B_D, 6OBC_A_B, 6OBE_A_B, 6OC7_H_L, 6OCG_A_B, 6ODD_A_B, 6OJ7_A_C, 6OM4_A_B, 6ON9_A_B, 6ONB_A_B, 6ONO_C_D, 6OP2_B_D, 6OPD_A_B, 6OR0_A_B, 6OSH_L_H, 6OSV_L_H, 6OTC_H_L, 6OVE_A_B, 6OVK_R_B, 6OVN_A_B, 6OXC_A_B, 6OXD_A_B, 6P23_A_B, 6P27_A_B, 6P2C_A_B, 6P43_A_C, 6P4Y_H_L, 6P4Z_B_D, 6P79_H_L, 6P8O_A_B, 6P8S_A_B, 6P8U_A_B, 6PBH_A_B, 6PDI_A_B, 6PDR_L_H,

6PDU_L_H, 6PE7_H_L, 6PEC_L_H, 6PEU_A_B, 6PHG_A_B, 6PJX_A_B, 6PLH_A_B, 6PV9_A_B, 6PXC_A_U, 6PYV_A_B, 6PZG_L_H, 6PZP_A_B, 6Q00_A_B, 6Q19_L_H, 6Q1G_L_H, 6Q3K_A_B, 6Q3P_A_B, 6Q41_B_A, 6Q43_B_C, 6Q6X_B_C, 6Q85_C_D, 6Q8G_C_D, 6Q8Q_A_B, 6Q9F_A_B, 6QB3_A_X, 6QBA_A_B, 6QBG_A_B, 6QBH_A_B, 6QCB_A_B, 6QCU_L_H, 6QDY_B_C, 6QFC_A_B, 6QFK_A_B, 6QGR_A_B, 6QGT_A_B, 6QGW_A_B, 6QIO_A_B, 6QJ7_A_B, 6QQ7_A_B, 6QTA_A_B, 6QTB_A_D, 6QTR_A_B, 6QUP_A_B, 6QZY_A_B, 6R00_A_B, 6R15_A_B, 6R2G_A_C, 6R2I_A_B, 6R3V_A_B, 6R6M_A_B, 6R6O_A_B, 6R7W_A_B, 6R8I_A_B, 6R8K_B_C, 6RFK_E_S, 6RH7_A_B, 6RHC_A_P, 6RJZ_A_P, 6RKG_B_C, 6RKI_A_P, 6RKM_A_P, 6RL3_A_P, 6RL6_A_P, 6RM9_A_C, 6RMV_A_B, 6RN7_A_B, 6RN9_A_B, 6RNE_A_B, 6RNI_A_B, 6RNK_A_B, 6ROT_L_H, 6RP1_B_C, 6RP5_A_B, 6RSW_A_C, 6RTW_A_B, 6RUE_A_D, 6RWH_A_P, 6RWI_A_P, 6RWS_A_P, 6RWU_A_P, 6S07_A_C, 6S0Y_A_B, 6S1U_A_B, 6S22_A_F, 6S29_A_C, 6S3F_A_B, 6S40_A_P, 6S4I_A_B, 6S4J_A_B, 6S5A_A_H, 6S5P_B_D, 6S7G_A_D, 6S9Q_A_P, 6SBA_A_B, 6SIP_A_P, 6SQK_A_B, 6SWT_A_B, 6SYI_A_B, 6T2D_A_B, 6T3Y_A_B, 6T40_A_C, 6T4A_L_H, 6T52_L_H, 6T54_L_H, 6T55_L_H, 6T56_L_H, 6T57_L_H, 6T89_L_H, 6T8A_L_H, 6TDO_A_B, 6TDP_A_B, 6TDT_H_L, 6TGX_A_B, 6TKG_L_H, 6TKH_L_H, 6TKI_L_H, 6TKL_L_H, 6TQP_A_B, 6TRN_A_B, 6TTP_A_B, 6TTX_A_B, 6TYU_A_B, 6TYV_A_B, 6TYW_A_B, 6TYZ_A_B, 6U0J_A_B, 6U12_A_B, 6U1T_H_L, 6U22_A_C, 6U3B_A_B, 6U54_A_B, 6U55_A_B, 6U8M_A_B, 6UA3_A_B, 6UC5_H_L, 6UCE_L_H, 6UCF_L_H, 6UG8_L_H, 6UOE_H_L, 6UUD_H_L, 6UXE_A_D, 6UYP_A_B, 6UYQ_A_B, 6UYR_A_B, 6UYT_A_B, 6UYV_A_B, 6UYX_C_A, 6UYY_A_B, 6UZM_A_B, 6V2E_A_B, 6V2O_A_B, 6V3J_A_G, 6V4E_A_B, 6V4F_A_B, 6V4G_A_B, 6V4Q_H_L, 6V94_A_B, 6V9F_B_C, 6V9J_A_B, 6V9L_A_B, 6V9M_A_B, 6V9N_A_B, 6V9O_A_B, 6VB0_A_B, 6VB1_A_B, 6VB2_A_B, 6VBO_L_H, 6VER_A_B, 6VES_A_B, 6VFD_A_B, 6VI2_B_D, 6VJT_H_L, 6VK6_A_B, 6VNT_A_B, 6VQD_A_B, 6VQE_A_B, 6VRY_L_H, 6VTC_A_B, 6VTM_D_G, 6VW9_A_B, 6W00_L_H, 6W1D_A_D, 6W4H_A_B, 6W4W_L_H, 6W61_A_B, 6W7Z_A_B, 6W9L_A_B, 6W9M_A_B, 6WAS_L_A, 6WDU_A_B, 6WG0_H_L, 6WGN_A_C, 6WI2_A_D, 6WIH_A_D, 6WIY_H_L, 6WJ0_H_L, 6WJT_A_C, 6WKL_A_B, 6WMD_A_B, 6WOS_H_L, 6WUD_A_B, 6WVN_A_B, 6WY9_A_B, 6WZY_A_B, 6X00_A_B, 6X0C_A_B, 6X5G_A_B, 6X8H_A_B, 6XAW_A_B, 6XC1_A_C, 6XCA_H_L, 6XDL_A_B, 6XDT_B_D, 6XE3_A_B, 6XIB_A_B, 6XID_A_B, 6XIF_A_B, 6XJR_A_C, 6XOY_A_B, 6XPB_A_B, 6XRH_A_B, 6XRU_A_B, 6XT0_A_B, 6XTG_L_H, 6XTL_A_C, 6XTM_A_C, 6XTN_A_C, 6XTO_A_C, 6XTQ_A_C, 6XTR_A_C, 6XUD_H_L, 6XUK_H_L, 6XZF_A_B, 6XZU_A_B, 6Y02_L_H, 6Y05_A_B, 6Y0B_A_B, 6Y0U_C_D, 6Y0V_C_D, 6Y1K_H_L, 6Y1L_H_L, 6Y1M_H_L, 6Y26_A_B, 6Y27_A_B, 6Y28_A_B, 6Y29_A_B, 6Y2U_A_B, 6Y3B_A_B, 6Y49_H_L, 6Y4O_A_B, 6Y89_A_B, 6Y8C_A_B, 6Y9H_L_H, 6YA6_A_B, 6YA7_A_B, 6YA8_A_B, 6YB6_L_H, 6YCR_A_B, 6YD0_B_D, 6YDI_B_D, 6YDU_B_D, 6YHG_L_H, 6YHJ_L_H, 6YHQ_H_L, 6YIF_A_B, 6YMP_L_H, 6YN3_L_H, 6YNA_A_B, 6YNC_A_B, 6YNR_A_B, 6YNT_A_B, 6YOT_A_B, 6YPP_A_B, 6YQG_A_B, 6YQI_A_B, 6YQJ_A_B, 6YQK_A_B, 6YQV_L_H, 6YSJ_L_H, 6YSX_L_H, 6YU8_A_B, 6YW1_A_B, 6YW4_A_B, 6YYX_A_B, 6YZ5_E_F, 6Z1V_A_B, 6Z2F_A_B, 6ZDT_A_B, 6ZMP_A_B, 6ZQS_A_B, 6ZRV_A_B, 6ZUG_L_H, 6ZUH_L_H, 6ZUU_L_H, 6ZUW_L_H, 6ZUX_L_H, 6ZV7_L_H, 7A48_A_B, 7A77_A_B, 7A78_A_B, 7B1I_B_C, 7BMI_A_B, 7BNT_B_C, 7BP2_B_D, 7BP5_A_B, 7BP6_C_D, 7BPH_A_B, 7BQ1_A_B, 7BQ2_A_B, 7BQ3_A_B, 7BQ4_A_B, 7BQU_A_B, 7BQV_A_B, 7BU3_A_B, 7BU5_A_B, 7BZ5_H_L, 7BZK_A_B, 7C2B_A_C, 7C2G_A_G, 7CD8_B_A, 7CDL_G_M, 7CE4_A_B, 7CIZ_E_F, 7CMZ_A_B, 7CN7_A_C, 7CNC_A_B, 7CNS_A_B, 7CQ3_A_B, 7D2Q_C_G, 7D2S_A_B, 7D2Z_A_B, 7D5B_A_D, 7D6Q_A_C, 7D6R_A_C, 7D9E_A_B, 7DA0_B_C, 7DG2_A_C, 7DS8_A_B, 7E36_A_B, 7EAM_L_D, 7EAN_H_L, 7EK6_A_B, 7FAB_L_H, 7GCH_F_G, 7JHX_A_B, 7JLK_A_B, 7JMP_H_L, 7JYV_A_B, 7K2K_A_B, 7K3K_A_B, 7K3L_A_B, 7K81_A_G, 7K8N_A_B, 7K8R_H_L, 7K95_A_B, 7KD7_D_A, 7KGP_A_B, 7KGS_A_B, 7KMH_B_C, 7KMI_A_B, 7KPU_D_A, 7KZA_H_L, 7LFZ_A_B, 7LM9_H_L, 7LSG_H_L, 7M5F_A_C, 7MC5_A_M, 7MFU_A_D, 7MU2_A_C, 7NEH_H_L.

**Dataset S2. Dataset culled and used in Independent validation – 1 consisting of 106 binary PPI complexes (targets) from the Affinity benchmark v.2.** The comma-separated names consists of the PDB ID and the chain IDs for both receptor and ligand in the format: <PDB ID>_<chain ID of receptor>_<chain ID of ligand>. All these structures had annotated experimental binding affinities and/or free energies ($K_d$ or $\Delta G_{binding}$) associated with each entry (see section **2.1**, **Materials and Methods**).

1ACB_E_I, 1AK4_A_D, 1ATN_A_D, 1AVX_A_B, 1AVZ_B_C, 1AY7_A_B, 1B6C_A_B, 1BRS_A_D, 1BUH_A_B, 1BVN_P_T, 1DFJ_E_I, 1E6E_A_B, 1E96_A_B, 1EAW_A_B, 1EFN_B_A, 1EMV_A_B, 1EWY_A_C, 1F34_A_B, 1F6M_A_C, 1FC2_C_D, 1FFW_A_B, 1FLE_E_I, 1FQJ_A_B, 1GCQ_B_C, 1GL1_A_I, 1GLA_G_F, 1GPW_A_B, 1GRN_A_B, 1GXD_A_C, 1H1V_A_G, 1H9D_A_B, 1HE8_B_A, 1I2M_A_B, 1IBR_A_B, 1J2J_A_B, 1JIW_P_I, 1JTD_B_A, 1JTG_B_A, 1KAC_A_B, 1KTZ_A_B, 1KXP_A_D, 1LFD_B_A, 1M10_A_B, 1MAH_A_F, 1MQ8_A_B, 1NW9_B_A, 1OC0_A_B, 1OPH_A_B, 1PPE_E_I, 1PVH_A_B, 1PXV_A_C, 1QA9_A_B, 1R0R_E_I, 1R6Q_A_C, 1S1Q_A_B, 1T6B_X_Y, 1US7_A_B, 1UUG_A_B, 1WQ1_R_G, 1XQS_A_C, 1YVB_A_I, 1Z0K_A_B, 1ZHI_A_B, 1ZLI_A_B, 1ZM4_A_B, 2A9K_A_B, 2ABZ_B_E, 2AQ3_A_B, 2B42_A_B, 2BTF_A_P, 2C0L_A_B, 2FJU_B_A, 2GAF_D_A, 2GOX_A_B, 2HLE_A_B, 2HQS_A_H, 2HRK_A_B, 2I25_N_L, 2I9B_E_A, 2J0T_A_D, 2O3B_A_B, 2OOB_A_B, 2OUL_A_B, 2OZA_B_A, 2PCB_A_B, 2PTC_E_I, 2SIC_E_I, 2SNI_E_I, 2TGP_Z_I, 2UUY_A_B, 2VDB_A_B, 2WPT_A_B, 2X9A_D_C, 3BIW_A_E, 3BX7_A_C, 3BZD_A_B, 3CPH_G_A, 3DAW_A_B, 3F1P_A_B, 3H2V_A_E, 3K75_D_B, 3PC8_A_C, 3S9D_B_A, 3SGB_E_I, 4IZ7_A_B, 4M76_A_B.

**Dataset S2. Dataset culled and used in Independent validation – 2 consisting of 236 binary PPI complexes (targets) from Proximate and SKEMPI v.2.** The comma-separated names consists of the PDB ID and the chain IDs for both receptor and ligand in the format: <PDB ID>_<chain ID of receptor>_<chain ID of ligand>. All these structures had annotated experimental binding affinities and/or free energies ($K_d$ or $\Delta G_{binding}$) associated with each entry (see section **2.1, Materials and Methods**).

1A22_A_B, 1A4Y_A_B, 1ACB_E_I, 1AK4_A_D, 1B2S_A_D, 1B2U_A_D, 1B3S_A_D, 1B41_A_B, 1BP3_A_B, 1BRS_A_D, 1C1Y_A_B, 1C4Z_A_D, 1CF4_A_B, 1CSE_E_I, 1CSO_E_I, 1CT0_E_I, 1CT2_E_I, 1CT4_E_I, 1E50_A_B, 1E96_A_B, 1EAW_A_B, 1EFN_A_B, 1EMV_A_B, 1F47_A_B, 1F5R_A_I, 1FC2_C_D, 1FCC_A_C, 1FFG_A_B, 1FFW_A_B, 1FR2_A_B, 1FSS_A_B, 1FY8_E_I, 1GC1_G_C, 1GL0_E_I, 1GL1_A_I, 1GRN_A_B, 1GUA_A_B, 1H9D_A_B, 1HE8_A_B, 1IAR_A_B, 1JCK_A_B, 1JTD_A_B, 1JTG_A_B, 1K8R_A_B, 1KAC_A_B, 1KBH_A_B, 1KNE_A_P, 1KTZ_A_B, 1KU6_A_B, 1LFD_A_B, 1M9E_A_D, 1MAH_A_F, 1MQ8_A_B, 1OHZ_A_B, 1P69_A_B, 1P6A_A_B, 1PPF_E_I, 1R0R_E_I, 1S0W_A_C, 1S1Q_A_B, 1SBB_A_B, 1SBN_E_I, 1SGD_E_I, 1SGE_E_I, 1SGN_E_I, 1SGP_E_I, 1SGQ_E_I, 1SGY_E_I, 1SIB_E_I, 1SMF_E_I, 1T7C_A_B, 1TM1_E_I, 1TM3_E_I, 1TM4_E_I, 1TM5_E_I, 1TM7_E_I, 1TMG_E_I, 1TO1_E_I, 1UEA_A_B, 1UUZ_A_D, 1WQJ_I_B, 1X1W_A_D, 1X1X_A_D, 1XD3_A_B, 1XXM_A_C, 1Y1K_E_I, 1Y33_E_I, 1Y34_E_I, 1Y3B_E_I, 1Y3C_E_I, 1Y3D_E_I, 1Y48_E_I, 1Y4A_E_I, 1YCS_A_B, 1Z7X_W_X, 2A9K_A_B, 2ABZ_B_E, 2AJF_A_E, 2AW2_A_B, 2B0Z_A_B, 2B10_A_B, 2B11_A_B, 2B12_A_B, 2B42_A_B, 2B5I_A_B, 2BTF_A_P, 2C0L_A_B, 2C5D_A_C, 2CCL_A_B, 2DSQ_I_G, 2DVW_A_B, 2FTL_E_I, 2G2U_A_B, 2G2W_A_B, 2GOX_A_B, 2GYK_A_B, 2HLE_A_B, 2HRK_A_B, 2I26_N_L, 2I9B_A_E, 2J0T_A_D, 2J12_A_B, 2J1K_C_T, 2KSO_A_B, 2KWI_A_B, 2NOJ_A_B, 2NU0_E_I, 2NU1_E_I, 2NU2_E_I, 2NU4_E_I, 2O3B_A_B, 2OOB_A_B, 2PCB_A_B, 2PCC_A_B, 2REX_A_B, 2SGP_E_I, 2SGQ_E_I, 2SIC_E_I, 2VLN_A_B, 2VLO_A_B, 2VLP_A_B, 2VLQ_A_B, 2VN5_A_B, 2WPT_A_B, 3BIW_A_E, 3BK3_A_C, 3BP8_A_C, 3BT1_A_U, 3BTD_E_I, 3BTE_E_I, 3BTF_E_I, 3BTG_E_I, 3BTH_E_I, 3BTM_E_I, 3BTQ_E_I, 3BTT_E_I, 3BTW_E_I, 3BX1_A_C, 3C4P_A_B, 3D5R_A_C, 3D5S_A_C, 3EG5_A_B, 3EQS_A_B, 3EQY_A_C, 3F1S_A_B, 3KBH_A_E, 3KUD_A_B, 3LB6_A_C, 3LNZ_A_B, 3M62_A_B, 3M63_A_B, 3MZG_A_B, 3MZW_A_B, 3N06_A_B, 3N0P_A_B, 3N4I_A_B, 3NCB_A_B, 3NCC_A_B, 3NVN_B_A, 3NVQ_B_A, 3Q3J_A_B, 3Q8D_A_E, 3QHY_A_B, 3RF3_A_C, 3S9D_A_B, 3SE3_B_A, 3SE3_B_C, 3SE4_B_A, 3SE4_B_C, 3SEK_B_C, 3SF4_A_D, 3SGB_E_I, 3TGK_E_I, 3U82_A_B, 3UIG_A_P, 3UIH_A_P, 3UII_A_P, 3WWN_A_B, 4BFI_A_B, 4CPA_A_I, 4CVW_A_C, 4E6K_A_G, 4EKD_A_B, 4FZA_A_B, 4G0N_A_B, 4G2V_A_B, 4GNK_A_B, 4GU0_A_E, 4HRN_A_D, 4JEU_A_B, 4KRL_A_B, 4KRO_A_B, 4KRP_A_B, 4L0P_A_B, 4MYW_A_B, 4NFG_A_B, 4NZW_A_B, 4O27_A_B, 4OFY_A_D, 4RA0_A_C, 4RS1_A_B, 4UYP_A_D, 4UYQ_A_B, 4WND_A_B, 4XSS_B_E, 4Y0Y_E_I, 4Y61_A_B, 4YEB_A_B, 4YFD_A_B, 4YH7_A_B, 5CXB_A_B, 5CYK_A_B, 5E6P_A_B, 5F4E_A_B, 5K39_A_B, 5M2O_A_B, 5TAR_A_B, 5UFE_A_B, 5UFQ_A_C, 5XCO_A_B

**Note S1. Unresolved issues / unresponsed communications in order to procure enough experimental protein binding affinity data across different published datasets**

1. **dbMPIKT**: a database of kinetic and thermodynamic mutant protein interactions (http://deeplearner.ahu.edu.cn/web/dbMPIKT/) → cite not opening; No reply to address the issue after emailing.

2. **ProThermDB**: thermodynamic database for proteins and mutants revisited after 15 years; thermodynamic Database for Proteins and Mutants (ProThermDB) contains more than 32,000 data of several thermodynamic parameters such as melting temperature, free energy obtained with thermal and denaturant denaturation, enthalpy change, and heat capacity change along with experimental methods and conditions, sequence, structure, and literature information. (https://web.iitm.ac.in/bioinfo2/prothermdb/Organism.html) → No structural data, No Kd in the organism based data provided in .xlsx files (*https://web.iitm.ac.in/bioinfo2/prothermdb/Organism.html*) – No response from the support team in spite following the prescribed protocol for Downloading the entire database (*https://web.iitm.ac.in/bioinfo2/prothermdb/Downloads.html*) and repeated emails.

3. **PINT**: Protein–protein Interactions Thermodynamic Database (http://www.bioinfodatabase.com/pint/index.html) → redirecting to some protein manufacturing companies and unrelated commercial services: *http://ww1.bioinfodatabase.com/?subid1=624dfd50-ea23-11ed-8205-a1c46f4b1a2f*); No one cared to address the issue in spite of repeated emailing.

4. **ASEdb**: a database of alanine mutations (http://www.asedb.org/) → redirecting to GoDaddy services - not really what we are looking for independent validations

**Note S2. Detailed method for the SVM training and cross-validation in EnCPdock**

As mentioned in sections **2.2.2** (**Materials and Methods**) and **3.1 (Results and Discussion)** of the main manuscript, a ten-fold cross validation has been performed on the training dataset consisting of 3200 PPI complexes. In order to do that, the whole training dataset was first divided into ten equal parts (subsets) yielding a statistically significant number of datapoints (320 PPI complexes) in each subset. During training, each time, the regressor was trained on 9/10$^{th}$ of the training dataset and was tested on the leftover (1/10$^{th}$) fraction and this process was repeated for all combinations of the different subsets. One round of complete training in this way results in the attainment of 'predicted output' (PO) for each (target) entry in the training dataset. The predicted outputs (predicted $\Delta G_{binding\_norm}$) were then compared with their corresponding target functions (FoldX-derived $\Delta G_{binding\_norm}$) by means of evaluating the Pearson's correlation coefficient (r) on the entire training set (i.e., 3200 PPI complexes).

In addition, a threshold (i.e., cutoff) spanning the entire range of the target function was sampled at an interval of 0.025 from its minimum to maximum obtained values for each round of training to convert both the continuous functions (TF, PO) to binary (1/0) variables (see section **2.2.2.2, Materials and Methods**) each time leading to an optimized BACC score for that round of training (see section **2.2.3, Materials and Methods**). In order to find a suitable model that effectively describes the $\Delta G_{binding\_norm}$ as a robust function of the structural descriptors, a Gaussian Kernel function (Radial Basis Function or RBF Kernel) was used.

The SVM-training (10-fold cross-validation) was performed for a judiciously chosen range of values for the three main parameters part of svm_learn [1] guided by strong literature support: (i) the γ-parameter controlling the width of the RBF Kernel, (ii) the C-parameter presiding over the training error and margin width and (iii) the cost-function (ε). Initially, the SVM training was performed for a broad range of C and γ-values in a grid-search keeping the cost-function fixed (set to 0.1); the performance was tested for C-values between $2^{-15}$ to $2^{10}$, and the γ within the range of $2^{-10}$ to $2^{10}$ in $\log_2$ steps [2, 3]. The eventual aim was obviously to maximize the cross-validated r. The 'course-grained' search gave us an initial feel of the numbers based on which a more stringent search was invoked around those C and γ values which lead to the maximum r values in the initial search. Thus, in the subsequent fine-grained search, the values of the C, γ and ε were kept in the ranges of 1.0 to 5.0 (in steps of 0.5), 0.01 to 2.0 (in steps of 0.01) and 0.1 to 1.0 (in steps of 0.05) respectively (see section **2.2.2.1, Materials and Methods**).

**Figure S1. Variations of Pearson's correlation coefficient (r) with respect to the SVM parameters C, γ and ε:** Since we obtained maximum correlation at C= 4.5 and C=5.0 and at value of ε=1.00, the bar plot exhibiting r with respect to different γ values while C= 4.5, ε=1.00 (a), and C= 5.0, ε=1.00 (b), has been shown. (c) The maximum r obtained for different C values has been shown while ε=1.00 is kept fixed. (d) Keeping C= 4.5 and γ =0.05, the variations of ε has also been shown.

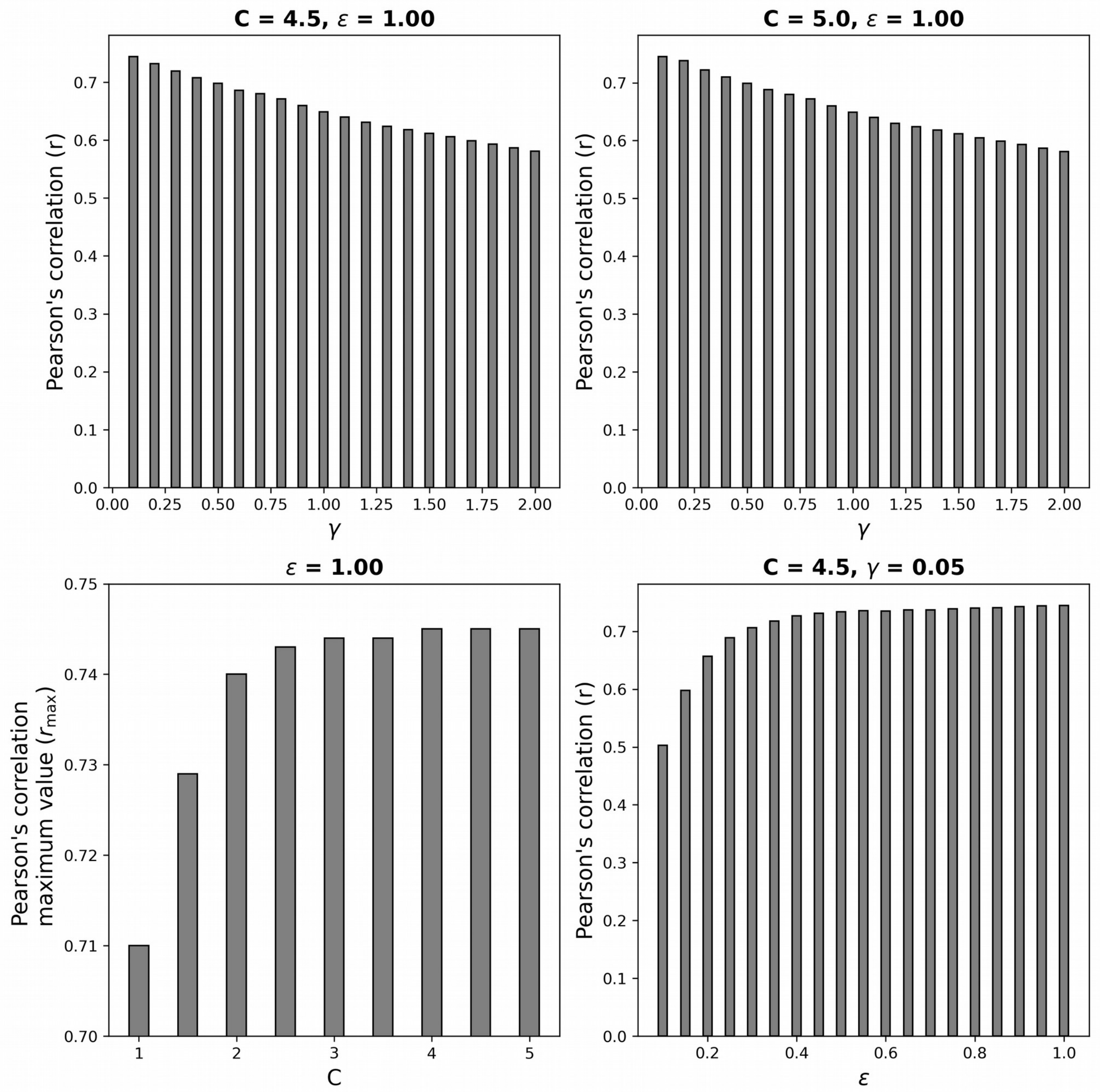

**Table S1. The top-performing SVM models together defining the EnCPdock predictor of protein binding energetics.** Each SVM model is tabulated in terms of its training parameters: C, γ and ε (see section **2.2.2**, **Materials and Mothods**) which were optimized in an 3-loop iterative cycle during the 10-fold cross-validation by maximizing the BACC score. Each of these models led to a BACC score of 0.833 with Pearson's correlation coefficients (r) narrowly varying within 0.744 to 0.745 obtained between the target function and predicted output. As can be seen in the Table, nine different (degenerate) combinations of C, γ and ε gave rise to the same hieghest BACC score which eventually led to the accumulation of 100 top SVM models (10 sets for each predictor, for 10-fold cross-validations) in the final predictor.

| **C** | **γ** | **ε** | **r** | **BACC** |
|---|---|---|---|---|
| 3.5 | 0.06 | 1.00 | 0.744 | 0.833 |
| 4.0 | 0.05 | 1.00 | 0.744 | 0.833 |
| 4.0 | 0.06 | 1.00 | 0.745 | 0.833 |
| 4.5 | 0.05 | 1.00 | 0.745 | 0.833 |
| 4.5 | 0.06 | 1.00 | 0.745 | 0.833 |
| 4.5 | 0.10 | 1.00 | 0.744 | 0.833 |
| 4.5 | 0.11 | 1.00 | 0.744 | 0.833 |
| 5.0 | 0.06 | 1.00 | 0.745 | 0.833 |
| 5.0 | 0.10 | 1.00 | 0.745 | 0.833 |

**Figure S2. The Receiver Operating Characteristics (ROC) curve pertaining to training and cross-validation of EnCPdock.** One of the ROC curves (TPR vs. FPR, see section **2.2.3**, **Materials and Methods**) characteristic of the top-performing models in cross-validation (i.e., BACC: 0.833) giving the hieghest correlation (r) of 0.745 between the target function and predicted output. The corresponding ROC-AUC (see section **2.2.3**) was found to be 0.75.

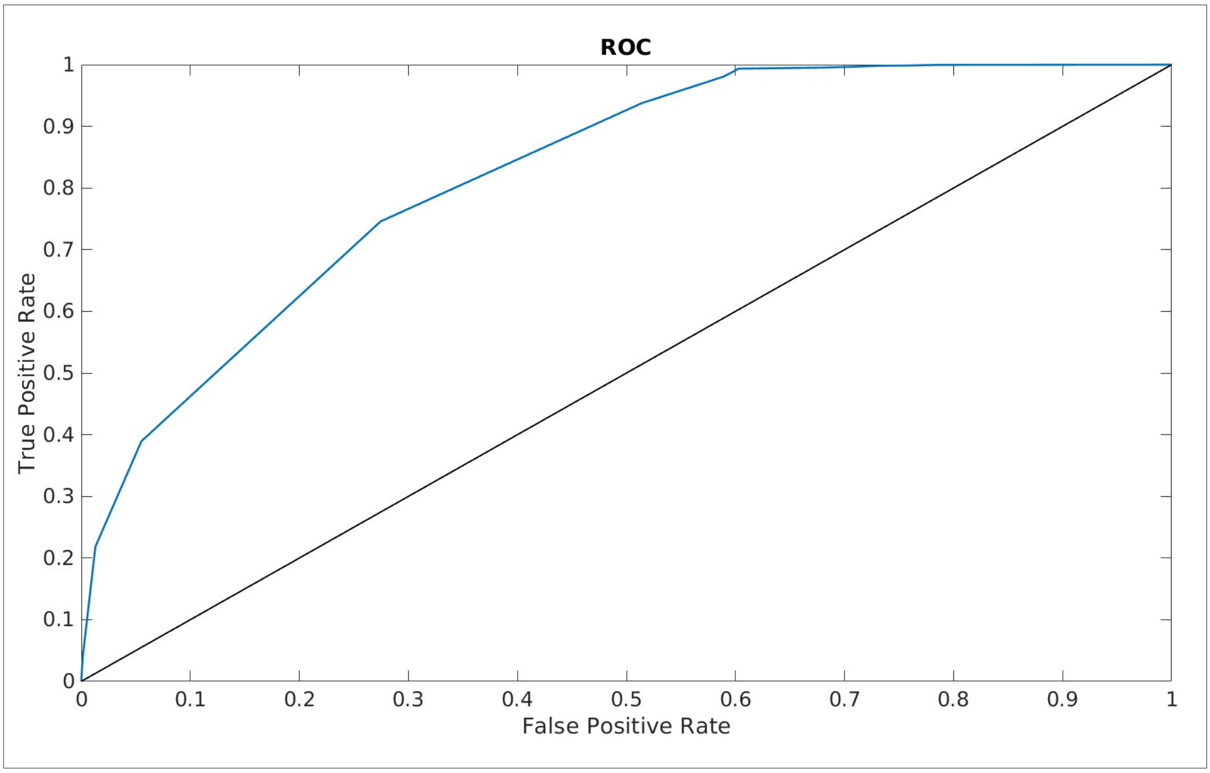

**Figure S3. Distribution of EnCPdock predictions across the top-performing SVM models.** The distribution of the ninety predicted $\Delta G_{EnCPdock\_norm}$ values (predicted by 90 top SVM models) for four PPI complexes (PDB IDs: 1ACB, 1BVN, 1KTZ and 1GPW shown in panels A, B, C and D respectively) randomly taken from the independent validation dataset. Together these and similar distributions justifies the choice of median as a better measure (more robbust) of central tendency than mean or mode.

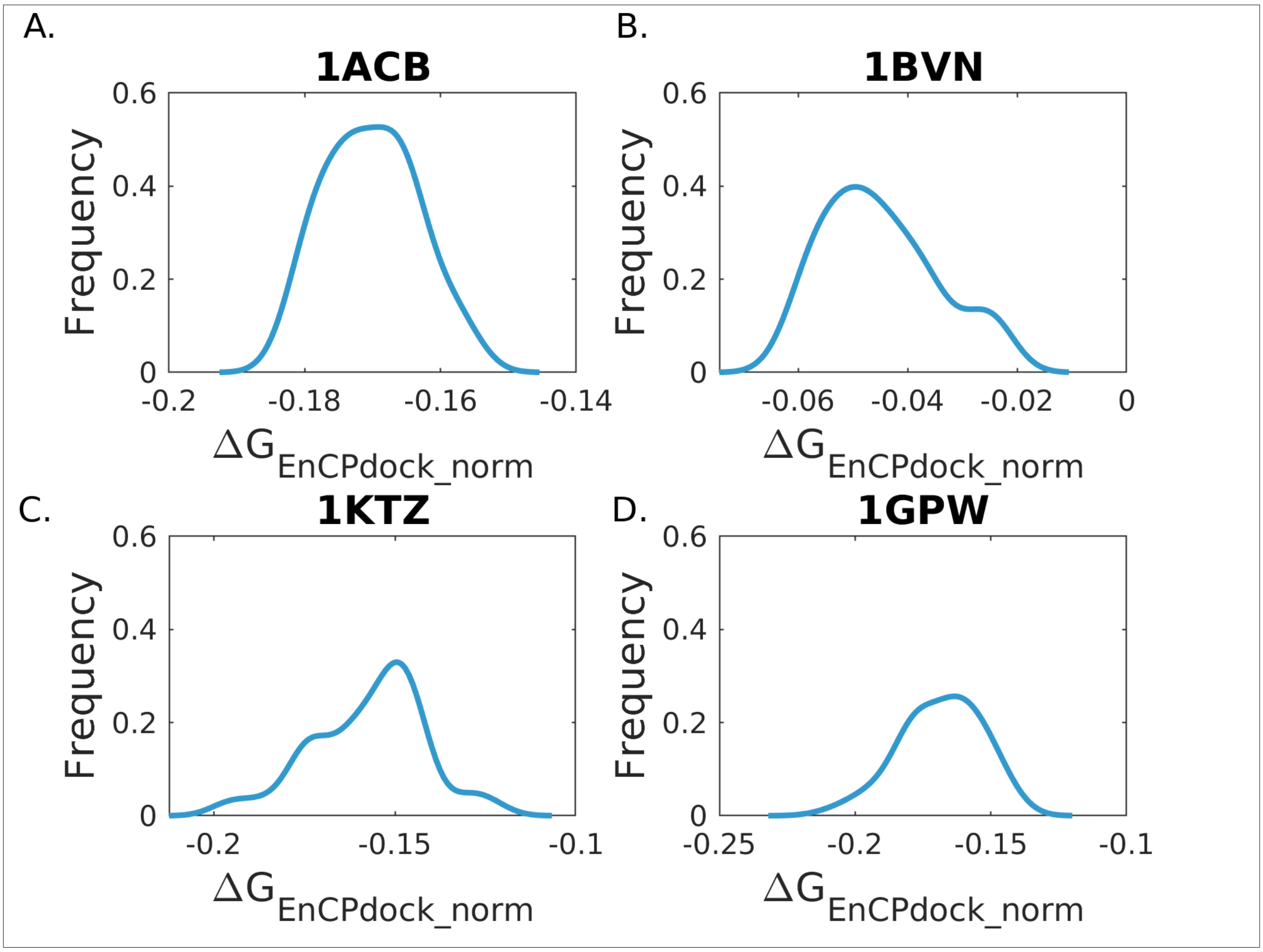

**Figure S4. Correlation (scatter) plots for independent validation - 1 (Affiint Benchmark v.2) for diffrent statistical descriptors (barrying meidan – the fianlly selected measure, dsipalyed as part of Main Text).** Plot of mean, mode, maximum and minimum values of $\Delta G_{EnCPdock\_norm}$ for 106 PPI complexes plotted against their corresponding $\Delta G_{FoldX\_norm}$ (panels A, B, C and D respectively) and $\Delta G_{exp\_norm}$ (panels E, F, G and H respectively) with their least-squares fitted lines. **Figure 4, 5** in the main-text portrays the same for medain, the chosen staitstical descriptor (for independent validations – 1, 2 repsectively).

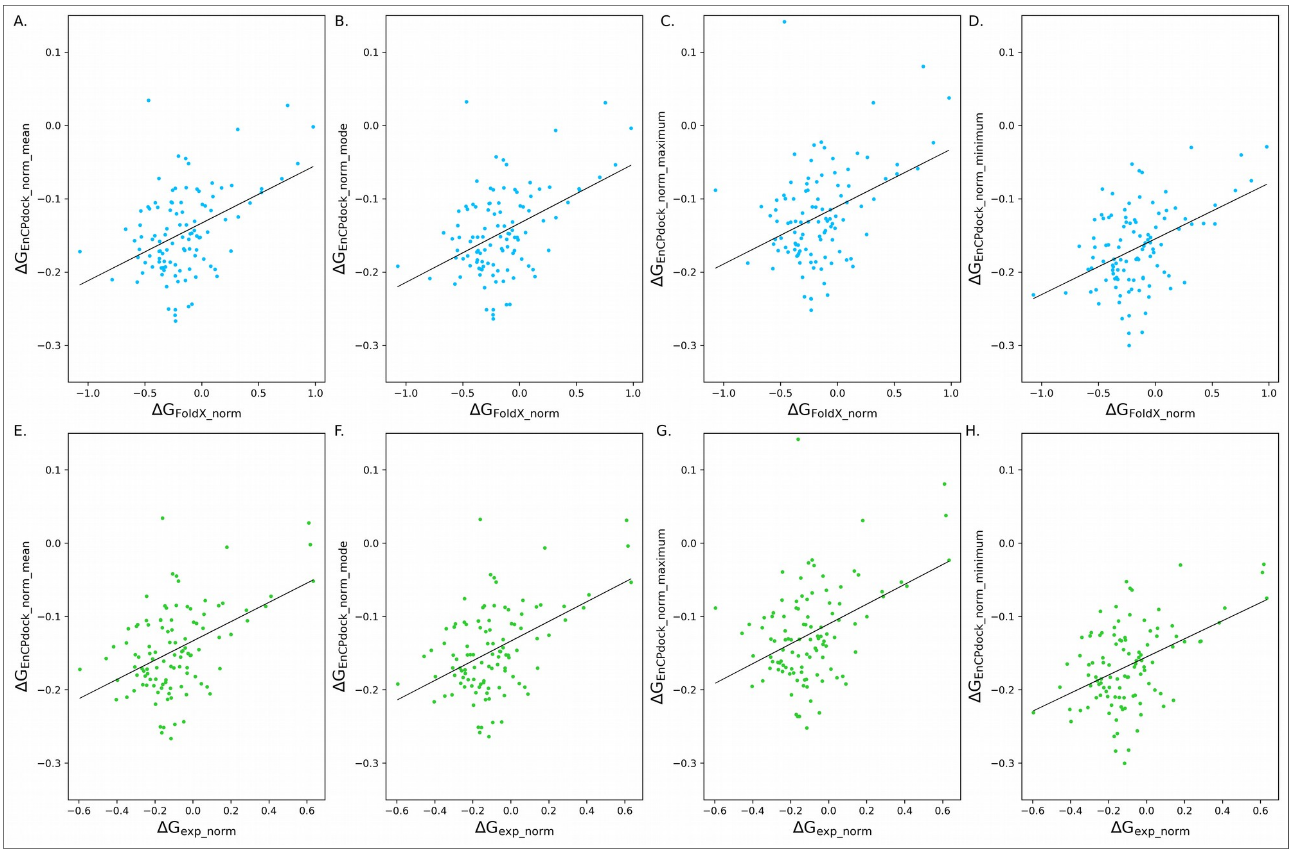

**Table S2. Correlations attained in independent validations.** Pearson's Correlation coefficients (r) for different descriptive statistical measures (central tendencies: mean, median, mode; extreme values: minimum, maximum) of normalized binding free energies predicted by EnCPdock ($\Delta G_{EnCPdock\_norm}$) and their corresponding FoldX-derived ($\Delta G_{FoldX\_norm}$) and experimental values ($\Delta G_{exp\_norm}$) attained in independent validation.

| Normalized $\Delta G_{binding}$ terms | $\Delta G_{FoldX\_norm}$ | $\Delta G_{exp\_norm}$ |
|---|---|---|
| $\Delta G_{EnCPdock\_norm-mean}$ | 0.44 | 0.46 |
| $\Delta G_{EnCPdock\_norm-median}$ | 0.45 | 0.48 |
| $\Delta G_{EnCPdock\_norm-mode}$ | 0.45 | 0.47 |
| $\Delta G_{EnCPdock\_norm-maximum}$ | 0.39 | 0.43 |
| $\Delta G_{EnCPdock\_norm-minimum}$ | 0.44 | 0.46 |

**Table S3. Performance of EnCPdock in independent validation – 1.** Performance is estimated in the Affinity benchmark dataset (1st independent validation, 106 targets) from confusion matrices subsequent to converting the continuous predicted output ($\Delta G_{EnCPdock\_norm}$) to 1-0 binaries based on different cut-offs spanning the attained ranges of $\Delta G_{FoldX\_norm}$ and $\Delta G_{exp\_norm}$ (sampled at a regular interval of 0.1) in the respective performance evaluations against the two parameters ($\Delta G_{FoldX\_norm}$, $\Delta G_{exp\_norm}$) taken as independent benchmarks. The performance is then measured based by the TPR, TNR and BACC values (see section **2.2.3**, **Materials and Methods**) for each cut-off.

| | **cut-offs** | **$\Delta G_{EnCPdock\_norm}$** | | |
|---|---|---|---|---|
| | | **TPR** | **TNR** | **BACC** |
| **$\Delta G_{FoldX\_norm}$** | -0.2 | 0.084 | 0.994 | 0.539 |
| | -0.1 | 0.366 | 0.969 | 0.667 |
| | 0.0 | 0.547 | 0.944 | 0.746 |
| | 0.1 | 0.657 | 0.929 | 0.793 |
| | 0.2 | 0.726 | 0.916 | 0.821 |
| | 0.3 | 0.772 | 0.907 | 0.84 |
| | 0.4 | 0.806 | 0.900 | 0.853 |
| | 0.5 | 0.832 | 0.895 | 0.863 |
| | 0.6 | 0.851 | 0.891 | 0.871 |
| | 0.7 | 0.867 | 0.887 | 0.877 |
| | 0.8 | 0.88 | 0.885 | 0.882 |
| | 0.9 | 0.891 | 0.884 | 0.887 |
| | 1.0 | 0.9 | 0.884 | **0.892** |
| **$\Delta G_{exp\_norm}$** | -0.2 | 0.065 | 0.973 | 0.519 |
| | -0.1 | 0.551 | 0.921 | 0.736 |
| | 0.0 | 0.742 | 0.884 | 0.813 |
| | 0.1 | 0.827 | 0.864 | 0.845 |
| | 0.2 | 0.871 | 0.852 | 0.862 |
| | 0.3 | 0.898 | 0.844 | 0.871 |
| | 0.4 | 0.916 | 0.838 | 0.877 |
| | 0.5 | 0.929 | 0.834 | 0.881 |
| | 0.6 | 0.938 | 0.830 | **0.884** |

**Table S4. Performance of EnCPdock in independent validation – 2.** Performance is estimated in the 'Proximate+SKEMPI – merged dataset' (2nd independent validation, 236 targets) from confusion matrices subsequent to converting the continuous predicted output ($\Delta G_{EnCPdock\_norm}$) to 1-0 binaries based on different cut-offs spanning the attained ranges of $\Delta G_{FoldX\_norm}$ and $\Delta G_{exp\_norm}$ (sampled at a regular interval of 0.1) in the respective performance evaluations against the two parameters ($\Delta G_{FoldX\_norm}$, $\Delta G_{exp\_norm}$) taken as independent benchmarks. The performance is then measured based by the TPR, TNR and BACC values (see section **2.2.3**, **Materials and Methods**) for each cut-off.

| | **Cut-off** | **$\Delta G_{EnCpdock\_norm}$** | | |
|---|---|---|---|---|
| | | **TPR** | **TNR** | **BACC** |
| **$\Delta G_{exp\_norm}$** | -0.7 | 1.000 | 0.915 | 0.957 |
| | -0.6 | 0.600 | 0.806 | 0.703 |
| | -0.5 | 0.633 | 0.719 | 0.676 |
| | -0.4 | 0.753 | 0.724 | 0.738 |
| | -0.3 | 0.872 | 0.778 | 0.825 |
| | -0.2 | 0.987 | 0.857 | **0.922** |
| **$\Delta G_{FoldX\_norm}$** | -0.7 | 1.000 | 0.762 | 0.881 |
| | -0.6 | 0.467 | 0.641 | 0.554 |
| | -0.5 | 0.667 | 0.486 | 0.576 |
| | -0.4 | 0.854 | 0.414 | 0.634 |
| | -0.3 | 0.945 | 0.444 | 0.695 |
| | -0.2 | 0.983 | 0.571 | **0.777** |
